# Characterisation and Genetic Architecture of Major Depressive Disorder Subgroups Defined by Weight and Sleep Changes

**DOI:** 10.1101/2022.08.30.504498

**Authors:** Sally Marshall, Mark J Adams, Kathryn L Evans, Rona J Strawbridge, Archie Campbell, Caroline Hayward, Andrew McIntosh, Pippa Thomson

**Affiliations:** Centre for Genomic and Experimental Medicine, Institute of Genetics and Cancer, University of Edinburgh; Centre for Clinical Brain Sciences (Division of Psychiatry), University of Edinburgh; School of Health and Wellbeing, College of Medical, Veterinary and Life Sciences, University of Glasgow; Department of Medicine Solna, Karolinska Institutet; MRC Human Genetics Unit, Institute of Genetics and Cancer, University of Edinburgh

## Abstract

Major depressive disorder, MDD, is highly heterogeneous and thus subgroups with different underlying aetiologies have been postulated. The aim of this work is to further characterise depression subgroups defined using sleep and weight changes. Probable lifetime MDD cases (n = 26,662) from the UK Biobank were stratified into three subgroups defined by self-reported weight and sleep changes during worst episode: (i) increased weight and sleep (↑WS), (ii) decreased weight and sleep (↓WS) and (iii) the remaining uncategorised individuals. Analyses compared the depression characteristics, mental health scores, neurological and inflammatory comorbidities and genetic architecture between subgroups and with 50,147 controls from UK Biobank. In contrast to ↑WS depression, ↓WS depression had a higher age of onset and lower proportion reporting countless or continuous episodes compared to uncategorised individuals. The ↓WS depression also had a higher wellbeing score than the other subgroups. Analyses of subgroup comorbidities identified a novel association between ↑WS depression and epilepsy. Subgroup-specific GWAS identified three genome-wide significant loci associated with ↑WS in genes previously associated with immunometabolic traits and response to anticonvulsants. The effect of BMI adjustment in the genetic analyses of the subgroups and using broader weight-only definitions were also examined. The findings provide further evidence for differences in the characteristics and genetic architecture of depression subgroups defined by sleep and weight change and highlight the importance of dividing non-↑WS individuals into ↓WS and uncategorised subgroups in analyses, as ↓WS symptoms may identify a more acute depression subgroup.

## Introduction

Major depressive disorder (MDD) is a polygenic and highly prevalent psychiatric disorder. In 2017, the World Health Organisation (WHO) estimated that there are around 322 million people living with depression globally, a figure that has grown 18.4% since 2005(1). Depression has a complex underlying genetic architecture, evidenced by the 243 risk loci identified in a recent meta-analysis of >1.3 million individuals(2). MDD is also highly heterogeneous in presentation. For instance, in a study using the criteria in the Diagnostic and Statistical Manual of Mental Disorders 5 (DSM-5)(3), over 1,000 unique MDD diagnostic symptom profiles were observed in a clinical setting(4). The variation in symptoms, severity and remission observed between individuals has resulted in the hypothesis that MDD includes multiple subgroups, with potentially different underlying aetiologies(5).

One proposed subgroup is MDD with atypical features, or atypical depression, which is characterised in the DSM-5 as MDD with any two of the following symptoms: mood reactivity, interpersonal rejection sensitivity, “leaden paralysis”, weight gain, increased appetite, or hypersomnia(3). In particular, reversed neurovegetative symptoms (weight gain, increased appetite, and hypersomnia) have been observed to be highly specific to atypical depression(6). In contrast, weight loss, hypophagia and insomnia are frequently observed in ‘typical’ MDD patients. Thus, it is becoming increasingly common for both genetic(7–9) and epidemiological studies(10–12) to utilise these symptoms over the full DSM-5 criteria when examining atypical depression. Epidemiological studies show that individuals with atypical depression are more likely to be female, have an earlier age of onset, and have more frequent and more severe depressive episodes than individuals with non-atypical depression(13). Findings that have been replicated in the UK Biobank(12). Lower genetic correlations have been observed between atypical depression and both non-atypical depression (rg = 0.59)(14) and typical depression (rg = 0.54)(9), suggestive of differences in genetic architecture.

Atypical depression has also been associated with a number of inflammatory and metabolic traits and may be an immunometabolic subgroup of depression(13, 15). Genetic studies consistently show associations between atypical depression and polygenic risk scores (PRS) for body mass index (BMI)(7–9), triglycerides(7), leptin(8), C-reactive protein and daily alcohol use(9). In the UK Biobank, individuals with atypical depression were shown have higher rates of obesity, cardiovascular disease and metabolic syndrome (hypertension, diabetes, hypercholesterolemia) than those in a non-atypical depression group(12). However, less is known about neurological and inflammatory disease prevalence within subgroups. Differences in associations with atypical and melancholic depression have been observed in individuals with migraine and in migraine subgroups(16) and it has also been reported that MDD in individuals with multiple sclerosis was characterised by a higher risk of the atypical symptoms of weight gain and leaden paralysis(17). However, the number of individuals with MDD in these analyses were relatively small; 1,297 lifetime DSM-4 MDD individuals(16) and 369 past-12 month DSM-5 MDD individuals(17), respectively.

For the most part, studies investigating subgroups of depression defined using the neurovegetative symptoms of weight and sleep change in the UK Biobank(9, 12, 14) have compared the atypical subgroup of depression to either a non-atypical subgroup or to typical depression (defined as depression with decreased weight and sleep). As a result, the typical subgroup and individuals not categorised as either atypical or typical are not as well characterised, despite representing approximately 34% and 60% of MDD cases, respectively. Furthermore, the only published genome-wide association studies (GWAS) of neurovegetative subgroups of depression in the UK Biobank are case-control analyses of atypical and non-atypical MDD(14), which identified no loci associated with atypical depression and six loci associated with non-atypical depression (*p* < 5 × 10^-8^; *NEGR1*, *RPSAP2*, *OAS1*, *OLFM4, SUPT4H1*, *KDM4B*). Potential differences in genetic aetiology between depression subgroups may be better captured in case-only GWAS. In addition, there has been inconsistency in whether genetic analyses of neurovegetative subgroups are adjusted for BMI. Some have performed adjustment for BMI on the basis that a significant difference in BMI is observed between subgroups(9). Others argue that as there is evidence for a shared genetic basis between atypical depression and BMI, any adjustment may be an overcorrection(7) or that adjusting genetic analyses for heritable traits can introduce bias, if the relationship between the covariate and outcome if not well understood(18). Whilst GWAS of MDD overall do not correct for BMI (19–22).

The aim of this research was to build on previous work characterising neurovegetative depression subgroups in the UK Biobank(9, 12, 14). Firstly, by expanding the analyses to three comparator groups in order to investigate previously unexplored differences between decreased weight and sleep and uncategorised individuals. Secondly, by comparing the prevalence of neurological and inflammatory disease between subgroups. Lastly, by performing both case-control and case-only GWAS of subgroups, polygenic risk score analyses in an independent sample and examining the effect of BMI-adjustment in genetic analyses. Sensitivity analyses were also performed in subgroups defined by weight change only, to examine whether this broader definition represents a similar subgroup to those defined by both weight and sleep change.

## Materials and methods

### Participants

For all analyses, except the polygenic risk score (PRS) analyses, participants were drawn from the UK Biobank(23). As PRS analyses require individuals from an independent sample, participants were drawn from Generation Scotland(24, 25). Full participant, genotyping and phenotyping information for both cohorts can be found in the **supplementary methods**. In both cohorts the probable lifetime MDD cases (UK Biobank: n = 26,662, Generation Scotland: n = 1,506) and subsequent weight-sleep subgroup classifications were defined using participant responses to questionnaires based on the Composite International Diagnostic Interview Short Form (CIDI-SF). Controls were defined as individuals without a history of depressed mood, anhedonia or other mental illnesses (UK Biobank: n = 50,147, Generation Scotland: n = 7,667). In the UK Biobank there were 1,525 individuals reporting increased weight and sleeping more (↑WS) during their worst depressive episode, representing atypical depression, and 9,067 individuals reporting decreased weight and sleeping less (↓WS) during their worst episode, representing typical depression. The remaining individuals (n = 16,070) were included in analyses as a third subgroup, hereby referred to uncategorised depression.

## Depression Characteristics and Mental Health Scores

Sample (age, sex and BMI at initial assessment) and depression characteristics (age of onset, number of episodes and reporting countless or continuous episodes in the MHQ) were compared between subgroups. In addition four questionnaire-based scores relating to current mental health were also examined: neuroticism (26), patient health questionnaire 9 (PHQ-9)(27), generalised anxiety disorder 7 (GAD-7)(28) and wellbeing(29). Derivation of the neuroticism score (UKB field 20127) for UK Biobank participants is described in Smith *et al*. (26). The PHQ-9, GAD-7, and wellbeing scores were calculated utilising participants responses to the UK Biobank mental health questionnaire (MHQ) questions as described by Davis *et al*.(29). A higher neuroticism, PHQ-9 and GAD-7 score represented increased severity. The wellbeing score was coded such that a higher score represented better wellbeing. Not all individuals had measures for all scores, samples sizes for each score can be found in **Supplementary Table 1**. For numerical variables, where data were normally distributed, a t-test was used for comparisons between subgroups, otherwise a Wilcoxon test was used. For categorical variables, comparisons were performed using a Chi-square test of independence. For sample and depression characteristics, a result was considered statistically significant if the *p*-value for the comparison was less than a Bonferroni corrected threshold of 5.56 x 10^-3^ (0.05/9, 3 related traits, 3 depression subgroups). For mental health scores the Bonferroni corrected threshold for significance used was 4.17 x 10^-3^ (0.05/12, 4 related traits, 3 depression subgroups).

## Neurological and Inflammatory Disease Prevalence

Disease cases of epilepsy, migraine, multiple sclerosis, dementia, Crohn’s disease, ulcerative colitis, rheumatoid arthritis, asthma and eczema were identified using ICD10 codes in electronic health records or non-cancer illness self-reported to a trained nurse at in person assessment (UKB field 20002). Codes used to identify cases are shown in **Supplementary Table 3**. As dementia is an age-related disorder, parental dementia (UKB fields 20107 and 20110, disease code = 10) was used for analysis to reduce the effect of sample age on results. A Chi-square test of independence was utilised to compare disease counts between depression subgroups and between subgroups and controls, unless the expected values of greater than 20% of the cells of any given contingency table were less than 5(30). In which case a Fisher’s exact test was used to calculate *p*-values. A result was considered statistically significant if the *p*-value for the comparison was less than a Bonferroni corrected threshold of 8.33 x 10^-3^ for six comparisons (0.05/6, for each disorder 3 subgroup-control comparisons and 3 subgroup-subgroup comparisons).

### Genetic Analyses

All genetic analyses were adjusted for technical covariates (array, batch and centre; UKB field 54 & UKB field 22000) as well as 40 genetic ancestry principal components (UKB field 22009), age, sex and BMI (UKB field 21001). Analyses were then repeated without adjustment for BMI. There were some individuals for whom BMI was not available. Sample sizes for all analyses can be found in **Supplementary Table 1**. Adjustment was performed by regressing the phenotype on covariates prior to analyses, then using the residuals from each regression as the phenotype.

### Genome-wide Association Studies (GWAS)

For each depression subgroup both case-control and case-only GWAS were performed. Case-only GWAS compared ↑WS to ↓WS, ↑WS to uncategorised depression and ↓WS to uncategorised depression. All GWAS were conducted using Plink 1.9(31, 32), with a genome-wide significant threshold of *p* < 5 × 10^-8^ and suggestive significance threshold of *p* < 1 × 10^-5^. Post-GWAS analyses were performed in FUMA(33). All SNPs of suggestive significance (*p* < 1 × 10^-5^) and independent of one another at r^2^ < 0.6 were identified, independent lead SNPs were then defined as those independent from one another at r^2^ < 0.1. Post-hoc power calculations for case-control and case-only GWAS were performed using the R package ‘genpwr’(34), results can be found in the **Supplementary Figures 6 & 7**. Genome-wide significance for gene-based tests (performed using MAGMA(35) as part of post-GWAS analyses) was defined as *p* < 3 × 10^-6^ after Bonferroni correction for multiple testing, as there were 16,686 protein coded genes included in the analysis.

### Polygenic Risk Score Analyses

Polygenic risk score (PRS) analyses of case-control and case-only GWAS results from UK Biobank were performed using PRSice(36) in the independent Generation Scotland cohort. Only unrelated individuals were included in analyses, filtering of related individuals was performed in a manner that maximised the sample of the smallest group in each comparison. Samples sizes for each comparison are shown in the **Supplementary Table 2**. PRS analyses were adjusted for age, sex and 20 ancestry principal components. For each GWAS, PRS were calculated using a number of *p*-value thresholds (5 × 10^-8^, 1 × 10^-5^, 1 × 10^-4^, 5 × 10^-4^, 0.001, 0.005, 0.01, 0.1, 0.5, 1), the reported ‘best’ *p*-value threshold was considered to be the one with the most significant association with the corresponding Generation Scotland sample. The best *p*-values thresholds and number of SNPs in each PRS are shown in **Supplementary Tables 27-30**. A result was considered statistically significant if the *p*-value for the comparison was less than a Bonferroni corrected threshold of 4.17 × 10^-3^ for twelve comparisons (0.05/12).

### Genetic Correlation Analyses

Genetic correlations between subgroups and between subgroups and MDD (19) were calculated using LD score regression, implemented in ldsc, as this method is robust to sample overlap(37, 38). Genetic correlations were compared using a z-test:

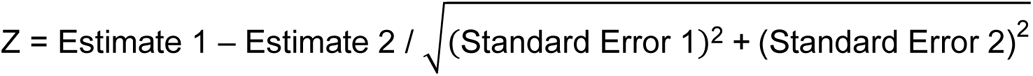

A result was considered statistically significant if the *p*-value for the comparison was less than a Bonferroni corrected threshold of 0.0167 for three comparisons (0.05/3).

### Sensitivity Analyses

All analyses were repeated in more broadly defined depression subgroups using weight change during their worst episode only. In these analyses, atypical depression was represented by a weight gain subgroup (↑W, n = 5,015), typical depression by a weight loss subgroup (↓W, n = 10,924), and there were 10,723 individuals that were not categorised as weight gain or weight loss in the uncategorised subgroup (**Supplementary Figure 1**). Controls were defined as in the main analyses.

## Results

### Sample and Depression Subgroup Characteristics

The sample and depression characteristics of each depression subgroup are shown in **Table 1** (**Supplementary Tables 6-8**). Consistent with what has previously been reported comparing ↑WS to a non-↑WS subgroup(12), the ↑WS subgroup had a significantly higher BMI at initial assessment, earlier age of onset of depression and higher number of depressive episodes compared with both the other subgroups (*p* < 0.001). However, dividing the non-↑WS subgroup into ↓WS and uncategorised individuals revealed that there was no significant difference in the proportion of females between the ↑WS and ↓WS subgroups (*p* = 0.63), but the proportion of females was significantly higher in both the ↑WS and ↓WS subgroups compared to uncategorised individuals (*p* < 0.001). Furthermore, compared to uncategorised individuals, the ↓WS subgroup had a significantly lower BMI at initial assessment, later age of onset and lower proportion of individuals reporting “Too many [episodes] to count/One episode ran into the next” (*p* < 0.001). Overall, these results suggest there are differences in depression characteristics between the ↓WS subgroup and uncategorised individuals not captured by previous studies.

**Table 1.** Sample and Depression Characteristics of Weight-Sleep Depression Subgroups.

|  | ↑WS<br>Subgroup | ↓WS<br>Subgroup | Uncategorised<br>Subgroup |
| --- | --- | --- | --- |
| **Sample Characteristics** |  |  |  |
| Age at Initial Assessment (years) | 52.38<br>(7.10) | 55.11<sup>†</sup><br>(7.45) | 54.95<sup>†</sup><br>(7.55) |
| Proportion of<br>Females (%) | 77.11<sup>‡</sup> | 76.52<sup>‡</sup> | 65.02 |
| Body Mass Index* (kg/m<sup>2</sup>) | 29.72<br>(26.36-33.64) | 25.18<br>(22.88-28.18) | 26.61<br>(23.98-30.01) |
| **Depression Characteristics** |  |  |  |
| Age of Onset (years) | 32.42<br>(14.58) | 36.38<br>(14.07) | 35.32<br>(14.58) |
| Number of Episodes* | 3<br>(1-4) | 1<br>(1-3) | 1<br>(1-3) |
| Proportion of Individuals reporting: "Too<br>many to count/one episode ran into the<br>next" (%) | 37.44% | 13.83% | 21.59% |

*For characteristics denoted by \* median and interquartile range (IQR) are shown, rather than mean and standard deviation, as data were not normally distributed. All the comparisons were significant ( $p < 5.56 \times 10^{-3}$ ) except those denoted by matching symbols (<sup>†</sup> $p = 0.10$ , <sup>‡</sup> $p = 0.63$ , <sup>||</sup> $p = 0.18$ ).*

### Mental Health Scores

Scores of current mental health were compared between all three subgroups (**Figure 1**, **Supplementary Table 8**). The ↑WS subgroup had significantly higher scores for neuroticism, PHQ-9 and GAD-7 (**Figures 1a, 1b** & **1c**) and lower wellbeing scores (**Figure 1d**) compared to the other subgroups (*p* < 0.001). In contrast, the ↓WS subgroup had a significantly higher wellbeing score (**Figure 1d**) and lower neuroticism, PHQ-9 and GAD-7 scores (**Figures 1a, 1b** & **1c**) than the other subgroups. In addition, individual PHQ-9 symptoms were reported ‘more than half the days’ or ‘nearly every day’ for the last two weeks by a significantly larger proportion of the ↑WS subgroup and by a significantly smaller proportion of the ↓WS subgroup compared to other subgroups (**Figure 1e**, **Supplementary Table 9**, *p* < 0.001). These results suggest that individuals in the ↑WS subgroup have poorer mental health and individuals in the ↓WS better mental health at the time of completing the mental health questionnaire.

**Figure 1.**
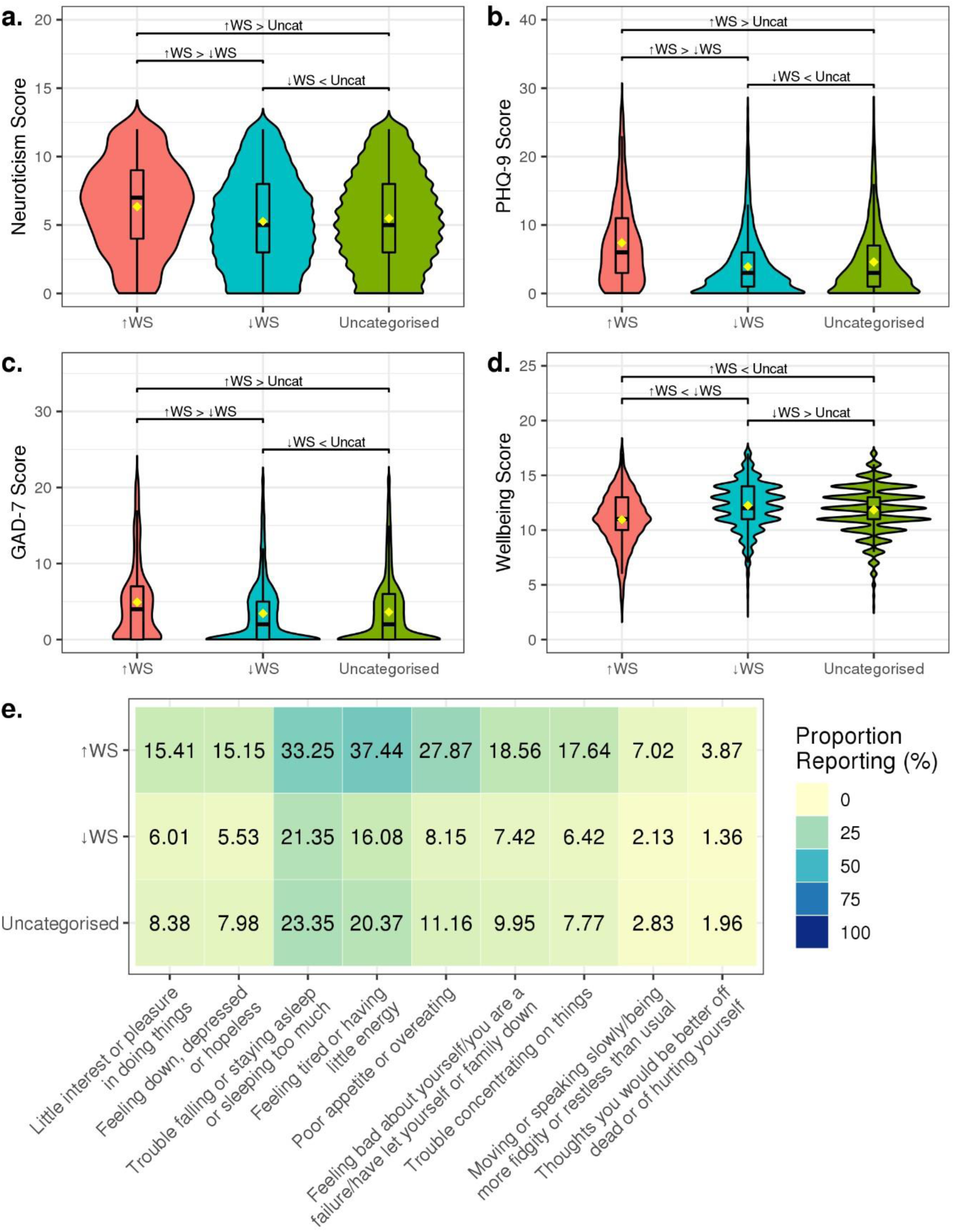
The mental health scores: (a.) neuroticism, (b.) PHQ-9, (c.) GAD-7, and (d.) wellbeing of each subgroup. The distribution of data is shown as a boxplot of the median and interquartile range, whiskers for each boxplot are calculated using the formula: quartile + 1.5*IQR and outliers are not shown. All comparisons were significant at p < 4.17 x 10^-3^. The directions of effect are noted (> greater than, < less than) (e.) Heatmap of proportion reporting experiencing each PHQ-9 depressive symptom ‘More than half the days’ or ‘Nearly every day’ over the last two weeks for each subgroup. All comparisons were significant at p < 7.14 x 10^-3^ (0.05/24).

In previous analyses in the UK Biobank, ↑WS depression has been associated with higher rates of obesity, cardiovascular disease and metabolic syndrome(12). As the wellbeing score contains a measure for happiness with health, it is possible that this measure alone drives the lower wellbeing scores observed in ↑WS depression. However, investigation of the individual measures contained within the wellbeing score (“General happiness”, “Happiness with health”, and “Life thought to be meaningful”) showed that, compared to the uncategorised individuals, the ↑WS subgroup had a significantly lower score and the ↓WS subgroup a significantly higher score in each individual measure (**Supplementary Table 5**, *p* < 0.001).

### Neurological and Inflammatory Co-morbidities

Potential differences in neurological and inflammatory disease prevalence between the three subgroups were investigated (**Figure 2**, **Supplementary Tables 14 & 15**). Overall, there were significantly increased odds of co-morbid epilepsy, migraine, rheumatoid arthritis, asthma and eczema in lifetime MDD cases compared to controls. For epilepsy, migraine and asthma, significantly higher odds of comorbidity were observed within the ↑WS subgroup than in lifetime MDD or the other subgroups (*p* < 3.60 x 10^-3^). This pattern is strongest in epilepsy where the majority of the co-morbidity seen in lifetime MDD may be explained by specific comorbidity in the ↑WS subgroup (**Figure 2a**). A nominal increase in the odds of comorbid parental dementia was observed in the uncategorised individuals (*p* < 0.024), despite there being no significant difference in comorbidity in lifetime MDD. Suggesting that, for certain comorbidities, differences between the subgroups may be hidden when only examining lifetime MDD.

**Figure 2.**
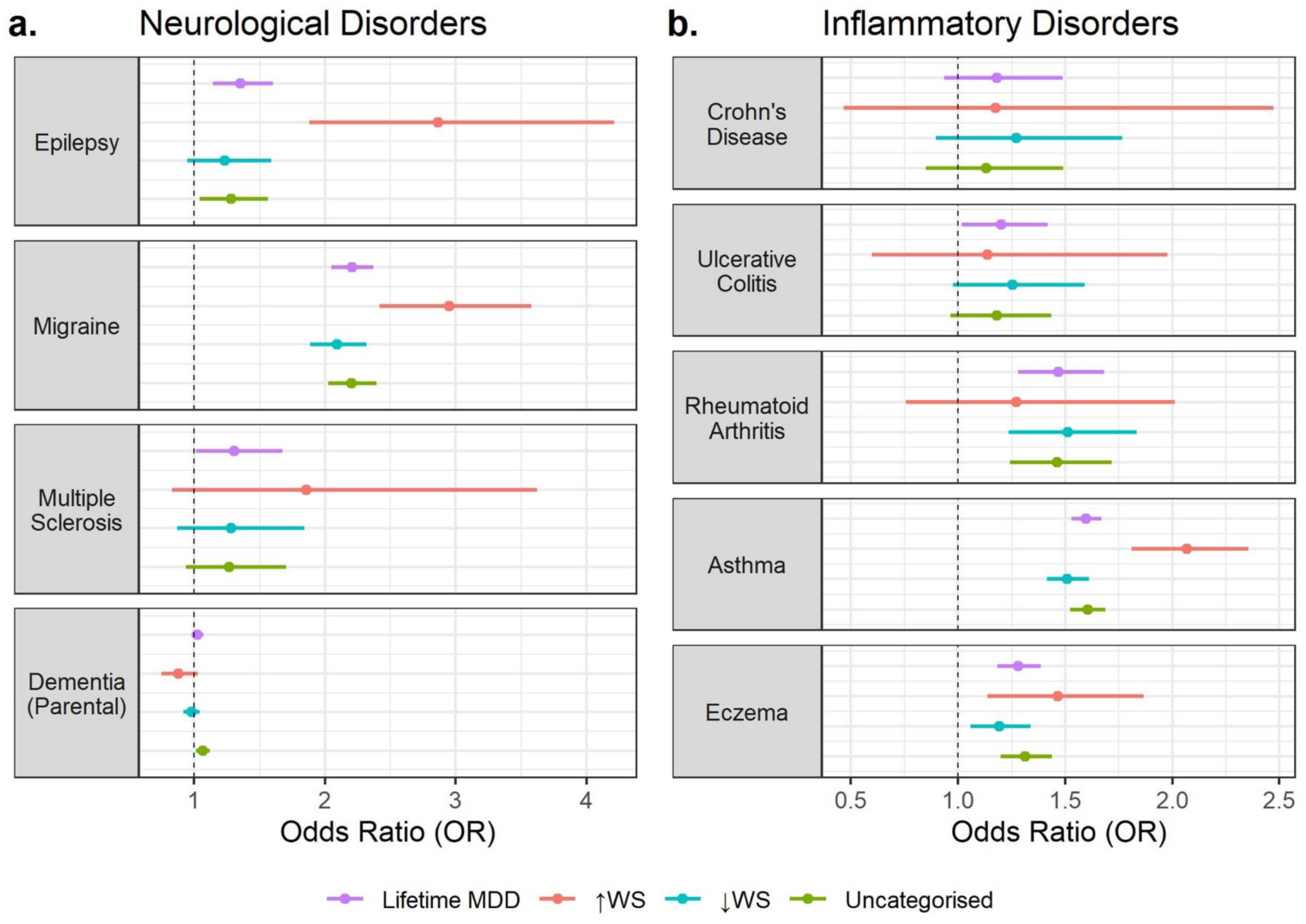
The odds ratios and 95% confidence intervals of each depression subgroup compared to controls in (a.) four neurological and (b.) five inflammatory disorders of interest.

### Genome-wide Association Studies

Case-control and case-only GWAS were performed to investigate the genetic architecture of depression subgroups. To examine the effect of BMI adjustment on results, all GWAS were performed adjusted and unadjusted for BMI. All SNPs below ‘suggestive’ significance (*p* < 1 x 10^-5^) in each GWAS are shown in **Supplementary Tables 18-21**. Across all GWAS performed, only three loci of genome-wide significance (*p* < 5 × 10^-8^) were identified, all in the ↑WS case-control analyses adjusted for BMI. These were rs7608034 (*ALK*, β = 1.14, 95% CI = 0.73-1.55, *p* = 3.09 × 10^-8^, **Supplementary Figure 8**), rs1554675 (*EPHB1*, β = 0.92, 95% CI = 0.63-1.21, *p* = 8.29 × 10^-10^, **Supplementary Figure 9**) and rs1426041 (*CLSTN2*, β = 0.68, 95% CI = 0.48-0.88, *p* = 5.12 × 10^-11^, **Supplementary Figure 10**). When analyses were unadjusted for BMI, only the *CLSTN2* locus remained above suggestive significance in ↑WS case-control analyses (β = 0.41, 95% CI = 0.25-0.57, *p* = 3.07 × 10^-7^). The *CLSTN2* SNP, rs1426041, was also associated at suggestive significance in BMI-adjusted and unadjusted case-only GWAS of ↑WS vs ↓WS (βadj = 0.42, 95% CI = 0.26-0.57, *p* = 8.33 × 10^-8^; βunadj = 0.29, 95% CI = 0.17-0.41, *p* = 8.04 × 10^-6^) and ↑WS vs uncategorised individuals (βadj = 0.34, 95% CI = 0.21-0.47, *p* = 3.31 × 10^-7^; βunadj = 0.29, 95% CI = 0.17-0.41, *p* = 7.19 × 10^-6^, **Figure 3**).

**Figure 3.**
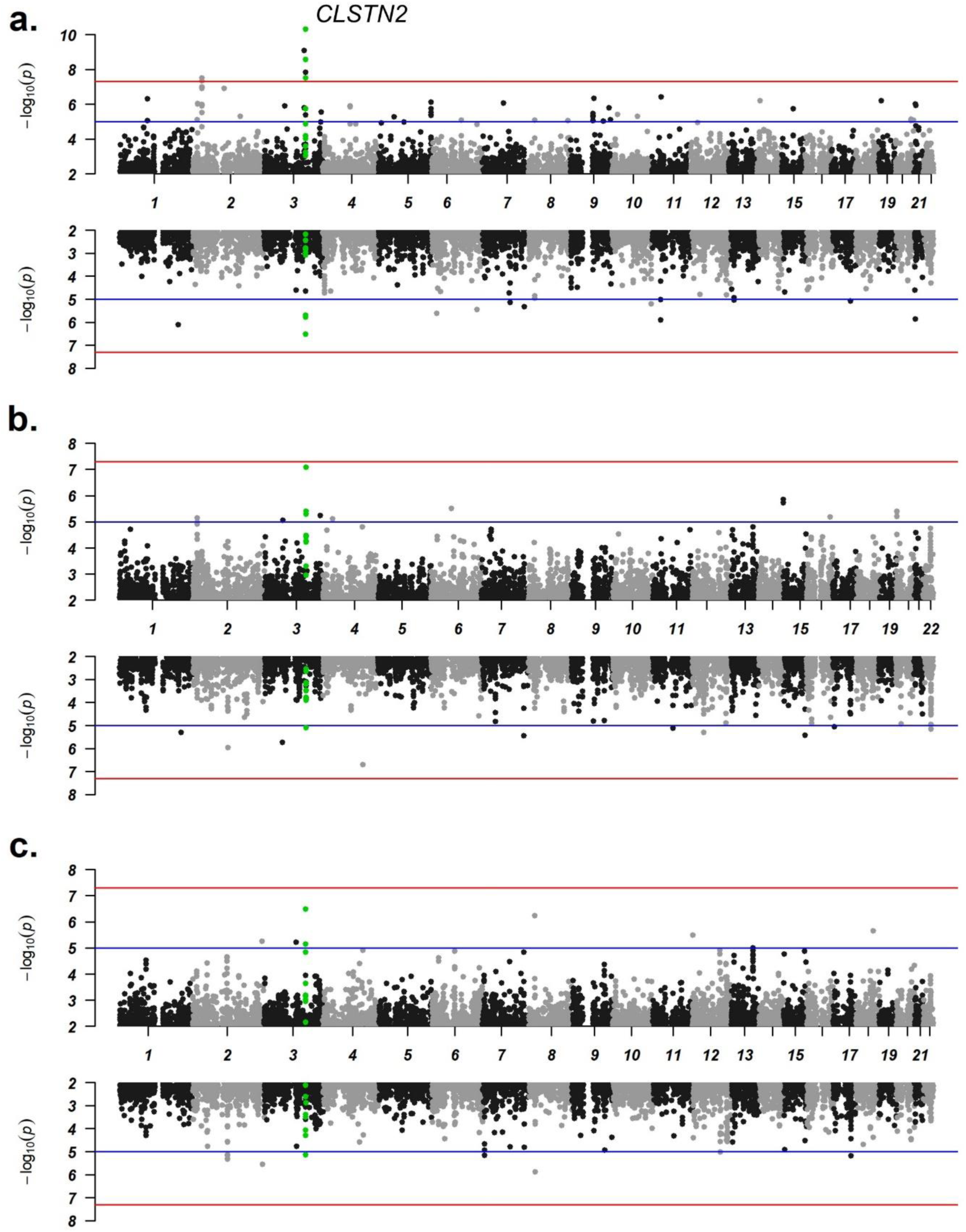
Miami plots of the (a.) ↑WS case-control analyses (b.) ↑WS vs ↓WS case-only analyses and (c.) ↑WS vs uncategorised case-only analyses. Analyses adjusted for BMI are shown in the top plot and analyses unadjusted for BMI are shown in the bottom plot. Chromosomal location is on the x-axis and −log10 of the p-value for each SNP is on the y-axis. The red line indicates the threshold for genome-wide significance (5.00 × 10^-8^) and the blue dashed line indicates the threshold for “suggestive” significance (1.00 × 10^-5^). Loci within the CLSTN2 gene (chr3: 139,654,027-140,296,239) are highlighted in green.

MAGMA gene-based tests (**Supplementary Tables 23-24**) identified three genes significantly associated (*p* < 3 × 10^-6^) with the subgroups in case-control analyses adjusted for BMI. *RGS14* was associated with the ↑WS subgroup (*p* = 3.76 × 10^-7^), *AKNAD1* was associated with the ↓WS subgroup (*p* = 1.14 × 10^-6^) and *ZNF365* was associated with the uncategorised subgroup (*p* = 1.15 × 10^-6^). *AKNAD1* (*p* = 1.14 × 10*^-6^*) and *ZNF365* (*p* = 1.33 × 10^-6^) remained significantly associated when analyses were not adjusted for BMI.

### Polygenic Risk Score Analyses

The results from the UK Biobank GWAS were used to calculate the subgroup-specific PRS and tested for associations with weight-sleep subgroups in the independent Generation Scotland cohort (**Figure 4**, **Supplementary Tables 27-30**). Regardless of whether GWAS were adjusted or unadjusted for BMI, case-control PRS for the ↓WS and uncategorised subgroups and ↓WS vs uncategorised case-only PRS were significantly associated with their corresponding subgroups after correction for multiple testing. The ↑WS vs ↓WS case-only PRS was nominally associated in a sample of ↑WS and control individuals, but only when using the GWAS adjusted for BMI. Lastly, the ↑WS case-control PRS was negatively associated with the ↑WS and controls, ↑WS and ↓WS, and ↑WS and uncategorised samples when using the PRS unadjusted for BMI. However, this may be due to greater influence of a small number of negatively associated SNPs. A high-resolution plot of the ↑WS PRS in a ↑WS and control sample (**Supplementary Figure 22**) shows a spike in the PRS model fit at very low *p*-value thresholds. At *p*-value thresholds 0.1 or above associations were positive, although not significant. Overall, these results suggest that case-control PRS of the ↓WS and uncategorised subgroups are specific to these subgroups in an independent sample, which could indicate differences in the underlying genetic architecture of these depression subgroups.

**Figure 4.**
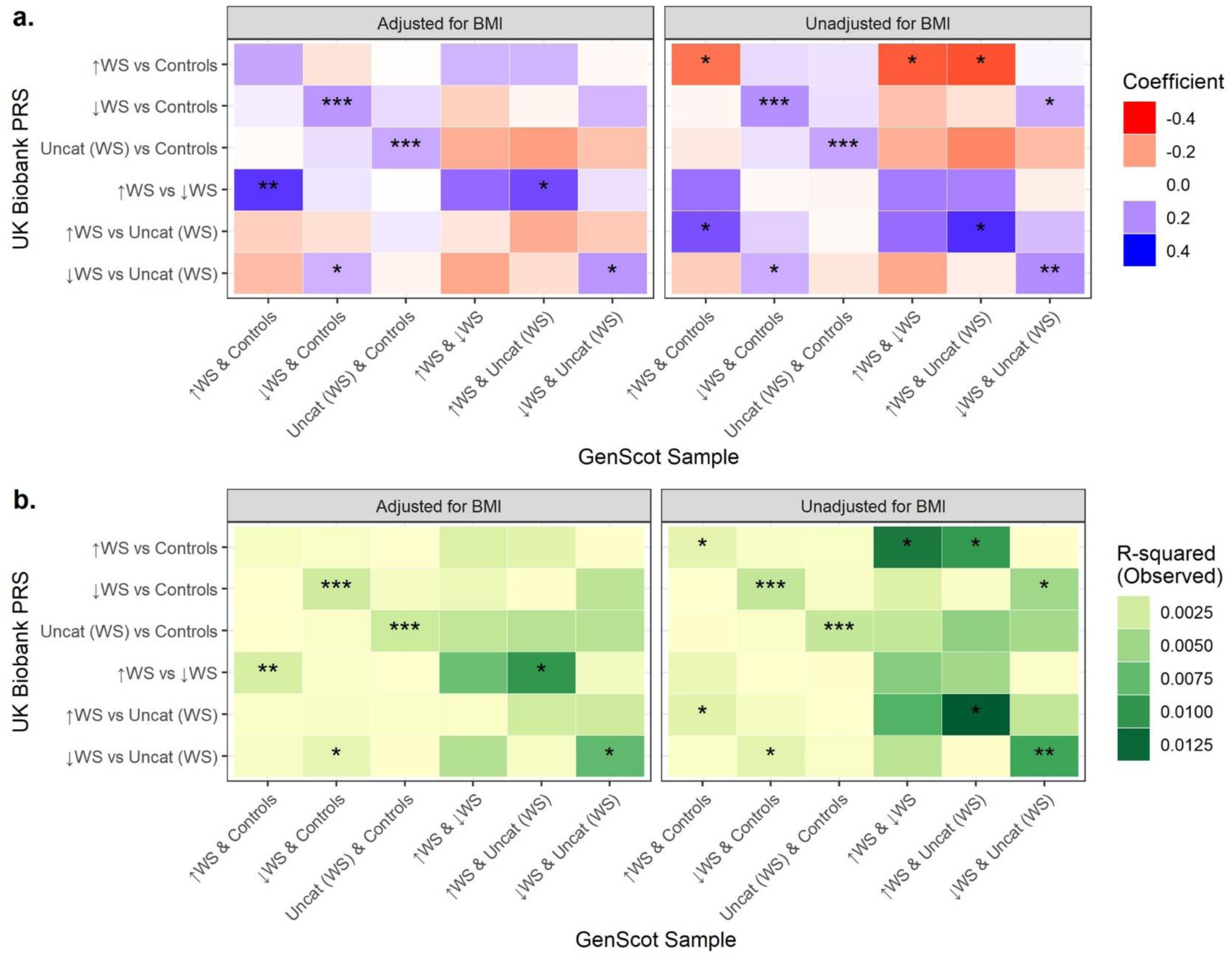
Heat maps of the results of the association analyses of PRS calculated using UK Biobank case-control and case-only GWAS summary statistics in Generation Scotland case-control and case-only samples. (a.) Heat map of the coefficient values from each analysis both using adjusted and unadjusted GWAS summary statistics to calculate PRS, and (b.) heat map of the r2 values from each analysis both using adjusted and unadjusted GWAS summary statistics to calculate PRS. Significance: *p > 0.05. **p > 0.05/6, ***p > 0.05/12. PRS = polygenic risk score, GenScot = Generation Scotland, Uncat (WS) = uncategorised depression (weight-sleep).

### Genetic Correlation Analyses

Genetic correlation analyses were performed between subgroups and clinically defined MDD (Wray *et al*., 2018) and between the subgroups (**Figure 5, Supplementary Table 31**) using the subgroup-specific case-control GWAS results. The genetic correlation with MDD was lowest with the ↑WS subgroup, and highest with the uncategorised individuals. However, both the ↑WS and ↓WS subgroups had genetic correlations with MDD that were significantly lower than 1 (**Figure 5a**.). Consistent with previous analyses in the UK Biobank(9), the lowest genetic correlation observed between subgroups was between ↑WS and ↓WS. However, this was only significantly lower than 1 when analyses were adjusted for BMI. The same pattern was also observed between ↑WS and uncategorised individuals (**Figure 5b**.). Overall, these results suggest that the uncategorised subgroup has the most similar architecture to clinically defined MDD and that the addition of BMI as a covariate may affect the estimates of genetic correlations involving the ↑WS subgroup but does not appear to affect estimates for the ↓WS and uncategorised subgroups.

**Figure 5.**
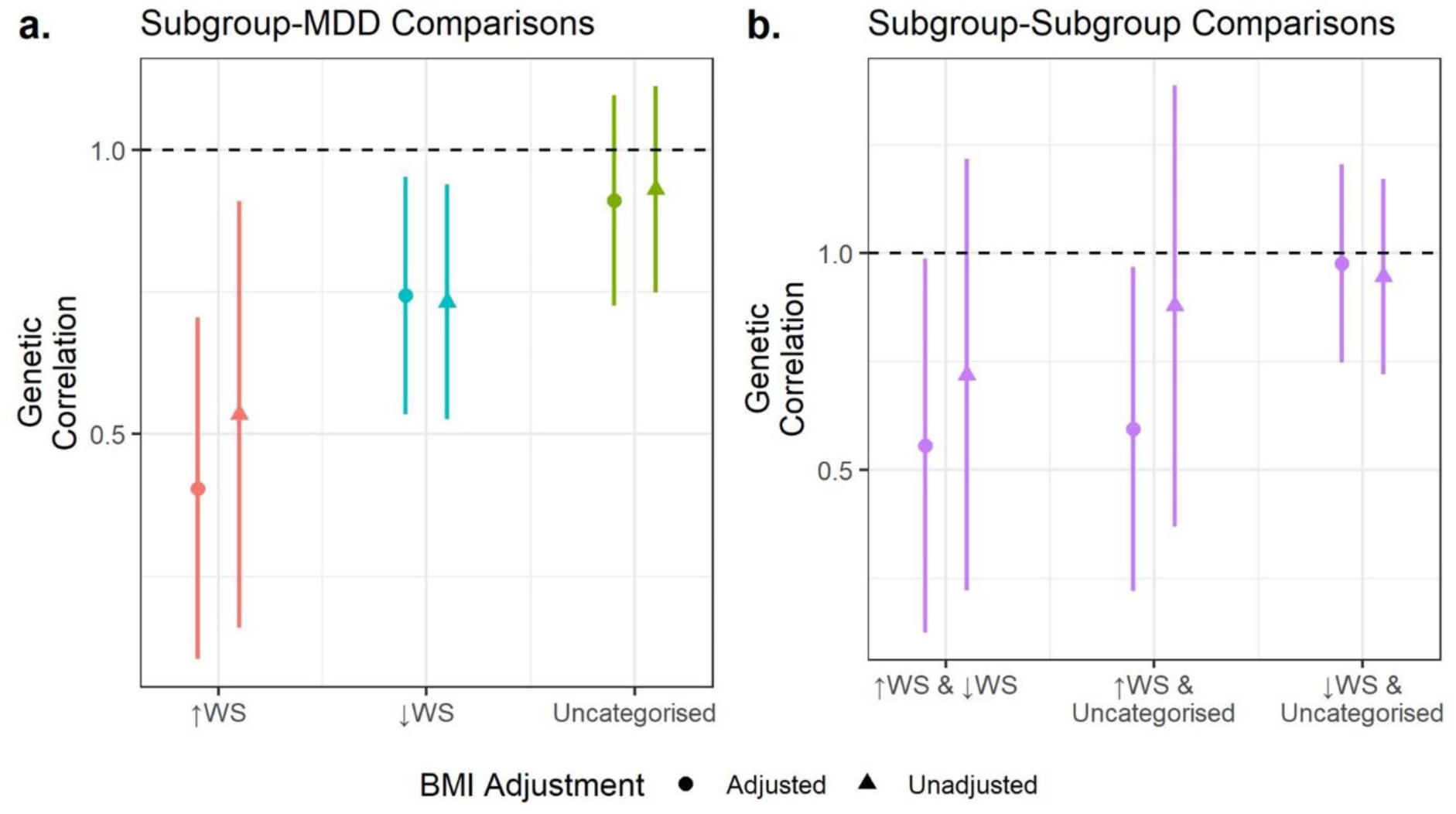
Forest plots of the genetic correlations and 95% confidence intervals of (a.) subgroup vs MDD (Wray et al., 2018) and (b.) subgroup vs subgroups. The black dashed line represents a genetic correlation of 1.00. Results adjusted (circular points) and unadjusted (triangular points) for BMI are shown.

### Sensitivity Analyses

All analyses were repeated with subgroups defined using weight change only in order to examine the characteristics of more broadly defined neurovegetative subgroups. This resulted in the addition of 3,490 and 1,857 previously uncategorised individuals into the subgroups representing atypical depression (↑W, n = 5,015) and typical depression (↓W, n = 10,924), respectively. There was a significantly higher proportion of previously uncategorised individuals in the ↑W subgroup (69.59%) than the ↓W subgroup (17.00%, Χ^2^ = 4262.1, *p* < 0.001). A total of 10,723 individuals remained uncategorised.

Similar characteristics were observed in the weight-only subgroups compared to those including sleep symptoms (**Supplementary Figures 2-5**, **Supplementary Table 4**), except that the highest proportion of females was observed in the ↓W rather than ↑W subgroup (**Supplementary Table 10**). However, there were fewer significant differences between ↑W and ↓W to uncategorised individuals. In particular, differences between ↑W and uncategorised subgroups in age of onset (**Supplementary Table 11**) and between ↓W and uncategorised subgroups in GAD-7 scores (**Supplementary Table 12**) and proportion currently reporting certain PHQ-9 symptoms (sleep disturbances, appetite changes, trouble concentrating, being noticeably slow, fidgety or restless, and thoughts of self-harm) **Supplementary Table 13**) were no longer significant after correction for multiple testing when sleep was excluded from the subgroup definition.

The case-only GWAS analyses of ↑W vs ↓W identified a further locus of suggestive significance, rs4688064 (mapped to *LSAMP*), (βadj = 0.16, *p* = 1.22 × 10^-6^, βunadj = 0.15, *p* = 5.36 × 10^-8^). Which also had nominally significant opposing effects in the case-control GWAS (↑W βadj & βunadj = 0.12, *p* < 3.00 x 10^-4^; ↓W βadj & βunadj = −0.05, *p* < 5.34 x 10^-4^), (**Supplementary Figures 13 & 14**, **Supplementary Tables 18-21**). Nominal associations with the *LSAMP* locus were observed in the ↑WS vs ↓WS case-only GWAS (βadj = 0.19, βunadj = 0.18, *p* < 1.17 x 10^-3^); and the ↓WS case-control GWAS (βadj = −0.06, βunadj = - 0.06, *p* < 5.23 x 10^-3^, **Supplementary Figure 19, Supplementary Tables 18-21**).

Only the uncategorised case-control PRS calculated using weight-only GWAS was significantly associated in a sample of uncategorised individuals and controls (r^2^ > 0.00309, *p* < 1.54 x 10^-3^, **Supplementary Figures 20 & 21**, **Supplementary Tables 27-30**). The ↓WS case-control PRS was significantly associated in a weight-only subgroup sample; explaining a higher proportion of the variance in the ↓W and control sample (r^2^ > 0.00252, *p* < 3.83 x 10^-3^) than ↓W case-control PRS (r^2^ > 0.00174, *p* < 0.0156). Finally, unlike their weight-sleep counterparts, only the genetic correlation between ↓W and MDD was significantly lower than 1. In addition, there was no effect of adjusting for BMI on between subgroup genetic correlations in weight-only subgroups (**Supplementary Figures 23, Supplementary Table 31**). Overall, these results suggest that certain differences captured by the weight-sleep subgroup definitions are not well distinguished by weight-only definitions.

## Discussion

The aim of this study was to extend previous investigations of depression subgroups defined using sleep and weight changes in the UK Biobank(9, 12, 14). Phenotypic characteristics, neurological and inflammatory comorbidities and genetic architecture of three depression subgroups were investigated: (i) ↑WS, (ii) ↓WS, as well as (iii) uncategorised individuals. For the first time, differences between ↓WS and uncategorised individuals and the effect of adjusting genetic analyses of depression subgroups for BMI were examined.

It is common in analyses of atypical depression, regardless of definition, to primarily compare to a group of non-atypical individuals (12, 14). However, further dividing non-atypical individuals into ↓WS (typical) and uncategorised subgroups revealed significant differences in age of depression onset and proportion reporting “countless or continuous” depressive episodes. The ↓WS subgroup also had consistently better mental health scores than both the ↑WS and uncategorised subgroups. It is possible that the ↑WS and ↓WS subgroups identify more severe and less severe subgroups of depression. However, as the PHQ-9, GAD-7 and wellbeing score assess current mental health, another potential explanation is that the ↑WS and ↓WS symptoms discriminate between chronic and acute depression subgroups.

In addition to the previously reported cardiovascular and metabolic diseases associated with atypical depression in the UK Biobank(12), this study examined comorbidity with neurological and inflammatory disorders. There were a higher number of individuals with epilepsy, migraine and asthma in the ↑WS subgroup compared to the other depression subgroups. Indeed, the increased prevalence of epilepsy in lifetime MDD compared to controls appears to be driven almost entirely by the ↑WS subgroup. Interestingly, all three genome-wide significant loci associated with ↑WS implicated genes which had been previously associated with response or reaction to anticonvulsant medications(39–41). These anticonvulsants (carbamazepine, lamotrigine and valproate) are also prescribed as mood stabilisers in bipolar disorder(42), which is observed at higher rates in atypical compared to non-atypical depression(12, 13). A recent study of treatment resistant depression (TRD) identified an anticonvulsant (stiripentol) as a candidate compound for drug repurposing for TRD and reported a negative association between TRD and ↓WS depression and a positive association between TRD and the ↑W depression (43).

Subgroup-specific genome-wide association analyses in this study identified only three genome-wide significant loci, associated with ↑WS depression in analyses adjusted for BMI. These were rs7608034 (Anaplastic Lymphoma Receptor Tyrosine Kinase, *ALK*), rs1554675 (Ephrin Receptor B1, *EPHB1*) and rs1426041 (Calsyntenin 2, *CLSTN2*). A previous GWAS of depression with atypical features(17) did not identify any genome-wide significant loci, but utilised a broader definition of depression, a potentially less specific phenotype(24), and did not adjust for BMI. In gene-based tests, three genes were observed to be associated with the weight-sleep subgroups: *RGS14* (Regulator of G Protein Signalling 14, ↑WS unadjBMI), *AKNAD1* (AKNA Domain Containing 1, ↓WS adj and unadjBMI) and *ZNF365* (Zinc Finger Protein 365, uncategorised, adj and unadjBMI). Sensitivity analyses in weight-only subgroups identified a further locus of interest at suggestive significance: rs4688064 (Limbic System Associated Membrane Protein, *LSAMP*), in ↑W vs ↓W case-only analyses, regardless of BMI adjustment. The effects of this locus were in opposing directions in the atypical versus typical subgroups using either weight-sleep or weight-only subgroup definitions. PRS analyses also provided evidence for differences in the genetic architecture between depression subgroups. Although identifying no loci at genome-wide significance, PRS for the ↓WS and uncategorised subgroups were significantly associated with only their own subgroups in an independent sample. ↑WS PRS showed no significant association after correcting for multiple testing in any subgroup, although it was nominally associated with the ↑WS subgroup.

None of the genes identified above have been associated with MDD or depression in recent large scale meta-analyses(2, 19, 21, 22). However, *CLSTN2* has recently been nominally associated with remission after antidepressant treatment(44). In rodents, increased expression of *CLSTN2* has been observed following a three-week administration of the antidepressant Venlefaxine, a serotonin-noradrenaline reuptake inhibitor(45). *AKNAD1* contains a SNP previously reported as negatively associated with recurrent MDD(46). This is consistent with the ↓WS and ↓W subgroups, as both were observed to have a significantly lower proportion of individuals reporting ‘Too many [episodes] to count’ or ‘One [episode] ran into the next’. Loci in the *LSAMP* gene have previously been associated with a number of psychiatric disorders including MDD, panic disorder and schizophrenia (reviewed by Innos *et al*. (47)). *EPHB1*, *LSAMP* and *ZNF365* have been associated with both insomnia and chronotype(48–50) and a locus in *CLSTN2* with a negative effect was associated with insomnia(48). *ALK*, *EPHB1* and *LSAMP* have all previously been associated with BMI(51–53). Additionally, *ALK* and *CLSTN2* have been associated with glucose homeostasis(54, 55) and *ALK* and *AKNAD1* have been associated with cholesterol levels(56–58). Lastly, *RGS14*, *AKNAD1* and *ZNF365* have all been associated with white blood cells(59, 60) and *LSAMP*, *RGS14, EPHB1, CLSTN2, RGS14* and *ZNF365* with inflammatory diseases(61–65). These results support an immunometabolic component to atypical depression, as previously suggested by Milaneschi *et al*(8).

It has been shown that adjusting GWAS for heritable covariates such as BMI can induce spurious associations if there is a non-causal correlation between the trait and covariate(18). Of the genes discussed above, only *CLSTN2*, *AKNAD1*, *ZNF365* and *LSAMP* remained associated with subgroups at suggestive significance or lower when analyses were not adjusted for BMI. However, as the phenotype was pre-adjusted for BMI before genetic analyses were performed, this may be due to a reduction in residual variance of the sample increasing the power to detect effects not mediated by BMI(66). When BMI was instead included as a covariate in association analyses, results for the loci in *ALK, EPHB1 and CLSTN2* were no different from the results observed in unadjusted analyses (Supplementary Table 33). The *LSAMP* locus showed little difference in effect size between results unadjusted for BMI, pre-adjusted for BMI or when BMI was included as a covariate (Supplementary Table 33). Of these four loci, only the *LSAMP* locus, rs4688064, was in significant linkage disequilibrium with a variant previously associated with BMI (rs768917(59, 67), d’ = 0.88, r^2^ = 0.59, *p* < 0.0001, Supplementary Table 34). Previous Mendelian randomisation studies also suggest there is a causal influence of BMI on atypical depression defined using weight and sleep(68), and the atypical symptom of increased appetite(69). In addition, subgroup-subgroup genetic correlations including the ↑WS subgroup were significantly lower than 1 when analyses were adjusted for BMI. Interestingly, this was not observed when weight-only subgroup definitions were used in sensitivity analyses. As the ↑W subgroup also had a significantly higher BMI than other subgroups, this suggests that the inclusion of sleep in subgroup definitions is a critical component. Overall, until the relationship between atypical depression and BMI is fully understood, it seems reasonable to treat results only observed in analyses adjusted for BMI with a degree of caution. Whilst acknowledging that with careful interpretation, performing both analyses both adjusted and unadjusted for BMI can be informative.

An important limitation to these analyses is the small sample of ↑WS individuals (n = 1,525). Analyses outside the UK Biobank have utilised weight/appetite-only definitions, as these symptoms are highly discriminatory(7) and result in larger samples than weight/appetite-sleep definitions. All analyses were repeated using broader, weight-only, definitions increasing the number of individuals representing atypical depression (n = 5,015). It was not possible to include information on appetite change due to data constraints. Although there were similarities between weight-only subgroups and their weight-sleep counterparts, some significant characteristics were not captured when information on sleep change was excluded from the definition. It is also important to note that, despite being highly discriminatory, weight and sleep symptoms are only a subset of those described in the DSM-5 criteria for atypical depression(3) and that weight change during worst episode could be influenced by antidepressant use(70). Additionally, measures were not all recorded at the same time. In particular, BMI was recorded at initial assessment, whilst depression status was derived from more recent online questionnaires. It is possible BMI could fluctuate between these time points, which could have implications for results as the relationship between BMI and weight-sleep subgroups is not fully understood. Other limitations to consider are that the UK Biobank participants are more likely to be female, older and live in less socioeconomically deprived areas(71). The generalisability of findings are further limited by the restriction of analyses to those with White British ancestry. Finally, both lifetime MDD and the depression subgroups were defined using self-reported depression symptoms during worst episode which could be subject to recall bias(72).

In conclusion, results of this work suggest there are important differences between ↑WS, ↓WS and uncategorised subgroups of depression which potentially identify more chronic and acute forms of MDD. Importantly, a specific enrichment for epilepsy in the ↑WS subgroup was identified, which appears to drive the overall association between epilepsy and lifetime MDD. In addition, several novel genetic associations with subgroups were identified which provide further evidence for an immunometabolic component to ↑WS depression. Future research should work towards fully elucidating this component. However, these analyses require replication in larger, independent samples, particularly those in the ↑WS subgroup.

## Acknowledgements

SM is supported by the Medical Research Council (MR/N013166/1). RJS is supported by a University of Glasgow LKAS fellowship and a UKRI Innovation-HDR-UK Fellowship (MR/S003061/1). AMMc is supported by the Wellcome Trust (104036/Z/14/Z and 220857/Z/20/Z). MJA is supported by Wellcome Trust (220857/Z/20/Z) awards to AMMc. CH was supported by an MRC University Unit Programme Grant “QTL in Health and Disease” (U. MC. UU 00007/10).

This research has been conducted using data from UK Biobank, a major biomedical database (https://www.ukbiobank.ac.uk/).

Generation Scotland received core support from the Chief Scientist Office of the Scottish Government Health Directorates [CZD/16/6] and the Scottish Funding Council [HR03006] and is currently supported by the Wellcome Trust [216767/Z/19/Z]. Genotyping of the GS:SFHS samples was carried out by the Genetics Core Laboratory at the Edinburgh Clinical Research Facility, University of Edinburgh, Scotland and was funded by the Medical Research Council UK and the Wellcome Trust (Wellcome Trust Strategic Award “STratifying Resilience and Depression Longitudinally” (STRADL) Reference 104036/Z/14/Z).

## Conflict of Interest

The authors declare no conflicts of interest.

## Notes

### Competing Interest Statement

The authors have declared no competing interest.

### Summary of Updates

(i) Subgroup SNP-heritability results have been removed. (ii) Results and discussion include additional analysis testing the association between subgroup PRS derived in UK Biobank and subgroups defined in an independent sample. (iii) Discussion has been reworked to better highlight how work builds on previous studies in UK Biobank and clarify interpretation of results from genetic analyses adjusted for BMI.

