## Supplementary_Materials for "Characterisation and Genetic Architecture of Major Depressive Disorder Subgroups Defined by Weight and Sleep Changes"

**Supplementary Methods**

**UK Biobank**

***Participants***

The UK Biobank is a population-based study which includes over 500,000 genotyped and extensively phenotyped participants aged between 40-69 at recruitment. The cohort has been described in detail elsewhere(1). This research was conducted using the UK Biobank Resource—application number 4844. UK Biobank received ethical approval from the NHS National Research Ethics Service North West (reference: 11/NW/0382). All participants gave full informed written consent. 157,366 of UK Biobank participants have completed an online Mental Health Questionnaire (MHQ), based on the Composite International Diagnostic Interview Short Form (CIDI-SF), which aims to identify mental health conditions without an in person clinical assessment(2). Only unrelated individuals of genetically defined White British ancestry as in Howard *et al*.(3), that had completed the MHQ were included in these analyses, participants in the Generation Scotland cohort and their relatives were also excluded (n = 122,639).

***Genotyping and Quality Control***

Genotyping of participants was performed using two arrays (the Applied Biosystems UK BiLEVE Axiom Array and the Applied Biosystems UK Biobank Axiom Array), which are highly overlapping and cover ~800,000 markers(4). Variants were excluded from analyses if they had a minor allele frequency (MAF) < 0.01, a high level of missingness (>0.05) or showed evidence of departure from Hardy-Weinberg equilibrium (HWE, *p* < 1 × 10^-5^).

***Phenotype Definitions***

***Probable Lifetime MDD***

Probable lifetime MDD cases (n = 26,662) were identified using the criteria outlined by Davis et al.(2). Briefly, these were individuals that reported depressed mood or anhedonia (or both) lasting longer than two weeks and impairing their daily life, as well as additional DSM-5 MDD symptoms (at least four in total) during their worst depressive episode. Individuals with a psychotic, manic, personality or substance abuse disorder (recorded in electronic health records or self-reported to a trained nurse at baseline, in the touchscreen assessment or in the MHQ) were excluded in line with DSM-5 criteria for depression(5). Further detail on the identification of individuals with probable lifetime MDD can be found in **Supplementary Figure 1.**

***Depression Subgroups***

Depression subgroups were defined in line with previous studies of subgroups with reversed neurovegetative symptoms in the UK Biobank(6, 7). Individuals reporting increased weight (UKB field 20536) and sleeping more (UKB field 20534) during their worst episode were categorised as increased weight and sleep (↑WS, n = 1,525) representing atypical depression. Individuals reporting decreased weight (UKB field 20536) and sleeping less (UKB field 20533 or UKB field 20535) during their worst episode were categorised as decreased weight and sleep (↓WS, n = 9,067) representing typical depression. Individuals with lifetime MDD that did not fall into either subgroup were included in analyses as a third ‘uncategorised’ subgroup (n = 16,070) (**Supplementary Figure 1**).

***Controls***

Individuals that reported never having experienced a depressed mood and anhedonia for more than two weeks were classified as controls (UKB fields 20446 & 20441). Unlike in Davis *et* al.(2), individuals were not excluded from the control group based on patient health questionnaire 9 (PHQ-9) score to avoid heavily screened so-called ‘clean’ controls, which can introduce bias to heritability estimates transformed to the liability scale(8, 9). Individuals with a depressive, psychotic, manic, personality or substance abuse disorder recorded in electronic health records (ICD10 codes, F05-07, F09, F10-39) or self-reported to a trained nurse at baseline (UKB field 20002), in the touchscreen assessment (UKB field 20126) or in the MHQ (UKB field 20544), were excluded from the control group.

**Generation Scotland**

***Participants***

The Generation Scotland: Scottish Family Health Study (GS:SFHS) is a family and population study including 23,690 adult participants recruited from Scottish general practices. The cohort is described in detail at by Smith *et al.*(10, 11). In 2015, 8,541 participants were recruited to a follow-up mental health study (Stratifying Resilience and Depression Longitudinally, STRADL(12)), where participants submitted responses to a questionnaire based on the CIDI-SF. STRADL received ethical approval from the NHS Tayside committee (reference 14/SS/0039). Only unrelated individuals were included in analyses, filtering of related individuals was performed in a manner that maximised the sample of the smallest group in each comparison. Samples sizes for each comparison are shown in **Supplementary Table 2**.

***Genotyping and Quality Control***

Genotype data were available for 18,725 participants. Individuals with ≥2% missingness were excluded from analyses. As were variants with a call rate ≤98%, Hardy Weinberg Equilibrium (HWE) *p*-value ≤ 1 × 10^-6^ and minor allele frequency MAF ≤1%.

***Phenotype Definitions***

Probable lifetime major depressive disorder (MDD) cases were defined in manner consistent with UK Biobank cases using participant responses to the CIDI-SF. These were individuals reporting a depressed mood or lack of interest of pleasure (anhedonia), as well as four additional DSM-5 symptoms, lasting for two weeks or more. In addition to be categorised as a case, participants had to report symptoms lasting ‘all day long’ or ‘most of the day’ and occurring ‘every day’ in this two-week period. Depression subgroups were defined using the same definition as in UK Biobank. Those reporting weight gain and sleeping more during worst episode were classified as ↑WS (n = 133), those reporting weight loss and trouble sleeping or waking early were classified as ↓WS (n = 575). All the remaining individuals were included in analyses as a third uncategorised subgroup (n = 798). All analyses were repeated in subgroups using a broader weight-only definition: weight gain (↑W, n = 397), weight loss (↓W, n = 667) and uncategorised (n = 442). Individuals without a two-week period of depressed mood or anhedonia, four additional DSM-V symptoms or with symptoms not lasting ‘all day long’ or ‘most of the day’ ‘every day’ for that two-week period were classified as controls (n = 7667). Individuals with symptoms of bipolar disorder or hypomanic episodes were excluded from analyses.

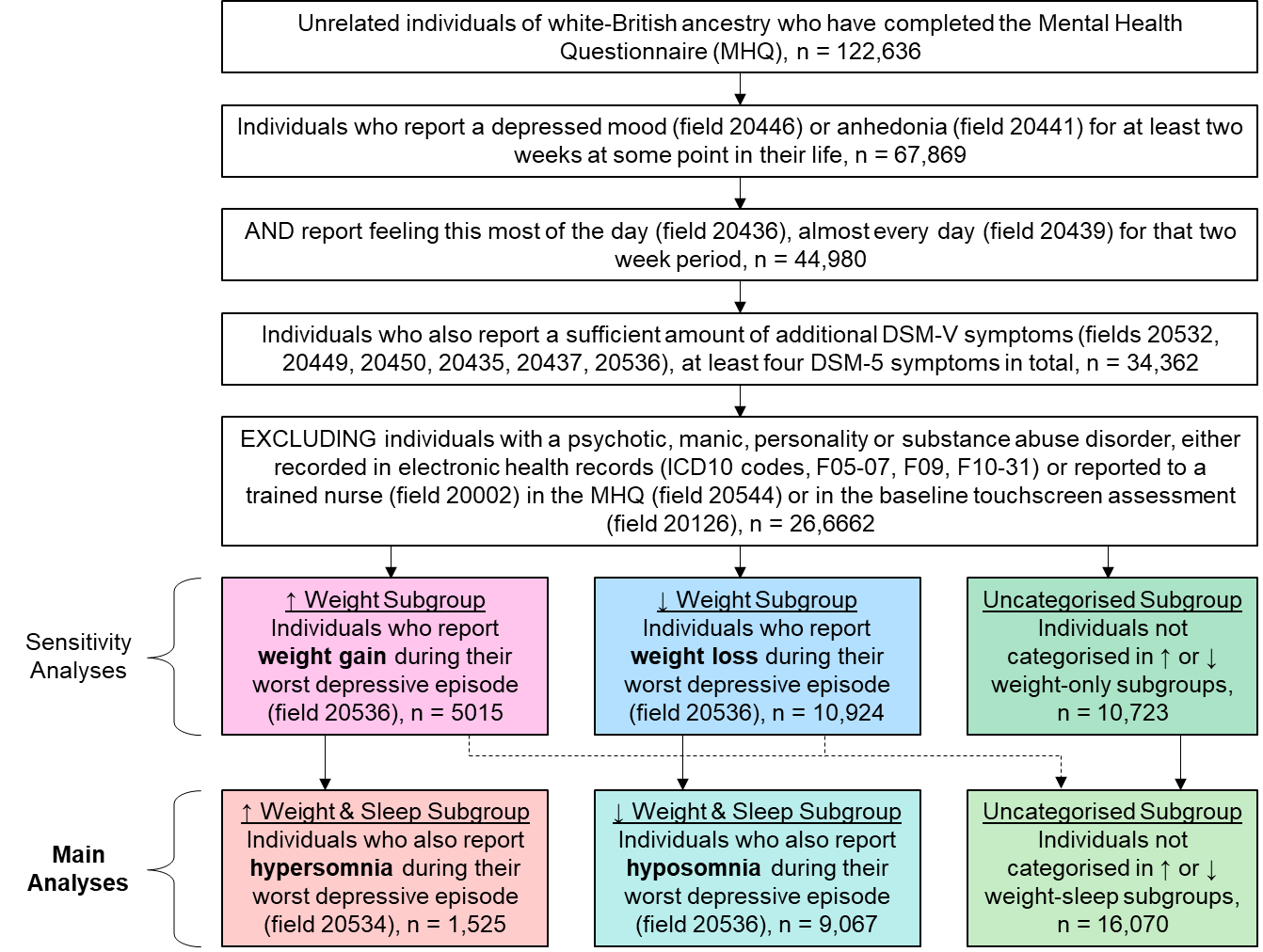
**Supplementary Figure 1**. Flowchart of definition of probable major depressive disorder and weight and sleep depression subgroups in UK Biobank

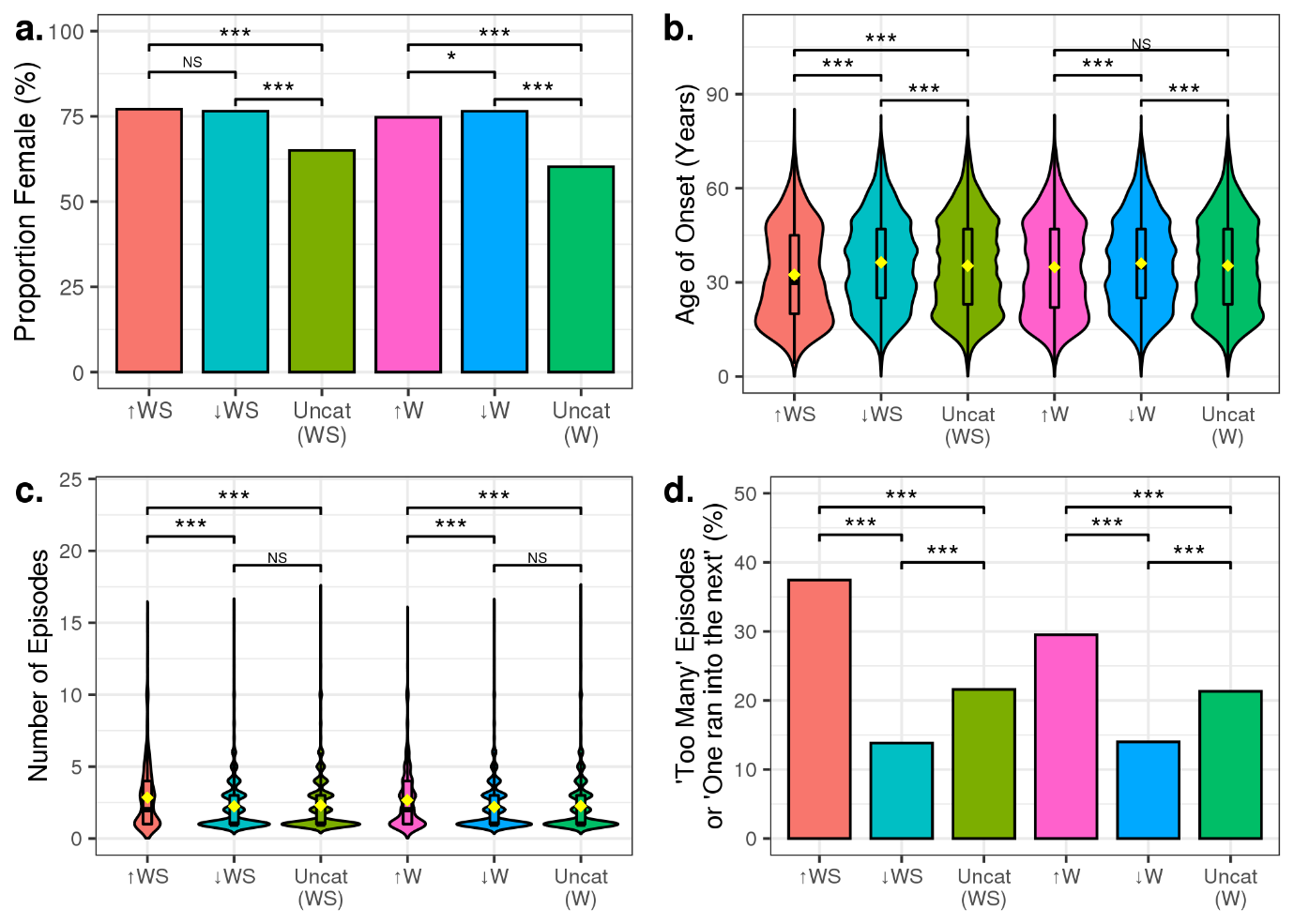
**Supplementary Figure 2.** The sample and depression characteristics of weight-sleep and weight-only depression subgroups

*The sample and depression characteristics of the depression subgroups: (a) the proportion of females (%), (b) age of onset, (c) number of depressive episodes, (d) proportion reporting ‘Too many episodes to count’ or ‘One episode ran into the next’. Proportions are shown as exact values in bar charts. Otherwise, the distribution of data is shown as boxplot of the median and interquartile range, whiskers for each boxplot are calculated using the formula: quartile + 1.5*IQR and outliers are not shown. Significant comparisons are denoted as follows: *** (p < 0.001), ** (p < 0.01), * (p < 0.05).*

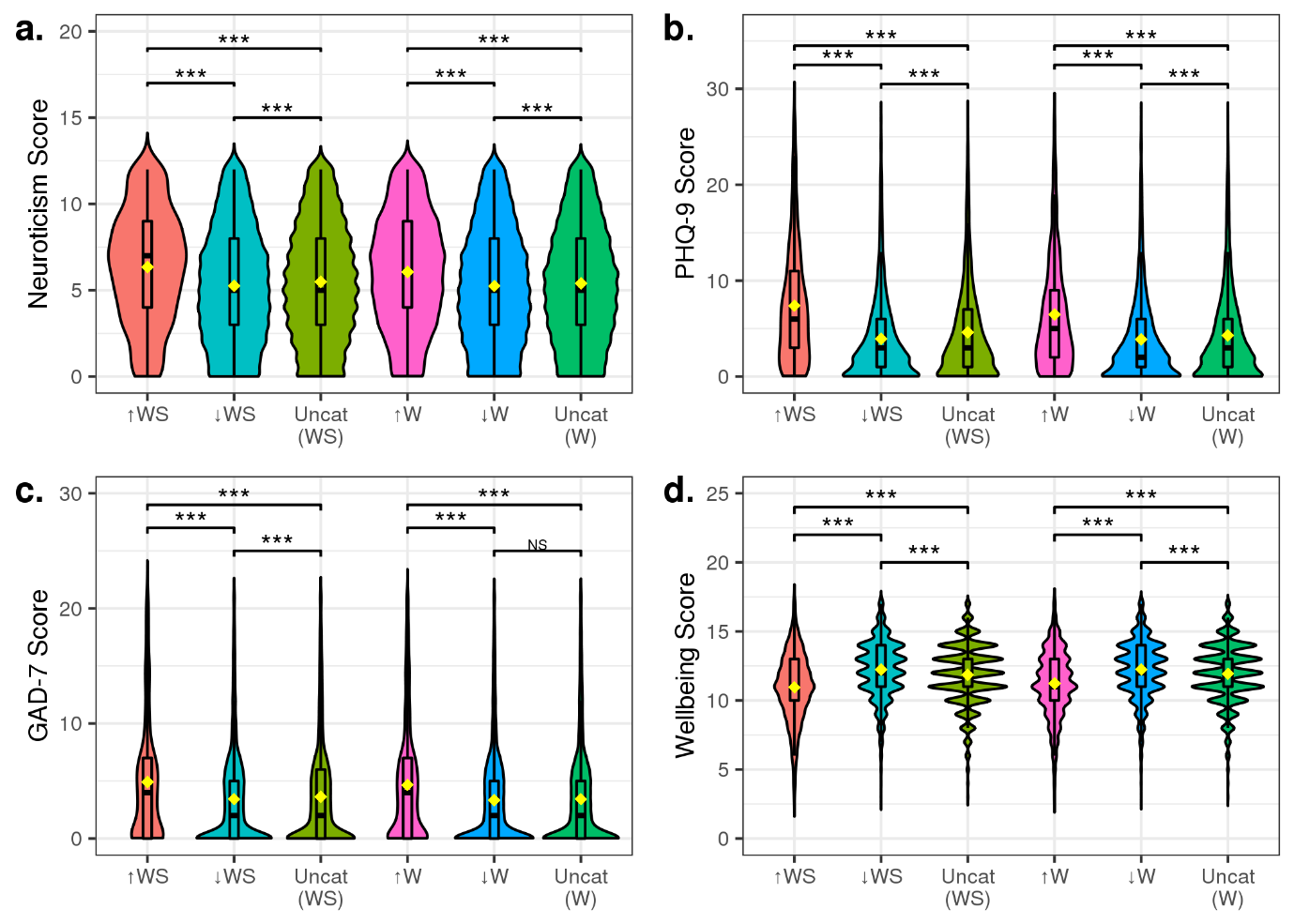
**Supplementary Figure 3**. The mental health scores of weight-sleep and weight-only depression subgroups

*The mental health scores: (a.) neuroticism, (b.) PHQ-9, (c.) GAD-7, and (d.) wellbeing of each subgroup. The distribution of data is shown as a boxplot of the median and interquartile range, whiskers for each boxplot are calculated using the formula: quartile + 1.5*IQR and outliers are not shown. Significant comparisons are denoted as follows: *** (p < 0.001), ** (p < 0.01), * (p < 0.05).*

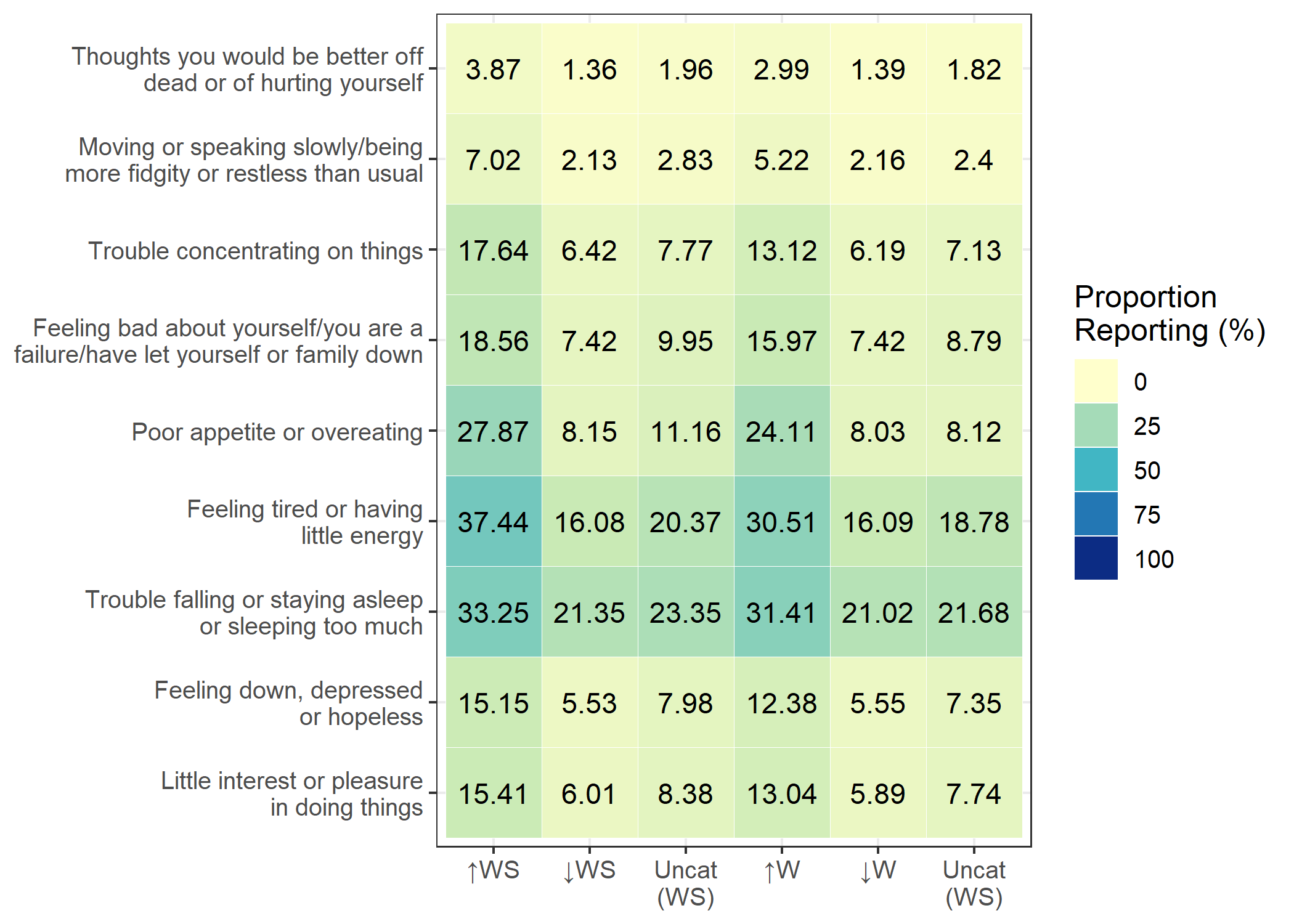
**Supplementary Figure 4.** Heat map of the responses to PHQ-9 questions in each weight-sleep and weight-only depression subgroup

*A heat map of the proportion of individuals who report experiencing each of the PHQ-9 symptoms ‘more than half the days’ or ‘nearly every day’ over the last two weeks, at the time of completing the UK Biobank mental health questionnaire, for each weight-sleep and weight-only subgroup.*

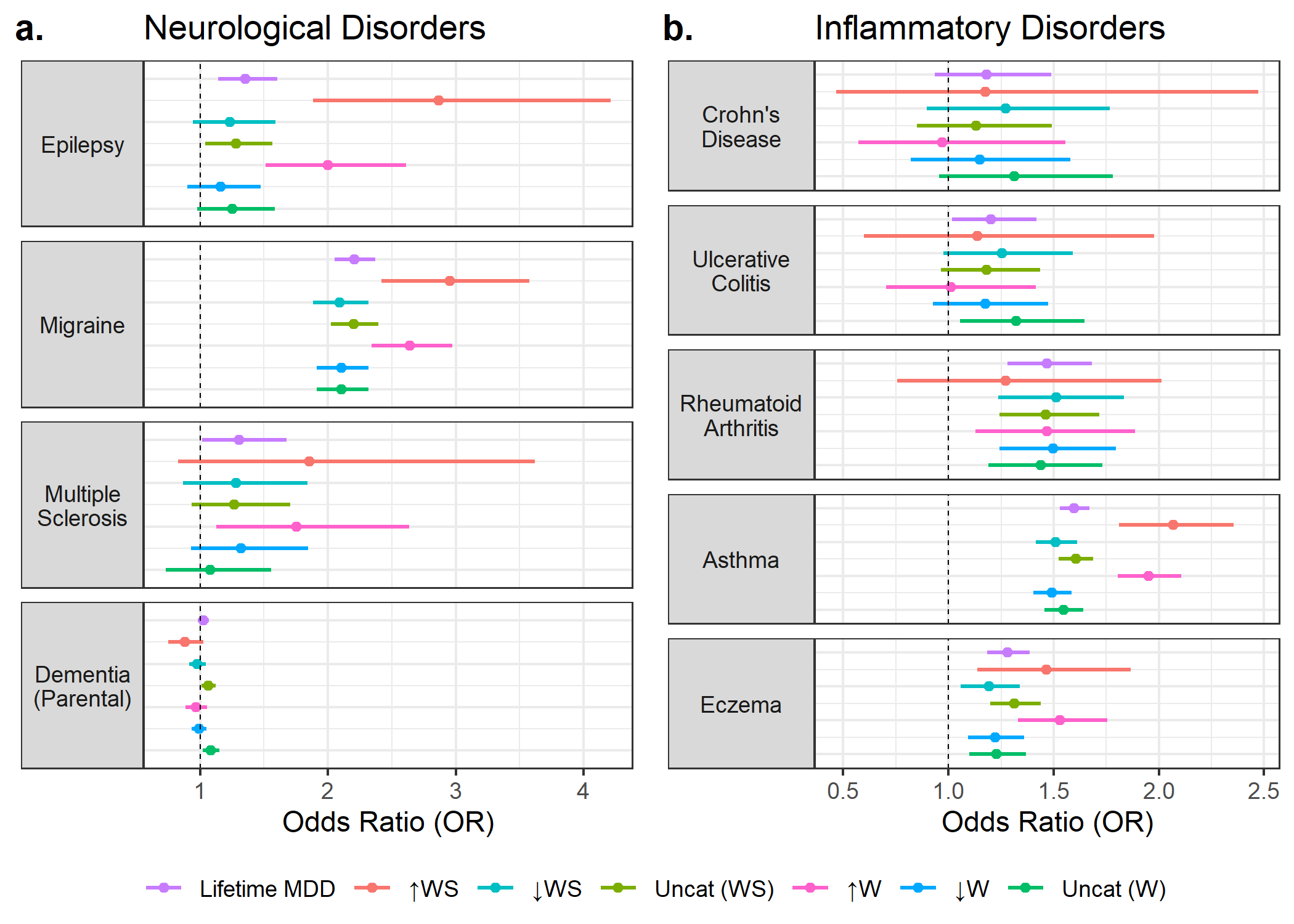
**Supplementary Figure 5.** Prevalence of neurological and inflammatory disorders in weight-sleep and weight-only depression subgroups

*The odds ratios and 95% confidence intervals of lifetime MDD and each weight-sleep and weight-only subgroup compared to controls for (a.) four neurological and (b.) five inflammatory disorders of interest.*

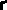

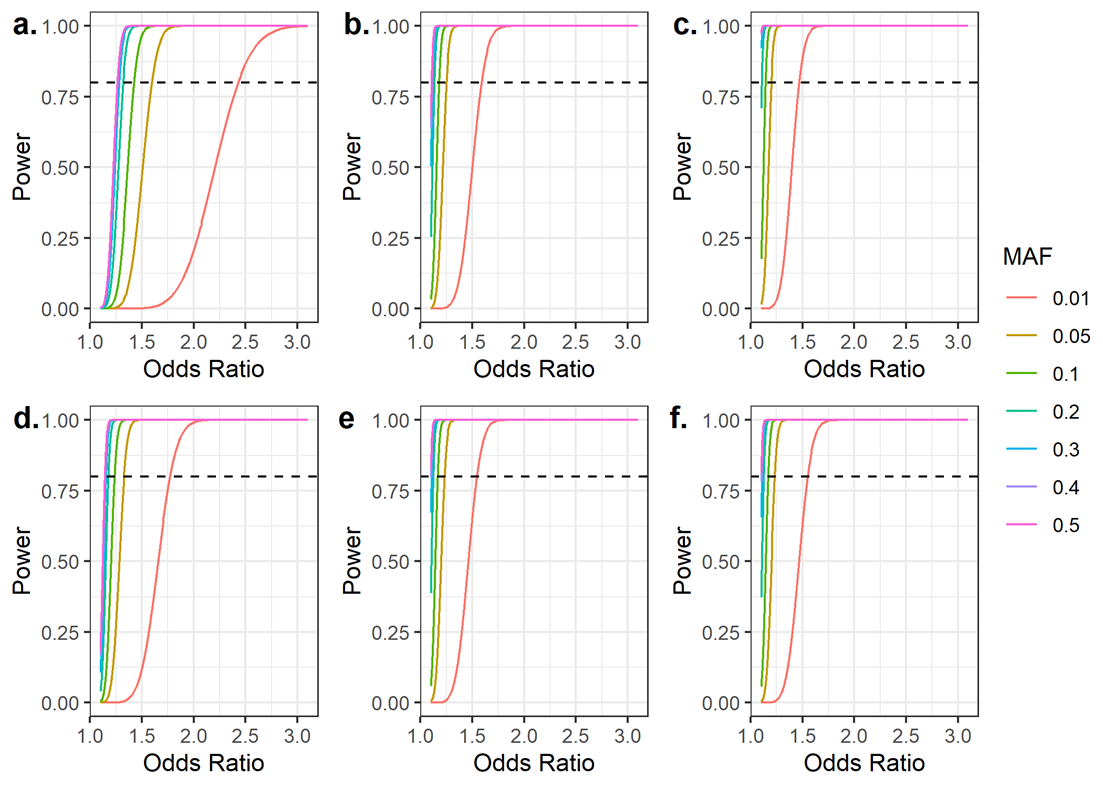
**Supplementary Figure 6.** Power calculations for case-control GWAS

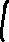

*Results of post-hoc power calculations for case-control GWAS across a range of minor allele frequencies. (a.) ↑WS vs controls, (b.) ↓WS vs controls, (c.) uncategorised (WS) vs controls, (d.) ↑W vs controls, (e.) ↓W vs controls, (f.) uncategorised (W) vs controls. Calculations assumed an additive logistic model.*

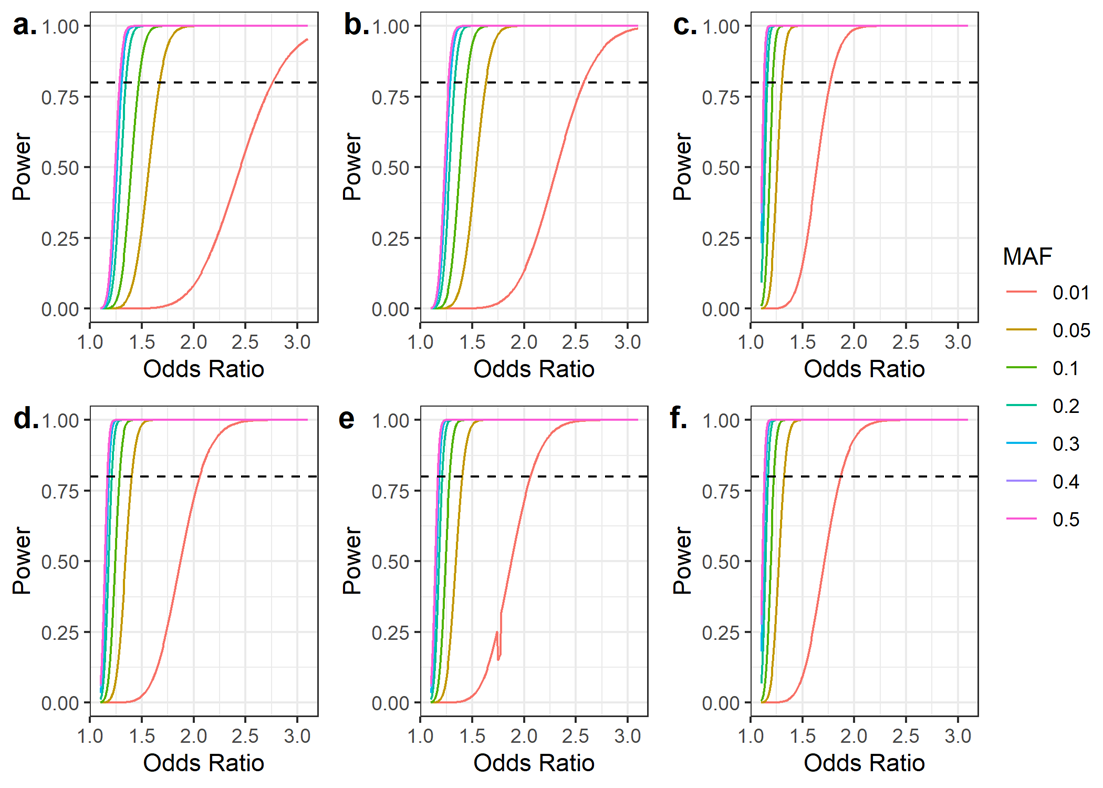
**Supplementary Figure 7.** Power calculations for case-only GWAS

*Results of post-hoc power calculations for case-control GWAS across a range of minor allele frequencies. (a.) ↑WS vs ↓WS, (b.) ↑WS vs uncategorised (WS), (c.) ↓WS vs uncategorised (WS), (d.) ↑W vs ↓W, (e.) ↑W vs uncategorised (W), (f.) ↓W vs uncategorised (W). Calculations assumed an additive logistic model.*

**Supplementary Figure 8**. Regional plot of *ALK* locus – Case-control GWAS of ↑WS subgroup

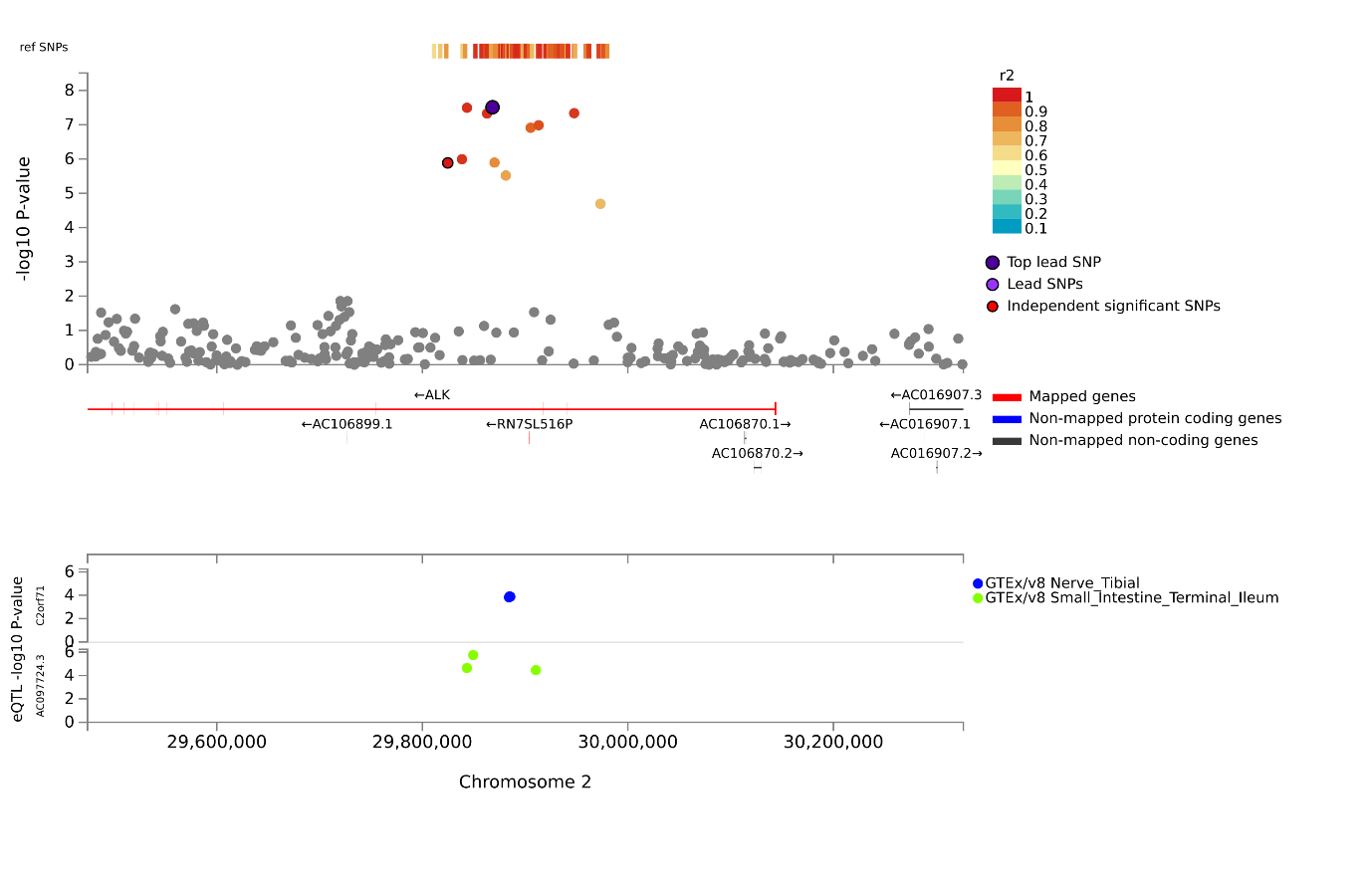

*Regional plot created in FUMA for the ALK locus associated with the ↑WS subgroup at genome-wide significance in analyses adjusted for BMI. Shown are the -log10 P-values of SNPs in the region; the mapped, non-mapped protein coding and non-mapped non-coding genes; as well as associated eQTLs and their -log10 P-value.*

**Supplementary Figure 9**. Regional plot of *EPHB1* locus – Case-control GWAS of ↑WS subgroup

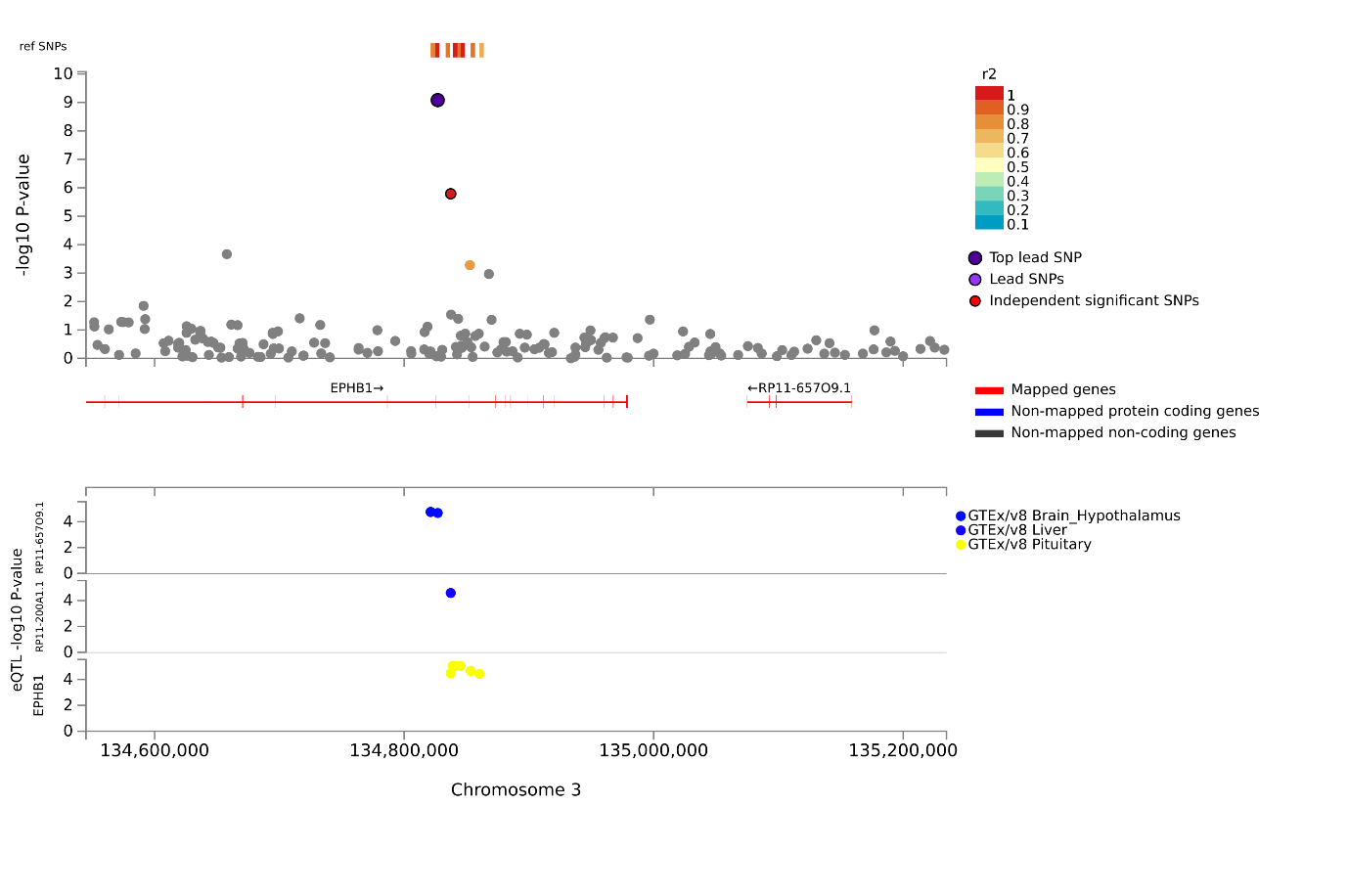

*Regional plot created in FUMA for the EPHB1* *locus associated with the ↑WS subgroup at genome-wide significance in analyses adjusted for BMI. Shown are the -log10 P-values of SNPs in the region; the mapped, non-mapped protein coding and non-mapped non-coding genes; as well as associated eQTLs and their -log10 P-value.*

**Supplementary Figure 10**. Regional plot of *CLSTN2* locus – Case-control GWAS of ↑WS subgroup

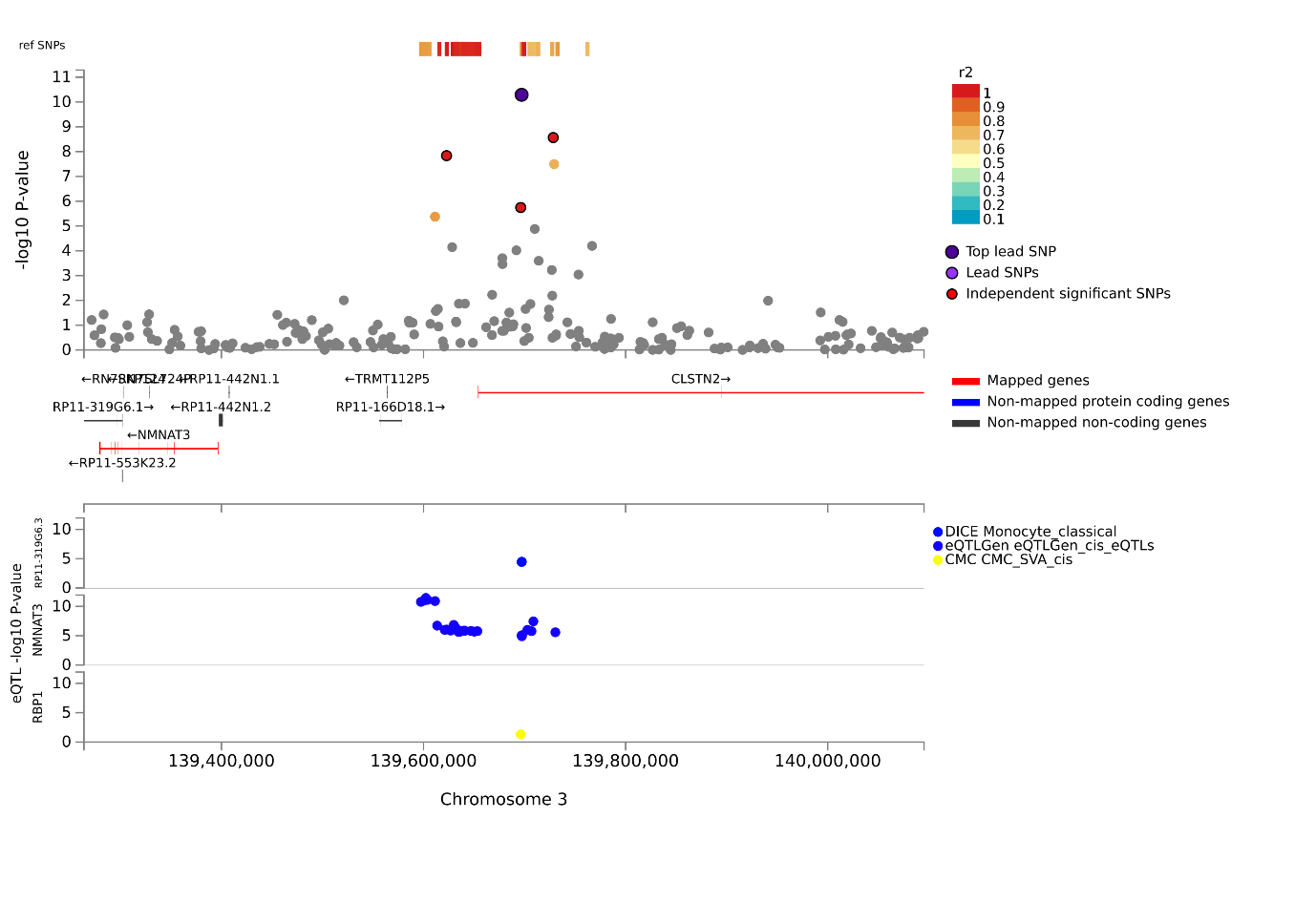

*Regional plot created in FUMA for the CLSTN2* *locus associated with the ↑WS subgroup at genome-wide significance in analyses adjusted for BMI. Shown are the -log10 P-values of SNPs in the region; the mapped, non-mapped protein coding and non-mapped non-coding genes; as well as associated eQTLs and their -log10 P-value.*

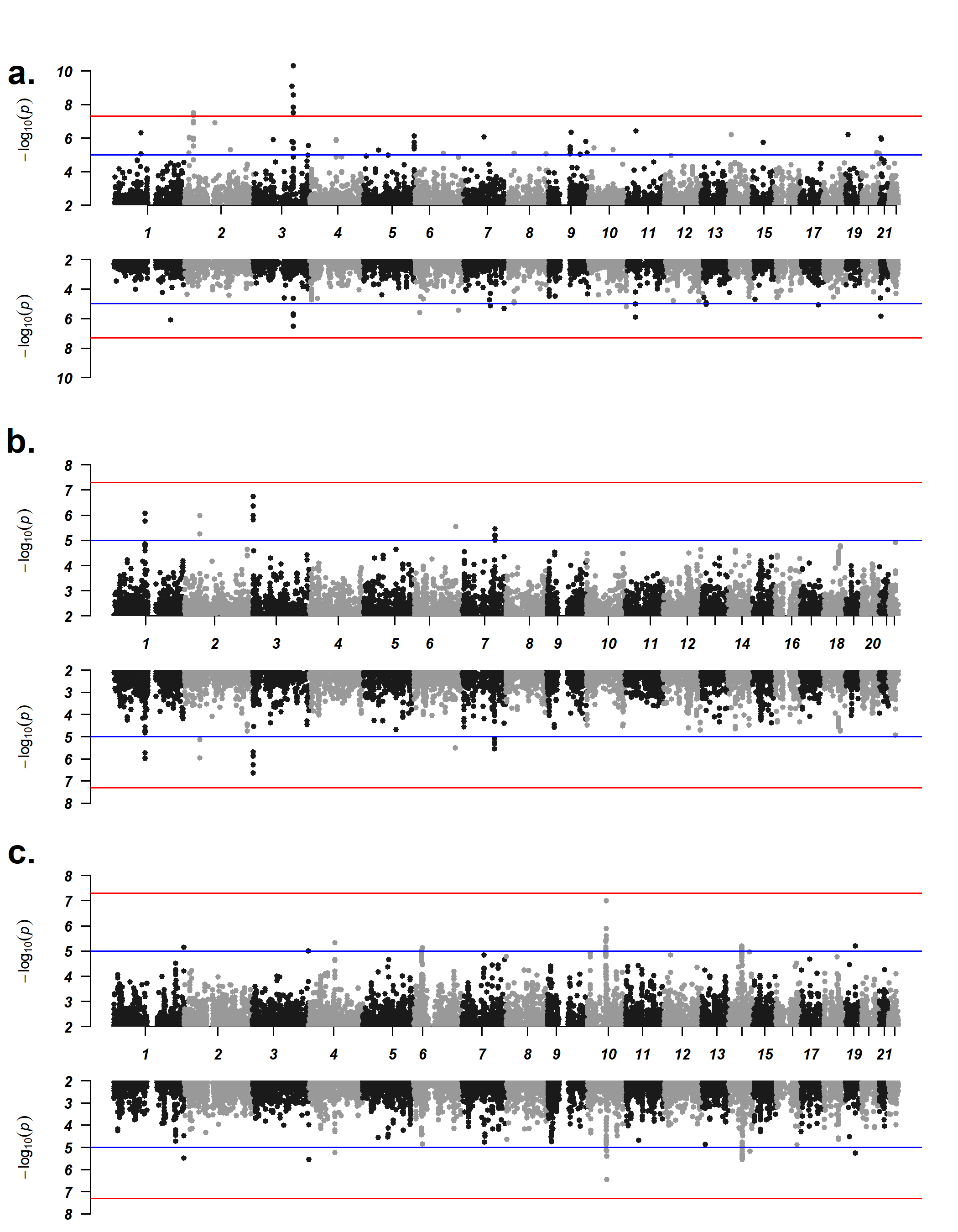
**Supplementary Figure 11**. Miami pots of weight-sleep subgroup case-control GWAS results

Miami plots of the (a.) ↑WS case-control GWAS (b.) ↓WS case-control GWAS and (c.) uncategorised case-control GWAS. Analyses adjusted for BMI are shown in the top plot and analyses unadjusted for BMI are shown in the bottom plot. Chromosomal location is on the x-axis and -log_10_ of the p-value for each SNP is on the y-axis. The red line indicates the threshold for genome-wide significance (5.00 × 10^-8^) and the blue dashed line indicates the threshold for “suggestive” significance (1.00 × 10^-5^).

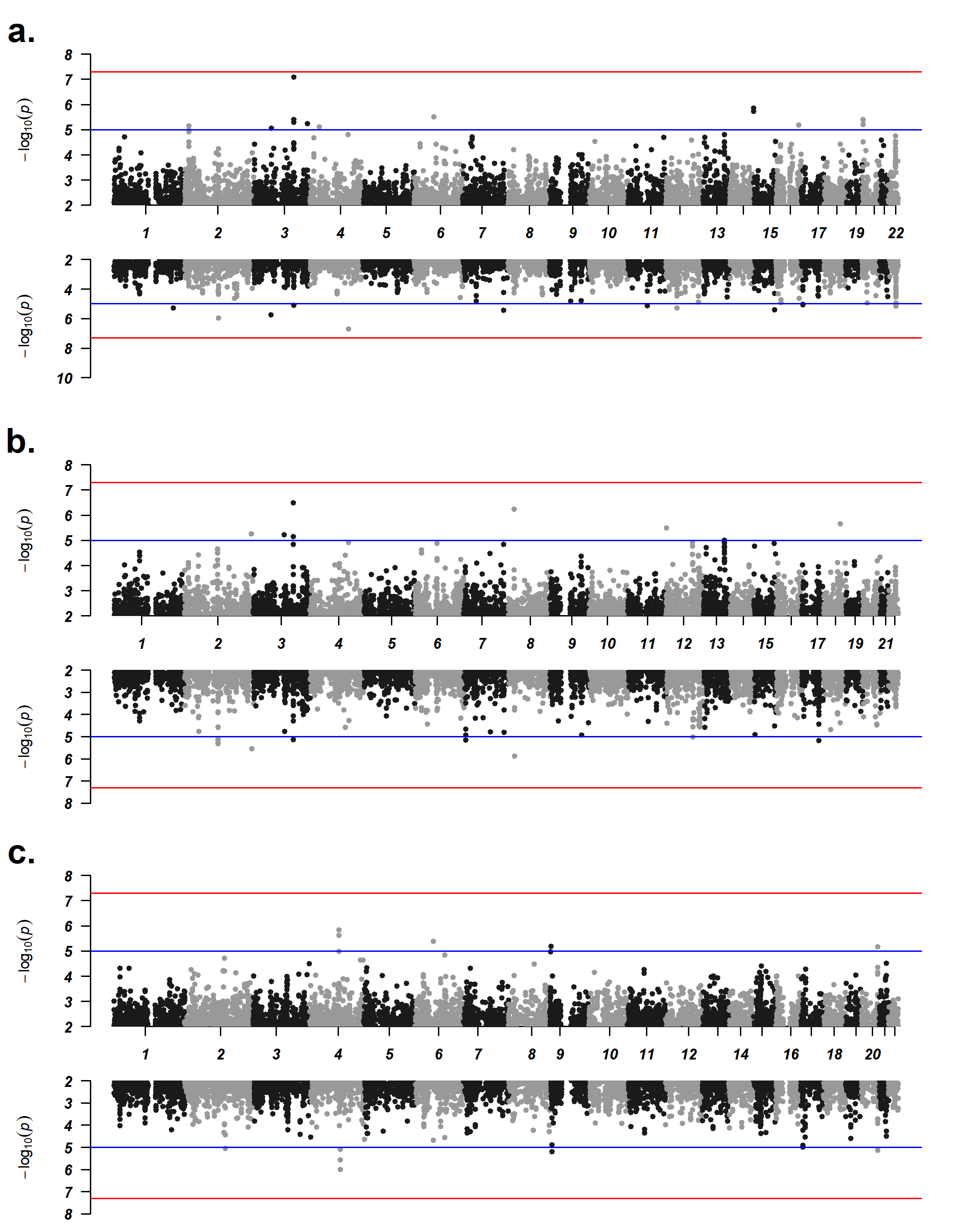
**Supplementary Figure 12**. Miami plots of weight-sleep subgroup case-only GWAS results

Miami plots of the (a.) ↑WS vs ↓WS case-only GWAS (b.) ↑WS and uncategorised case-only GWAS and (c.) ↓WS and uncategorised case-only GWAS. Analyses adjusted for BMI are shown in the top plot and analyses unadjusted for BMI are shown in the bottom plot. Chromosomal location is on the x-axis and -log_10_ of the p-value for each SNP is on the y-axis. The red line indicates the threshold for genome-wide significance (5.00 × 10^-8^) and the blue dashed line indicates the threshold for “suggestive” significance (1.00 × 10^-5^).

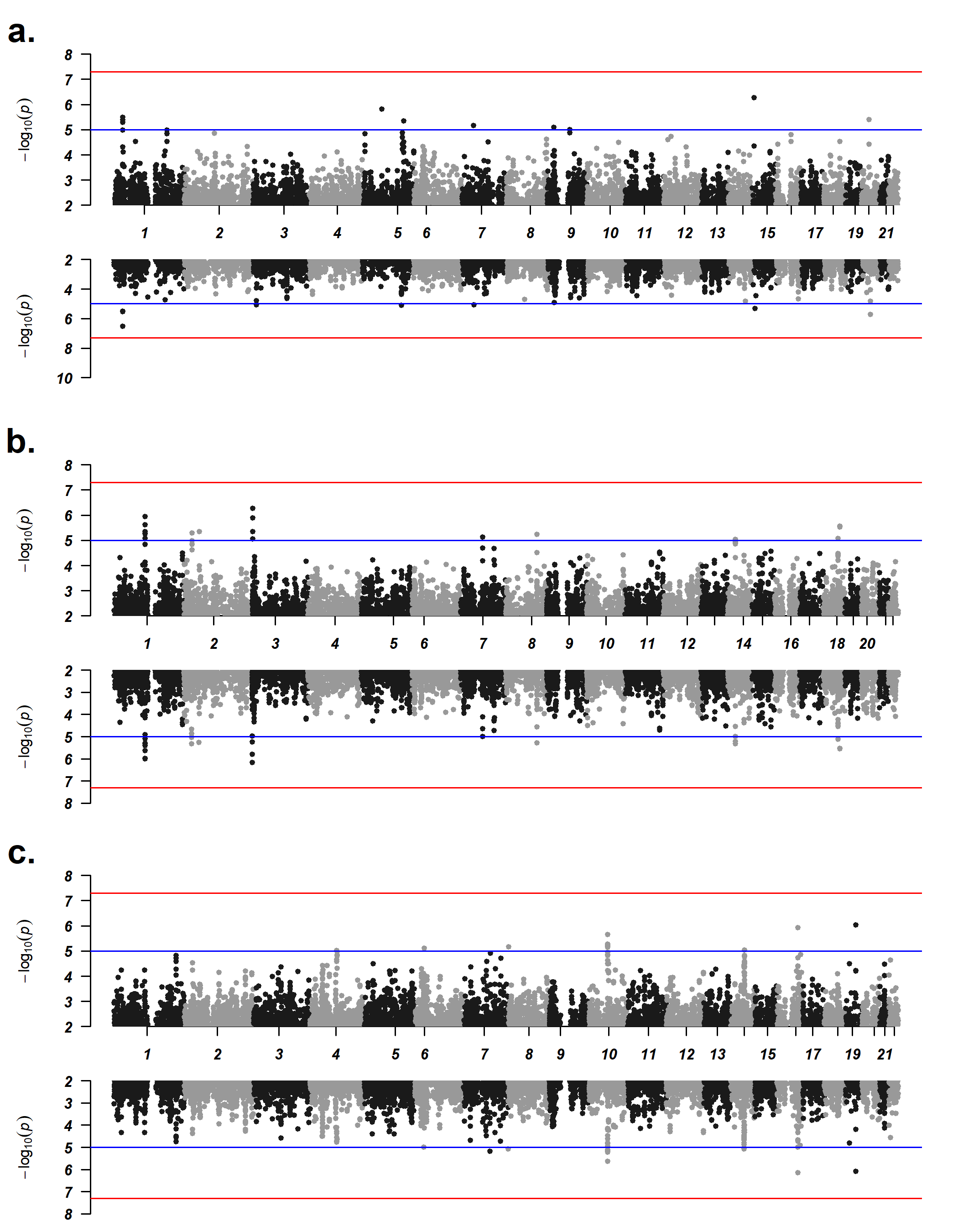
**Supplementary Figure 13**. Miami plots of weight-only subgroup case-control GWAS results

Miami plots of the (a.) ↑W case-control GWAS (b.) ↓W case-control GWAS and (c.) uncategorised case-control GWAS. Analyses unadjusted for BMI are shown in the top plot and analyses adjusted for BMI are shown in the bottom plot. Chromosomal location is on the x-axis and -log_10_ of the p-value for each SNP is on the y-axis. The red line indicates the threshold for genome-wide significance (5.00 × 10^-8^) and the blue dashed line indicates the threshold for “suggestive” significance (1.00 × 10^-5^).

**Supplementary Figure 14**. Miami plots of weight-only subgroup case-only GWAS results

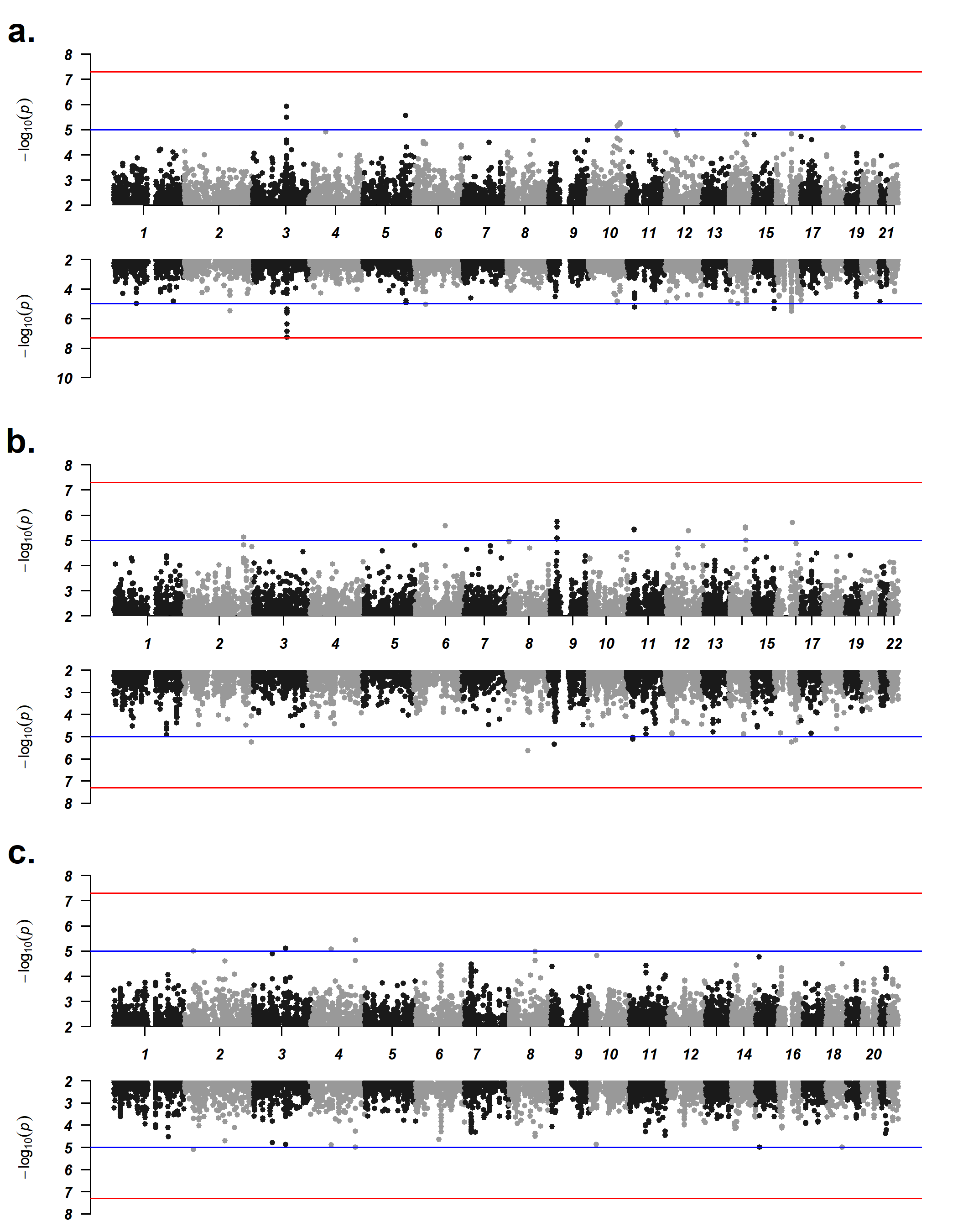
Miami plots of the (a.) ↑W vs ↓W case-only GWAS (b.) ↑W vs uncategorised case-only GWAS and (c.) ↓W vs uncategorised case-only GWAS. Analyses adjusted for BMI are shown in the top plot and analyses unadjusted for BMI are shown in the bottom plot. Chromosomal location is on the x-axis and -log_10_ of the p-value for each SNP is on the y-axis. The red line indicates the threshold for genome-wide significance (5.00 × 10^-8^) and the blue dashed line indicates the threshold for “suggestive” significance (1.00 × 10^-5^).

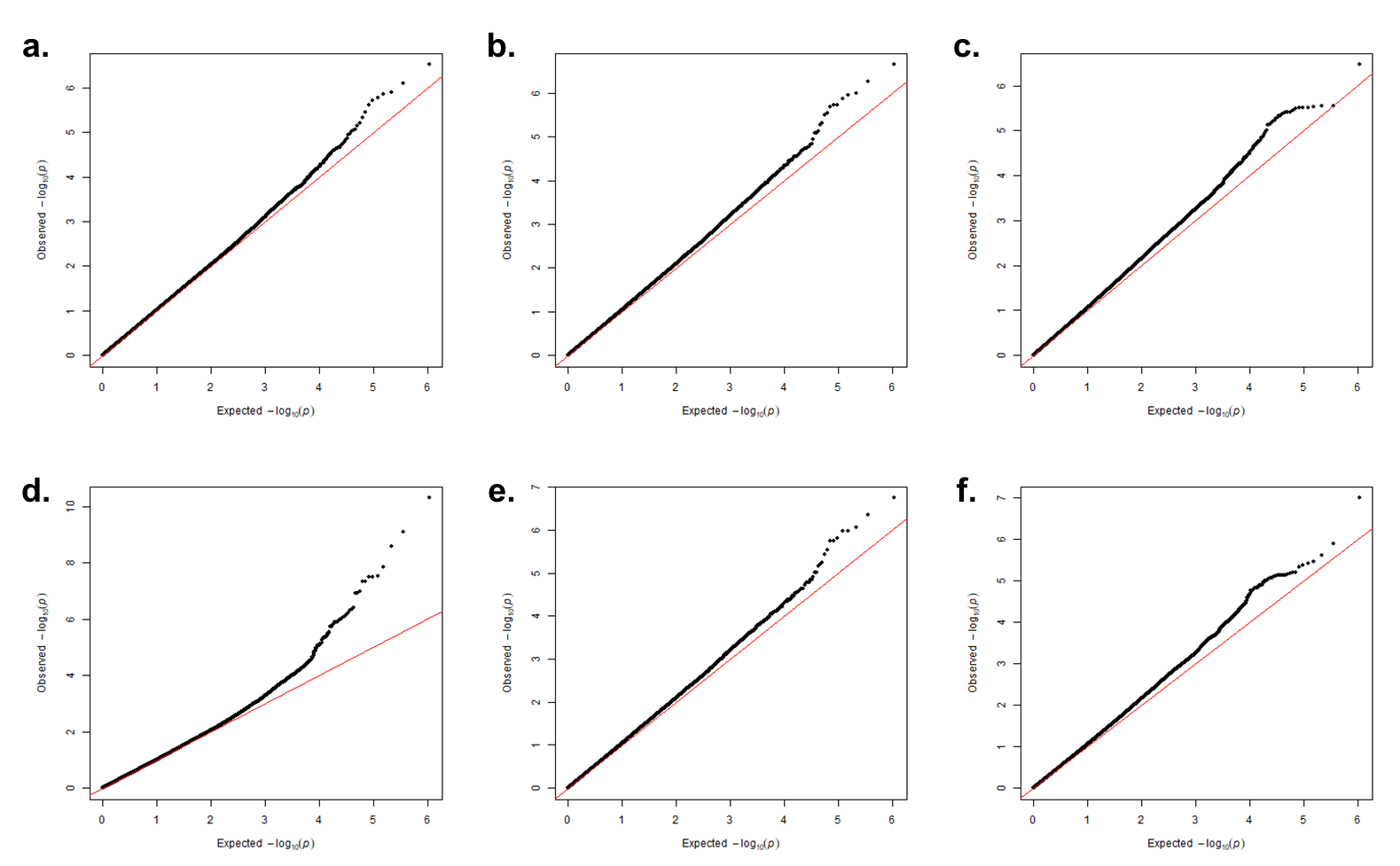
**Supplementary Figure 15.** QQ-plots of the results of the case-control analyses of weight-sleep subgroups unadjusted for BMI: (a.) ↑WS depression (λ = 1.025), (b.) ↓WS depression (λ = 1.054), (c.) Uncategorised depression (λ = 1.071) and adjusted for BMI: (d.) ↑WS depression (λ = 1.012), (e.) ↓WS depression (λ = 1.053), (f.) Uncategorised depression (λ = 1.063). The x-axis shows the expected -log_10_(*p*) and the y-axis the observed -log_10_(*p*).

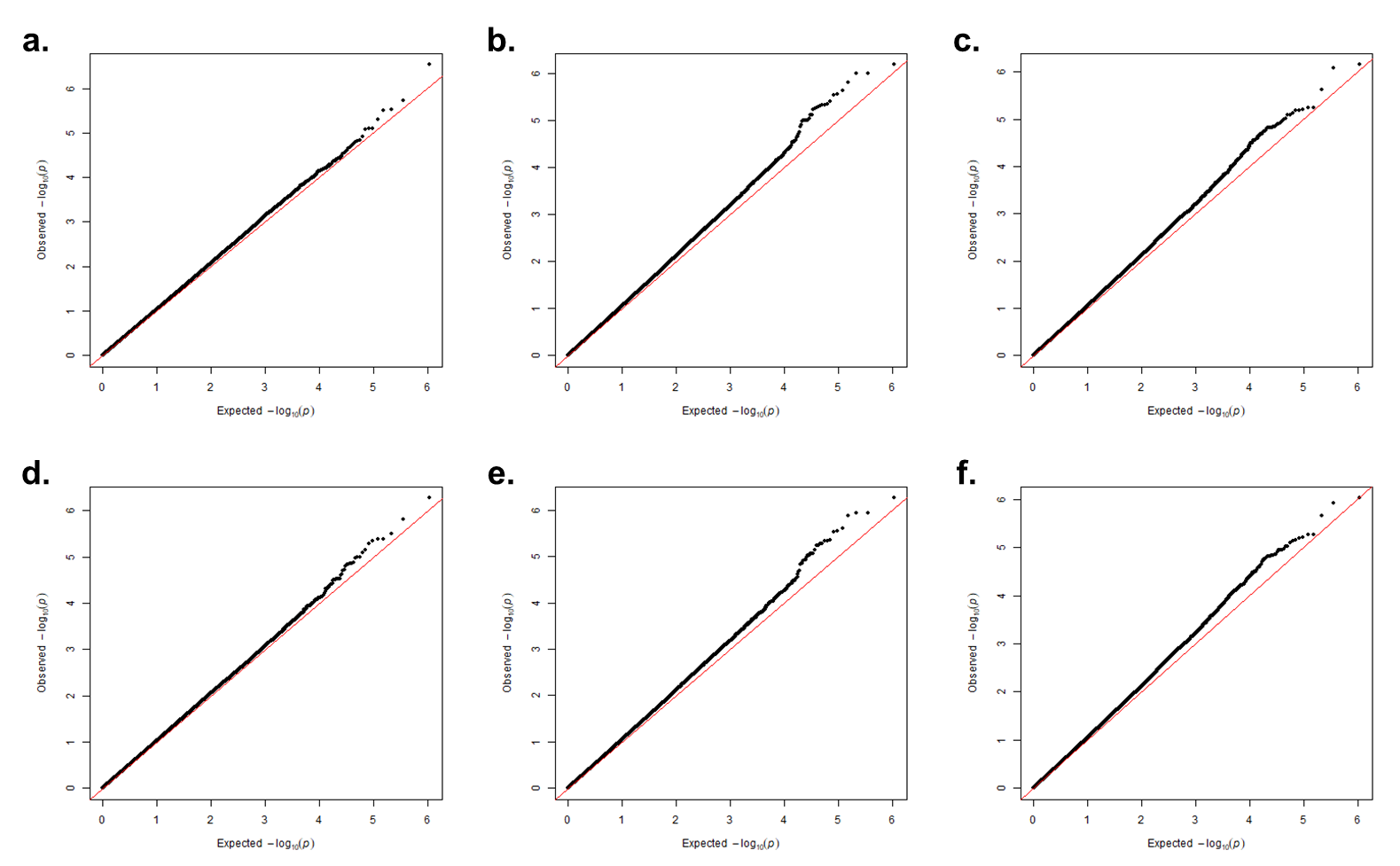
**Supplementary Figure 16.** QQ-plots of the results of the case-control analyses of weight-only subgroups unadjusted for BMI: (a.) ↑W depression (λ = 1.04), (b.) ↓W depression (λ = 1.06), (c.) Uncategorised depression (λ = 1.06) and adjusted for BMI: (d.) ↑W depression (λ = 1.02), (e.) ↓W depression (λ = 1.05), (f.) Uncategorised depression (λ = 1.03). The x-axis shows the expected -log_10_(*p*) and the y-axis the observed -log_10_(*p*).

**Supplementary Figure 17.** QQ-plots of the results of the case-only analyses of weight-sleep subgroups unadjusted for BMI: (a.) ↑WS and ↓WS depression (λ = 1.03), (b.) ↑WS and Uncategorised depression (λ = 1.01), (c.) ↓WS and Uncategorised depression (λ = 1.01) and adjusted for BMI: (d.) ↑WS and ↓WS
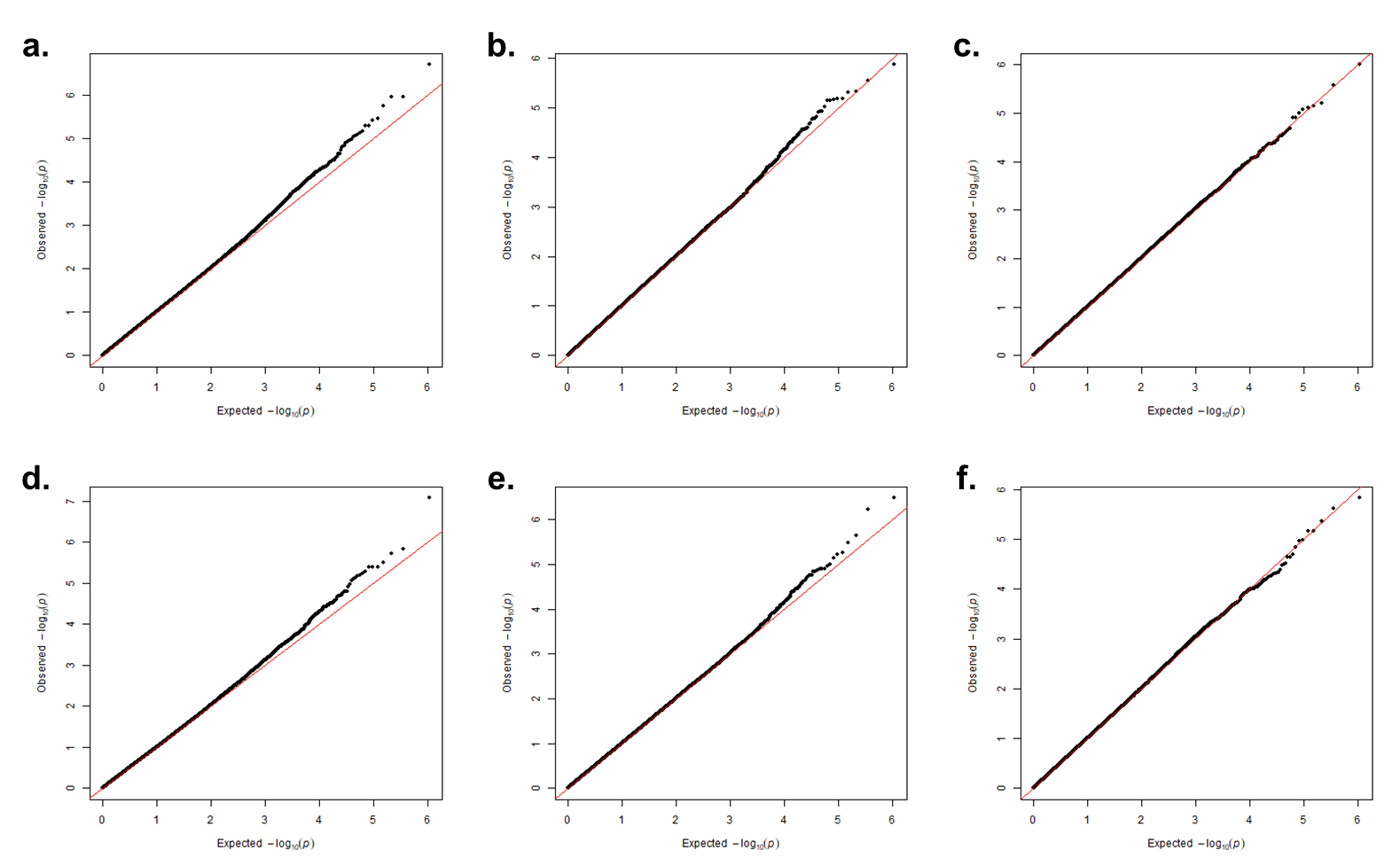
depression (λ = 1.01), (e.) ↑WS and Uncategorised depression (λ = 1.01), (f.) ↓WS and Uncategorised depression (λ = 1.01). The x-axis shows the expected -log_10_(*p*) and the y-axis the observed -log_10_(*p*).

**Supplementary Figure 18.** QQ-plots of the results of the case-only analyses of weight-only subgroups unadjusted for BMI: (a.) ↑W and ↓W depression (λ = 1.03), (b.) ↑W and Uncategorised depression (λ = 1.02), (c.) ↓W and Uncategorised depression (λ = 1.01) and adjusted for BMI: (d.) ↑W and ↓W depression (λ =
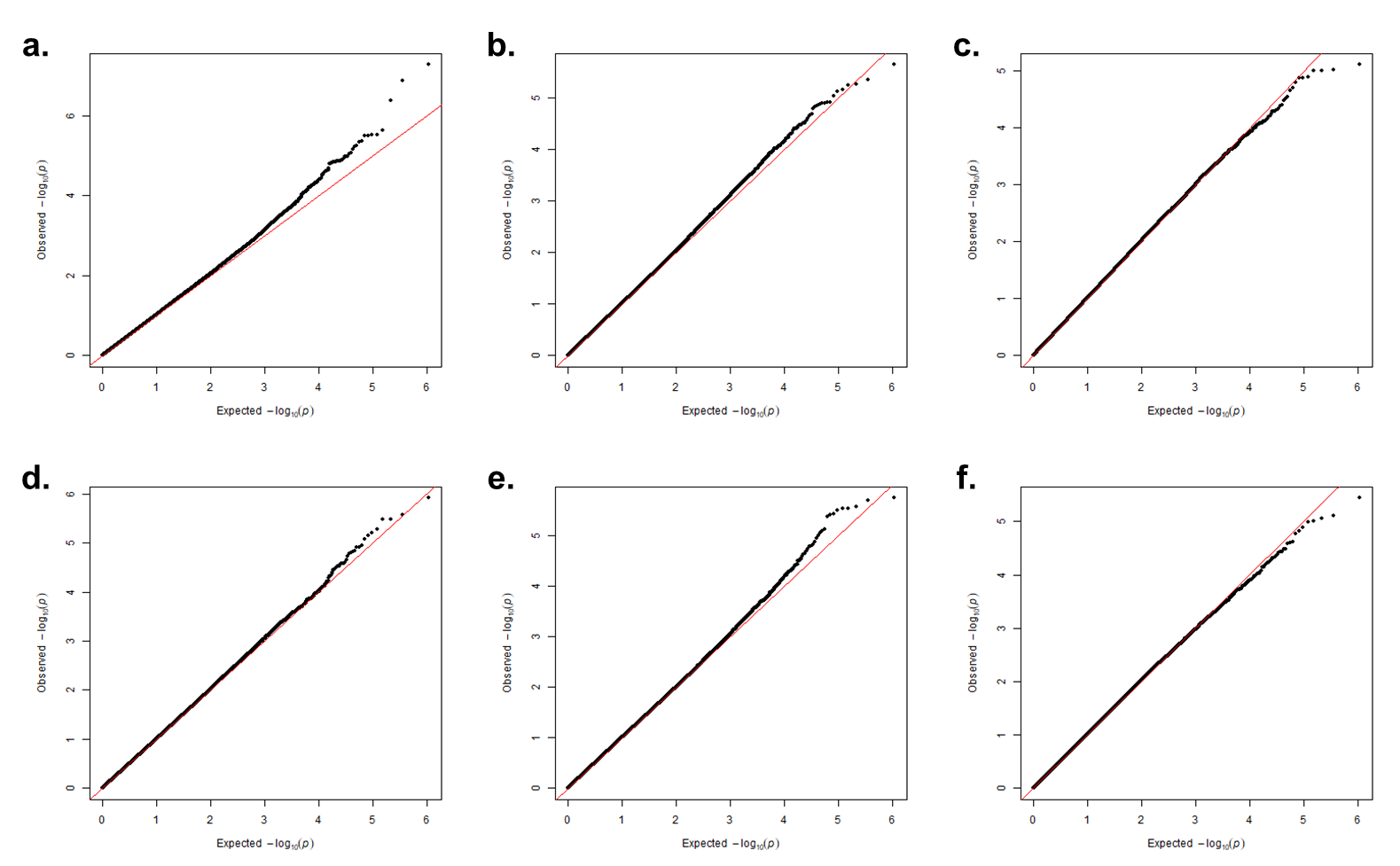
1.00), (e.) ↑W and Uncategorised depression (λ = 1.00), (f.) ↓W and Uncategorised depression (λ = 1.01). The x-axis shows the expected -log_10_(*p*) and the y-axis the observed -log_10_(*p*).

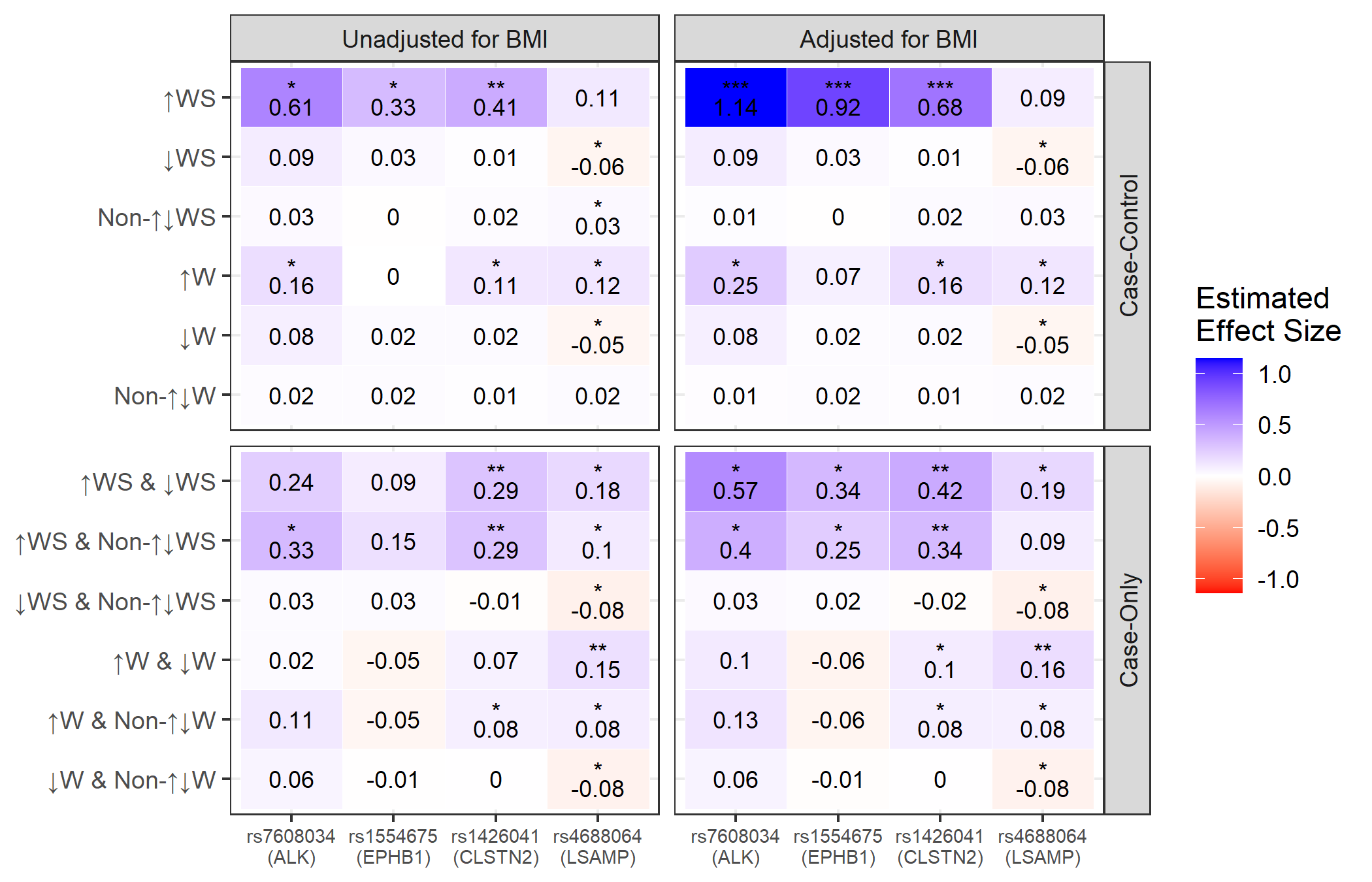
**Supplementary Figure 19**. Estimated effect of *ALK*, *EPHB1*, *CLSTN2* and *LSAMP* loci across all subgroup-specific GWAS

A heat map of the estimated effects of the four SNPs (rs7608034, rs1554675, rs1426041 and rs4688064) nearing or below genome-wide significance in at least one GWAS in each GWAS performed. Significance is denoted as follows: * = nominal significance (*p* < 0.05), ** = suggestive significance (*p* = 1 x 10^-5^) and *** = genome-wide significance (*p* < 5 x 10^-8^).

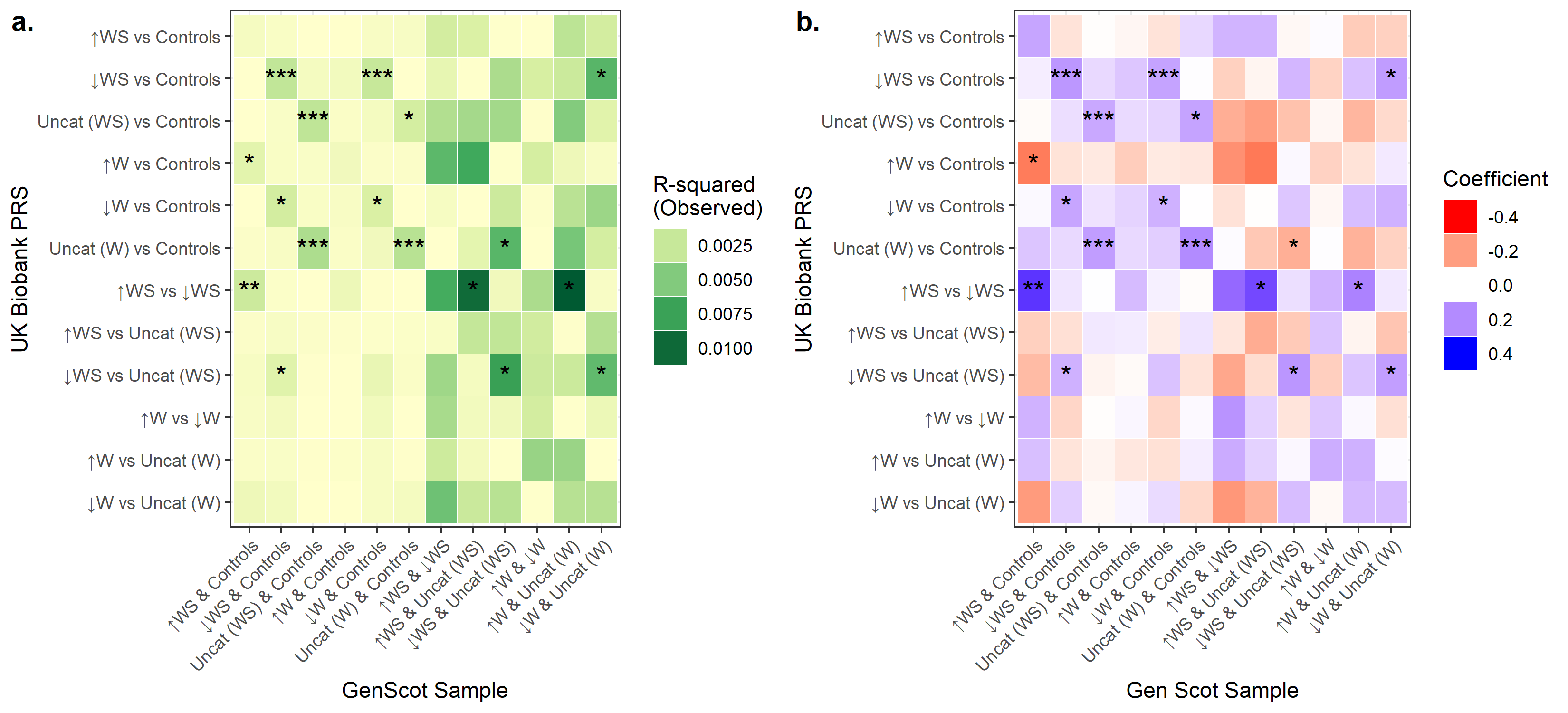
**Supplementary Figure 20**. Results of PRS analyses using UK Biobank subgroup case-control and case-only GWAS (adjusted for BMI).

*Heatmaps showing results from polygenic risk score analyses of depression subgroups using case-control and case-only UK Biobank GWAS adjusted for BMI. (a) shows the observed R-squared values and (b) the coefficient from each analysis. The ‘best’ p-value threshold and number of SNPs in each PRS are shown in supplementary tables 27 & 29. Significance thresholds: * = p < 0.05, ** = p < 0.05/6, *** = p < 0.05/12.*

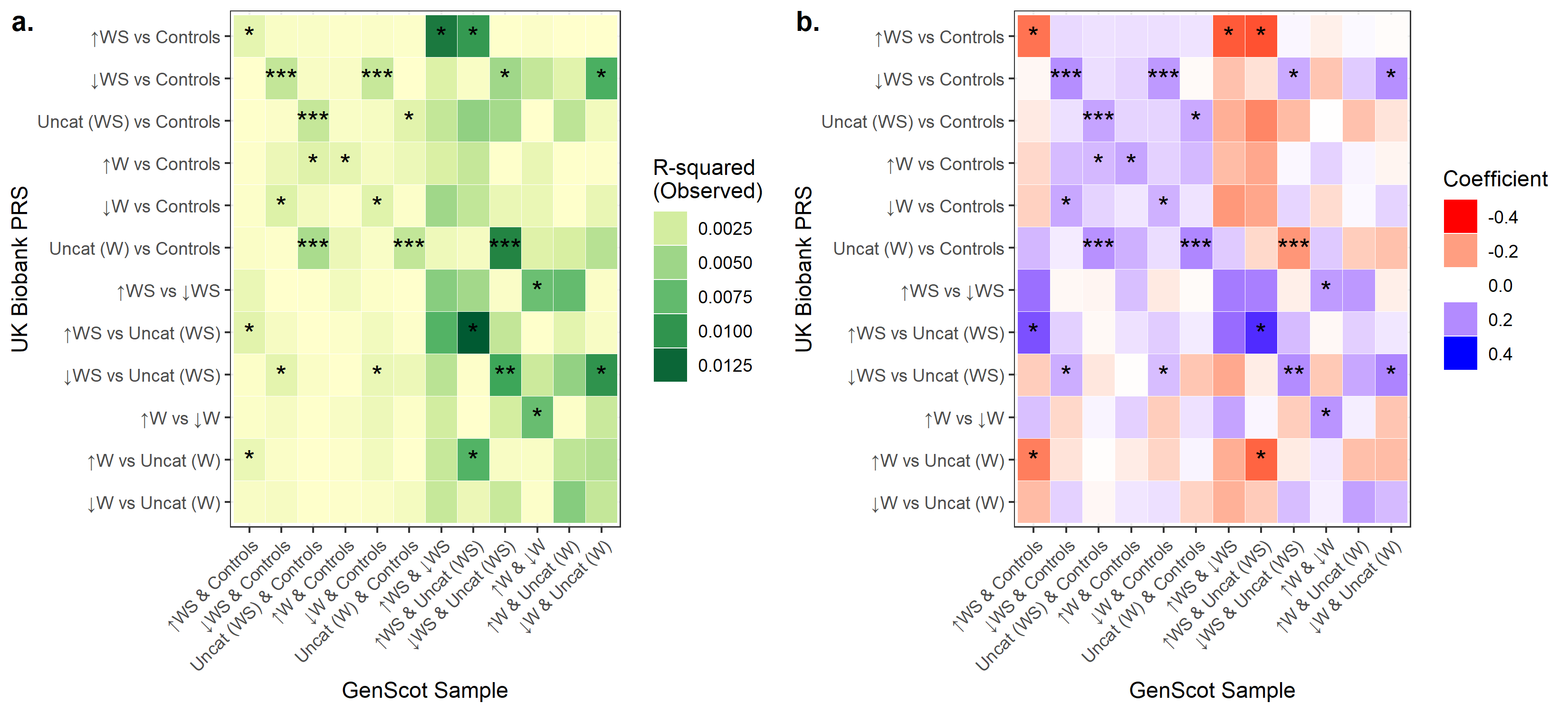
**Supplementary Figure 21**. Results of PRS analyses using UK Biobank subgroup case-control and case-only GWAS (unadjusted for BMI).

*Heatmaps showing results from polygenic risk score analyses of depression subgroups using case-control and case-only UK Biobank GWAS unadjusted for BMI. (a) shows the observed R-squared values and (b) the coefficient from each analysis. The ‘best’ p-value threshold and number of SNPs in each PRS are shown in supplementary tables 28 & 30. Significance thresholds: * = p < 0.05, ** = p < 0.05/6, *** = p < 0.05/12.*

**Supplementary Figure 22.** High resolution plots from PRS analyses in Generation Scotland sample

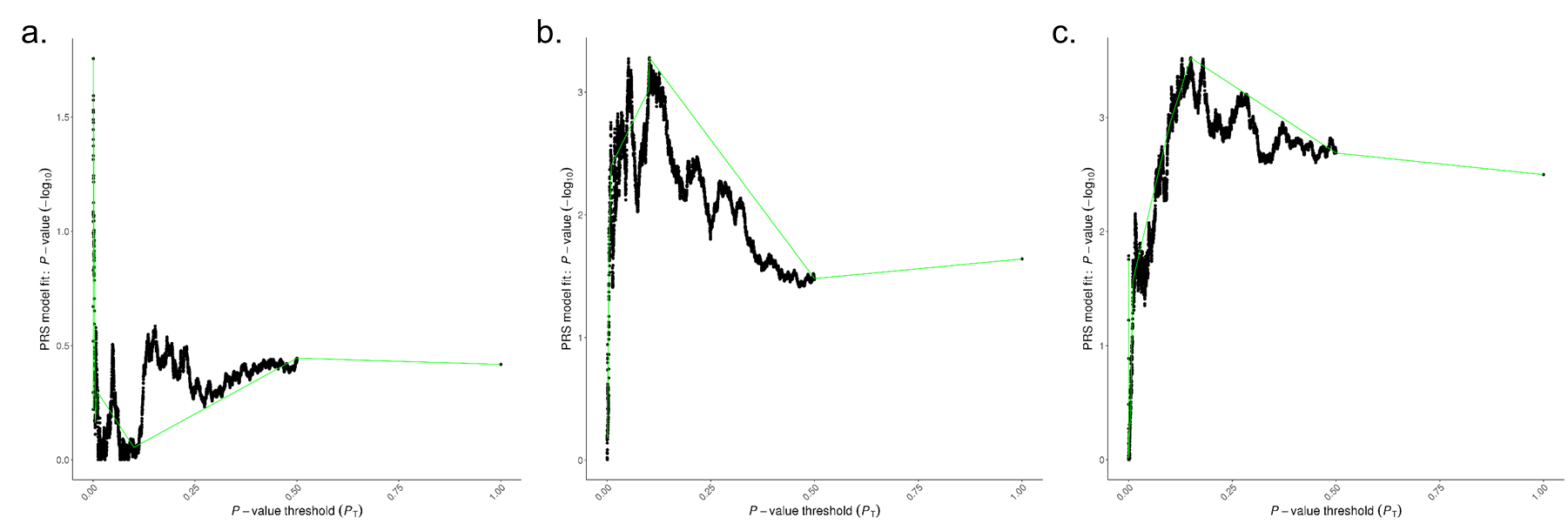

*High resolution plots across a range of p-values for (a.) a BMI adjusted ↑WS depression UKB PRS in a Generation Scotland sample of ↑WS depression cases and controls, (b.) a BMI adjusted ↓WS depression UKB PRS in a Generation Scotland sample of ↓WS depression cases and controls and (c.) a BMI adjusted uncategorised depression UKB PRS in a Generation Scotland sample of uncategorised depression cases and controls.*

**Supplementary Figure 23.** Genetic correlation results for subgroup-MDD and subgroup-subgroup comparisons

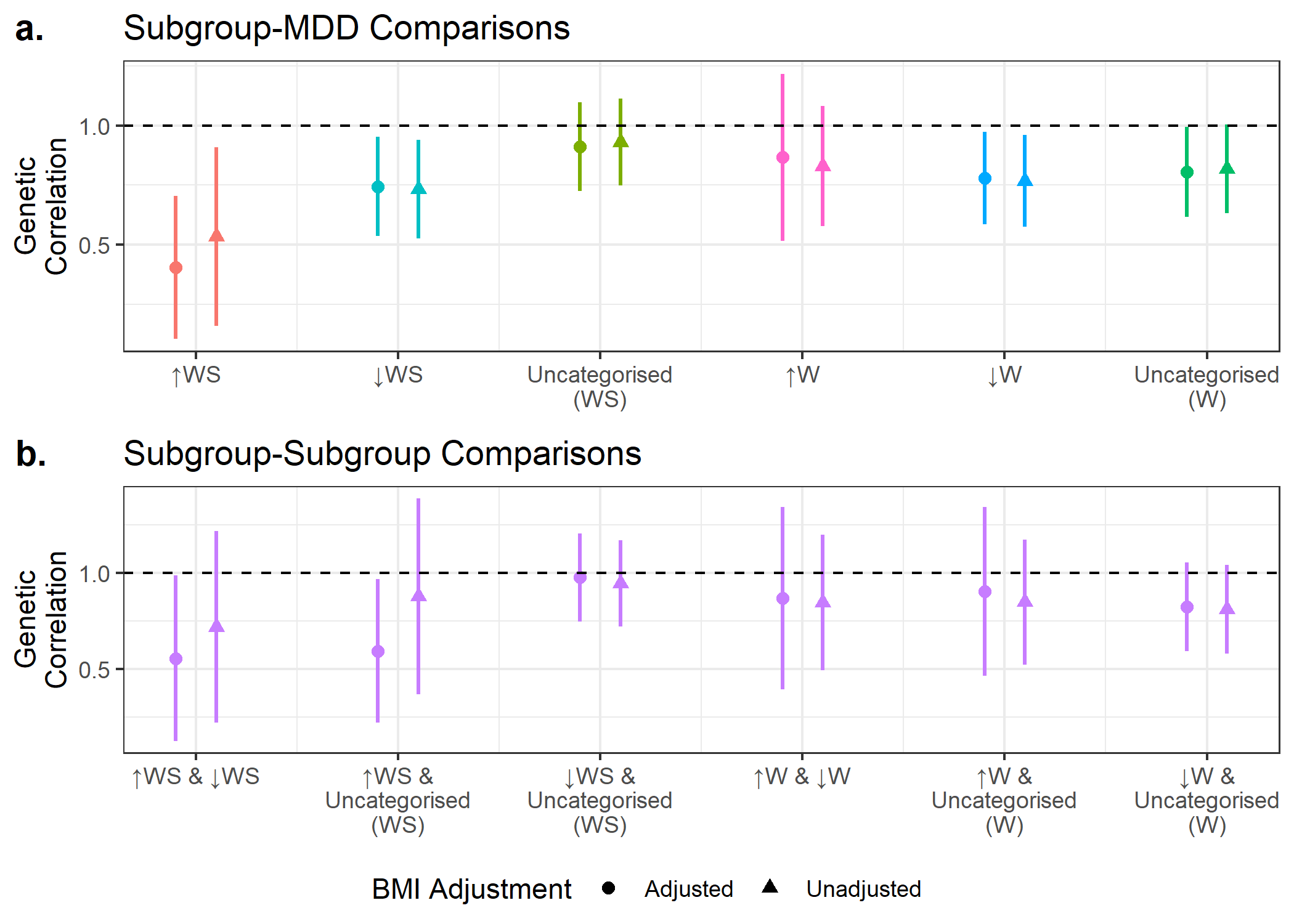

*Forest plots of the genetic correlations and 95% confidence intervals of (a.) subgroup-MDD comparisons and (b.) subgroup-subgroup comparisons. The black dashed line represents a genetic correlation of 1.00. Results adjusted and unadjusted for BMI are shown.*

**Supplementary Table 1**. The sample sizes for each depression subgroup for UK Biobank analyses where some individuals were missing data.

|  | **Lifetime MDD** | **Controls** | **↑WS** | **↓WS** | **Uncategorised (WS)** | **↑W** | **↓W** | **Uncategorised (W)** |
| --- | --- | --- | --- | --- | --- | --- | --- | --- |
| **Full Sample** | 26,662 | 50,147 | 1,525 | 9,067 | 16,070 | 5,015 | 10,924 | 10,723 |
| **BMI** | 26,599 | 50,064 | 1,522 | 9,050 | 16,027 | 5,001 | 10,901 | 10,697 |
| **Age of Onset** | 24,980 | - | 1,410 | 8,659 | 14,911 | 4,619 | 10,432 | 9,929 |
| **Number of Episodes** | 20,573 | - | 918 | 7,569 | 12,086 | 3,406 | 9,093 | 8,074 |
| **Neuroticism Score** | 22,210 | 43,122 | 1,269 | 7,645 | 13,296 | 4,199 | 9,166 | 8,845 |
| **PHQ-9 Score** | 26,662 | 50,147 | 1,525 | 9,067 | 16,070 | 5,015 | 10,924 | 10,723 |
| **GAD-7 Score** | 26,409 | 49,897 | 1,514 | 8,990 | 15,905 | 4,965 | 10,828 | 10,616 |
| **Wellbeing Score** | 25,979 | 48,836 | 1,484 | 8,856 | 15,639 | 4,897 | 10,658 | 10,424 |

**Supplementary Table 2.** Sample sizes for PRS analyses in Generation Scotland

| **Case-control Analysis** | | | **Case-only Analysis** | | |
| --- | --- | --- | --- | --- | --- |
| ***Comparison*** | ***Group 1*** | ***Group 2*** | ***Comparison*** | ***Group 1*** | ***Group 2*** |
| **↑WS & Controls** | 75 | 3085 | **↑WS & ↓WS** | 75 | 366 |
| **↓WS & Controls** | 375 | 2926 | **↑WS & Uncategorised (WS)** | 75 | 451 |
| **Uncategorised (WS) & Controls** | 464 | 2869 | **↓WS & Uncategorised (WS)** | 375 | 431 |
| **↑W & Controls** | 239 | 2994 | **↑W & ↓W** | 239 | 401 |
| **↓W & Controls** | 426 | 2894 | **↑W & Uncategorised (W)** | 239 | 224 |
| **Uncategorised (W) & Controls** | 262 | 2985 | **↓W & Uncategorised (W)** | 401 | 262 |

**Supplementary Table 3**. Disease codes (ICD-10, self-reported non-cancer illness) used to identify cases of disease of interest

|  | **ICD-10 Code(s)** | **Non-Cancer Illness Code** |
| --- | --- | --- |
| Neurological Disorders | | |
| Epilepsy | G40 | 1264 |
| Migraine | G43 | 1265 |
| Multiple Sclerosis | G35 | 1261 |
| Inflammatory Disorders | | |
| Crohn’s Disease | K50 | 1462 |
| Ulcerative Colitis | K51 | 1463 |
| Rheumatoid Arthritis | M06 | 1464 |
| Eczema/Dermatitis | L20-L30 | 1452 |
| Asthma | J45 | 1111 |

**Supplementary Table 4.** The sample and depression characteristics and mental health scores of weight-sleep and weight-only depression subgroups

|  | **↑WS** | **↓WS** | **Uncat (WS)** | **↑W** | **↓W** | **Uncat (W)** |
| --- | --- | --- | --- | --- | --- | --- |
| ***Sample Characteristics*** | | | |  |  |  |
| Age at Initial Assessment (years) | 52.38  (7.10) | 55.11^†^  (7.45) | 54.95^†^  (7.55) | 53.48  (7.25) | 55.14^††^  (7.50) | 55.22^††^  (7.59) |
| Proportion of  Females (%) | 77.11^‡^ | 76.52^‡^ | 65.02 | 74.73^‡‡^ | 76.50^‡‡^ | 60.23 |
| Body Mass Index* (kg/m^2^) | 29.72  (26.36-33.64) | 25.18  (22.88-28.18) | 26.61  (23.98-30.01) | 29.22  (26.15-33.05) | 25.14  (22.82-28.16) | 26.19  (23.75-29.34) |
| ***Depression Characteristics*** | | | |  |  |  |
| Age of Onset (years) | 32.42  (14.58) | 36.38  (14.07) | 35.32  (14.58) | 34.82^§^  (14.73) | 36.05  (14.22) | 35.31^§^  (14.49) |
| Number of Episodes* | 3  (1-4) | 1^‖^  (1-3) | 1^‖^  (1-3) | 2  (1-4) | 1^‖‖^  (1-3) | 1^‖‖^  (1-3) |
| Proportion of Individuals reporting: “Too many to count/one episode ran into the next” (%) | 37.44% | 13.83% | 21.59% | 29.51% | 14.00% | 21.32% |
| ***Mental Health Scores*** | | | |  |  |  |
| Neuroticism | 6.34  (3.26) | 5.26  (3.30) | 5.49  (3.30) | 6.05  (3.25) | 5.24  (3.31) | 5.40  (3.29) |
| PHQ-9* | 6  (3-11) | 3  (1-6) | 3  (1-7) | 5  (2-9) | 2  (1-6) | 3  (1-6) |
| GAD-7* | 4  (0-7) | 2  (0-5) | 2  (0-6) | 4  (0-7) | 2^◊^  (0-5) | 2^◊^  (0-5) |
| Wellbeing | 10.95  (2.45) | 12.25  (2.08) | 11.85  (2.18) | 11.24  (2.36) | 12.23  (2.09) | 11.93  (2.12) |

*For characteristics denoted by * median and interquartile range (IQR) are shown, rather than mean and standard deviation, as data were not normally distributed. Non-significant comparison pairs (p > 0.05) for subgroups defined using weight and sleep changes or weight changes only are highlighted with matching symbols (^†^p = 0.10, ^††^p = 0.46, ^‡^p = 0.63, ^‡‡^p = 0.016, ^§^p = 0.06, ^‖^p = 0.18, ^‖‖^p = 0.76, ^◊^p = 0.16). Uncat = uncategorised.*

**Supplementary Table 5**. Individual measures (mean & standard deviation) of the wellbeing score for each depression subgroup.

|  | ↑WS | ↓WS | Uncat (WS) | ↑W | ↓W | Uncat (W) |
| --- | --- | --- | --- | --- | --- | --- |
| General happiness | 4.08  (0.91) | 4.40  (0.82) | 4.30  (0.84) | 4.14  (0.88) | 4.41  (0.88) | 4.32  (0.83) |
| Happiness with health | 3.65  (1.12) | 4.27  (0.96) | 4.10  (1.02) | 3.79  (1.10) | 4.27  (0.97) | 4.16  (0.99) |
| Life thought to be meaningful | 3.22  (1.00) | 3.58  (0.86) | 3.45  (0.89) | 3.31  (0.95) | 3.57  (0.87) | 3.46  (0.88) |

*Uncat = uncategorised.*

**Supplementary Table 6**. Results from comparisons of sample characteristics between weight-sleep subgroups

|  | **Statistical Test** | **Test Statistic** | **Degrees of Freedom** | ***P*-value** |
| --- | --- | --- | --- | --- |
| ***Age at Initial Assessment*** | | | | |
| ↑WS vs ↓WS | Two Sample T-test | -13.82 | 2128 | ***≤ 2.20 x 10^-16^*** |
| ↑WS vs Uncategorised (WS) | Two Sample T-test | -13.46 | 1866 | ***≤ 2.20 x 10^-16^*** |
| ↓WS vs Uncategorised (WS) | Two Sample T-test | 1.64 | 19003 | 0.102 |
| ***Proportion of Females*** | | | | |
| ↑WS vs ↓WS | Chi-Square Test | 0.23 | 1 | 0.634 |
| ↑WS vs Uncategorised (WS) | Chi-Square Test | 90.32 | 1 | ***≤ 2.20 x 10^-16^*** |
| ↓WS vs Uncategorised (WS) | Chi-Square Test | 358.77 | 1 | ***≤ 2.20 x 10^-16^*** |
| ***Body Mass Index*** | | | | |
| ↑WS vs ↓WS | Wilcoxon Test | 10335714 | - | ***≤ 2.20 x 10^-16^*** |
| ↑WS vs Uncategorised (WS) | Wilcoxon Test | 16216932 | - | ***≤ 2.20 x 10^-16^*** |
| ↓WS vs Uncategorised (WS) | Wilcoxon Test | 58489428 | - | ***≤ 2.20 x 10^-16^*** |

Significant comparisons after correction for multiple testing are shown in **bold** (*p* < 5.56 x 10^-3^), nominally significant comparisons (*p* < 0.05) are *italicised*.

**Supplementary Table 7**. Results from comparisons of depression characteristics between weight-sleep subgroups

|  | **Statistical Test** | **Test Statistic** | **Degrees of Freedom** | ***P*-value** |
| --- | --- | --- | --- | --- |
| ***Age of Onset*** | | | | |
| ↑WS vs ↓WS | Two Sample T-test | -9.51 | 1862 | ***≤ 2.20 x 10^-16^*** |
| ↑WS vs Uncategorised (WS) | Two Sample T-test | -7.16 | 1687 | ***1.17 x 10^-12^*** |
| ↓WS vs Uncategorised (WS) | Two Sample T-test | 5.47 | 18618 | ***4.54 x 10^-8^*** |
| ***Number of Episodes*** | | | | |
| ↑WS vs ↓WS | Wilcoxon Test | 4114884 | - | ***≤ 2.20 x 10^-16^*** |
| ↑WS vs Uncategorised (WS) | Wilcoxon Test | 6494020 | - | ***≤ 2.20 x 10^-16^*** |
| ↓WS vs Uncategorised (WS) | Wilcoxon Test | 45255874 | - | 0.179 |
| ***Proportion reporting: “Too many to count/one episode ran into the next”*** | | | | |
| ↑WS vs ↓WS | Chi-Square Test | 508.70 | 1 | ***≤ 2.20 x 10^-16^*** |
| ↑WS vs Uncategorised (WS) | Chi-Square Test | 196.87 | 1 | ***≤ 2.20 x 10^-16^*** |
| ↓WS vs Uncategorised (WS) | Chi-Square Test | 228.37 | 1 | ***≤ 2.20 x 10^-16^*** |

Significant comparisons after correction for multiple testing are shown in **bold** (*p* < 5.56 x 10^-3^), nominally significant comparisons (*p* < 0.05) are *italicised*.

**Supplementary Table 8**. Results from comparisons of mental health scores between weight-sleep subgroups

|  | **Statistical Test** | **Test Statistic** | **Degrees of Freedom** | ***P*-value** |
| --- | --- | --- | --- | --- |
| ***Neuroticism*** | | | | |
| ↑WS vs ↓WS | Two Sample T-test | 10.89 | 1726 | ***8.98E-27*** |
| ↑WS vs Uncategorised (WS) | Two Sample T-test | 8.91 | 1526 | ***1.44E-18*** |
| ↓WS vs Uncategorised (WS) | Two Sample T-test | -4.74 | 15934 | ***2.17E-06*** |
| ***PHQ-9*** | | | | |
| ↑WS vs ↓WS | Wilcoxon Test | 9503894 | - | ***2.20E-123*** |
| ↑WS vs Uncategorised (WS) | Wilcoxon Test | 15838756 | - | ***1.17E-80*** |
| ↓WS vs Uncategorised (WS) | Wilcoxon Test | 66174002 |  | ***4.12E-34*** |
| ***GAD-7*** | | | | |
| ↑WS vs ↓WS | Wilcoxon Test | 8061984 | - | ***3.79E-32*** |
| ↑WS vs Uncategorised (WS) | Wilcoxon Test | 13971039 | - | ***4.89E-26*** |
| ↓WS vs Uncategorised (WS) | Wilcoxon Test | 69646429 | - | ***5.08E-04*** |
| ***Wellbeing*** | | | | |
| ↑WS vs ↓WS | Two Sample T-test | -19.21 | 1860 | ***3.39E-75*** |
| ↑WS vs Uncategorised (WS) | Two Sample T-test | -13.56 | 1713 | ***6.93E-40*** |
| ↓WS vs Uncategorised (WS) | Two Sample T-test | 14.18 | 19082 | ***2.21E-45*** |

Significant comparisons after correction for multiple testing are shown in **bold** (*p* < 4.17 x 10^-3^), nominally significant comparisons (*p* < 0.05) are *italicised*.

**Supplementary Table 9**. Results from comparisons of individual PHQ-9 symptoms between weight-sleep subgroups

|  | **Chi-square Value** | **Degrees of Freedom** | ***P*-value** |
| --- | --- | --- | --- |
| ***Little interest or pleasure in doing things*** | | | |
| ↑WS vs ↓WS | 167.68 | 1 | ***2.38E-38*** |
| ↑WS vs Uncategorised (WS) | 83.21 | 1 | ***7.37E-20*** |
| ↓WS vs Uncategorised (WS) | 46.49 | 1 | ***9.22E-12*** |
| ***Feeling down, depressed or hopeless*** | | | |
| ↑WS vs ↓WS | 186.38 | 1 | ***1.96E-42*** |
| ↑WS vs Uncategorised (WS) | 89.98 | 1 | ***2.40E-21*** |
| ↓WS vs Uncategorised (WS) | 52.76 | 1 | ***3.78E-13*** |
| ***Trouble falling or staying asleep or sleeping too much*** | | | |
| ↑WS vs ↓WS | 103.4 | 1 | ***2.74E-24*** |
| ↑WS vs Uncategorised (WS) | 73.73 | 1 | ***8.94E-18*** |
| ↓WS vs Uncategorised (WS) | 13.15 | 1 | ***2.87E-04*** |
| ***Feeling tired or having little energy*** | | | |
| ↑WS vs ↓WS | 383.3 | 1 | ***2.38E-85*** |
| ↑WS vs Uncategorised (WS) | 236.85 | 1 | ***1.91E-53*** |
| ↓WS vs Uncategorised (WS) | 69.44 | 1 | ***7.88E-17*** |
| Poor appetite or overeating | | | |
| ↑WS vs ↓WS | 516.89 | 1 | ***2.01E-114*** |
| ↑WS vs Uncategorised (WS) | 351.56 | 1 | ***1.94E-78*** |
| ↓WS vs Uncategorised (WS) | 57.53 | 1 | ***3.33E-14*** |
| ***Feeling bad about yourself, that you are a failure, that you have let yourself or family down*** | | | |
| ↑WS vs ↓WS | 195.76 | 1 | ***1.75E-44*** |
| ↑WS vs Uncategorised (WS) | 107.12 | 1 | ***4.18E-25*** |
| ↓WS vs Uncategorised (WS) | 44.74 | 1 | ***2.25E-11*** |
| ***Trouble concentrating on things*** | | | |
| ↑WS vs ↓WS | 220.92 | 1 | ***5.71E-50*** |
| ↑WS vs Uncategorised (WS) | 171.09 | 1 | ***4.28E-39*** |
| ↓WS vs Uncategorised (WS) | 15.39 | 1 | ***8.77E-05*** |
| ***Noticeably moving or speaking slowly, or more being fidgety or restless than usual*** | | | |
| ↑WS vs ↓WS | 111.55 | 1 | ***4.48E-26*** |
| ↑WS vs Uncategorised (WS) | 77.55 | 1 | ***1.30E-18*** |
| ↓WS vs Uncategorised (WS) | 11.12 | 1 | ***8.54E-04*** |
| ***Thoughts you would be better off dead or hurting yourself*** | | | |
| ↑WS vs ↓WS | 47.31 | 1 | ***6.05E-12*** |
| ↑WS vs Uncategorised (WS) | 23.48 | 1 | ***1.26E-06*** |
| ↓WS vs Uncategorised (WS) | 11.99 | 1 | ***5.36E-04*** |

Significant comparisons after correction for multiple testing are in **bold** (*p* < 1.72 x 10^-3^, 0.05/24), nominally significant comparisons (*p* < 0.05) are *italicised*.

**Supplementary Table 10**. Results from comparisons of sample characteristics between weight-only subgroups

|  | **Statistical Test** | **Test Statistic** | **Degrees of Freedom** | ***P*-value** |
| --- | --- | --- | --- | --- |
| ***Age at Initial Assessment*** | | | | |
| ↑W vs ↓W | Two Sample T-test | -13.35 | 10041 | ***2.20E-16*** |
| ↑W vs Uncategorised (W) | Two Sample T-test | -13.85 | 10217 | ***2.20E-16*** |
| ↓W vs Uncategorised (W) | Two Sample T-test | -0.74 | 21625 | 0.4602 |
| ***Proportion of Females*** | | | | |
| ↑W vs ↓W | Chi-Square Test | 5.77 | 1 | *0.01632* |
| ↑W vs Uncategorised (W) | Chi-Square Test | 314.97 | 1 | ***2.20E-16*** |
| ↓W vs Uncategorised (W) | Chi-Square Test | 662.87 | 1 | ***2.20E-16*** |
| ***Body Mass Index*** | | | | |
| ↑W vs ↓W | Wilcoxon Test | 40371575 | - | ***2.20E-16*** |
| ↑W vs Uncategorised (W) | Wilcoxon Test | 36205316 | - | ***2.20E-16*** |
| ↓W vs Uncategorised (W) | Wilcoxon Test | 49515962 | - | ***2.20E-16*** |

Significant comparisons after correction for multiple testing are shown in **bold** (*p* < 5.56 x 10^-3^), nominally significant comparisons (*p* < 0.05) are *italicised*.

**Supplementary Table 11**. Results from comparisons of depression characteristics between weight-only subgroups

|  | **Statistical Test** | **Test Statistic** | **Degrees of Freedom** | ***P*-value** |
| --- | --- | --- | --- | --- |
| ***Age of Onset*** | | | | |
| ↑W vs ↓W | Two Sample T-test | -4.80 | 8570 | ***1.65E-06*** |
| ↑W vs Uncategorised (W) | Two Sample T-test | -1.89 | 8879 | 0.05917 |
| ↓W vs Uncategorised (W) | Two Sample T-test | 3.69 | 20263 | ***0.0002271*** |
| ***Number of Episodes*** | | | | |
| ↑W vs ↓W | Wilcoxon Test | 17573700 | - | ***2.20E-16*** |
| ↑W vs Uncategorised (W) | Wilcoxon Test | 15529012 | - | ***2.20E-16*** |
| ↓W vs Uncategorised (W) | Wilcoxon Test | 36615788 | - | 0.7574 |
| ***Proportion reporting: “Too many to count/one episode ran into the next”*** | | | | |
| ↑W vs ↓W | Chi-Square Test | 539.22 | 1 | ***2.20E-16*** |
| ↑W vs Uncategorised (W) | Chi-Square Test | 125.55 | 1 | ***2.20E-16*** |
| ↓W vs Uncategorised (W) | Chi-Square Test | 199.32 | 1 | ***2.20E-16*** |

Significant comparisons after correction for multiple testing are shown in **bold** (*p* < 5.56 x 10^-3^), nominally significant comparisons (*p* < 0.05) are *italicised*.

**Supplementary Table 12**. Results from comparisons of mental health scores between weight-only subgroups

|  | **Statistical Test** | **Test Statistic** | **Degrees of Freedom** | ***P*-value** |
| --- | --- | --- | --- | --- |
| ***Neuroticism*** | | | | |
| ↑W vs ↓W | Two Sample T-test | 13.30 | 8274 | ***6.05E-40*** |
| ↑W vs Uncategorised (W) | Two Sample T-test | 10.64 | 8341 | ***2.92E-26*** |
| ↓W vs Uncategorised (W) | Two Sample T-test | -3.24 | 17992 | ***1.19E-03*** |
| ***PHQ-9*** | | | | |
| ↑W vs ↓W | Wilcoxon Test | 35744588 |  | ***3.76E-213*** |
| ↑W vs Uncategorised (W) | Wilcoxon Test | 33643606 |  | ***3.24E-144*** |
| ↓W vs Uncategorised (W) | Wilcoxon Test | 55175046 |  | ***9.71E-14*** |
| ***GAD-7*** | | | | |
| ↑W vs ↓W | Wilcoxon Test | 31481012 |  | ***5.80E-70*** |
| ↑W vs Uncategorised (W) | Wilcoxon Test | 30603834 |  | ***1.02E-61*** |
| ↓W vs Uncategorised (W) | Wilcoxon Test | 56849196 |  | 1.55E-01 |
| ***Wellbeing*** | | | | |
| ↑W vs ↓W | Two Sample T-test | -25.80 | 8558 | ***2.05E-141*** |
| ↑W vs Uncategorised (W) | Two Sample T-test | -17.60 | 8723 | ***3.82E-68*** |
| ↓W vs Uncategorised (W) | Two Sample T-test | 10.97 | 21054 | ***6.68E-28*** |

Significant comparisons after correction for multiple testing are shown in **bold** (*p* < 4.17 x 10^-3^), nominally significant comparisons (*p* < 0.05) are *italicised*.

**Supplementary Table 13**. Results from comparisons of individual PHQ-9 symptoms between weight-only subgroups

|  | **Chi-square Value** | **Degrees of Freedom** | ***P*-value** |
| --- | --- | --- | --- |
| ***Little interest or pleasure in doing things*** | | | |
| ↑W vs ↓W | 234.42 | 1 | ***6.48E-53*** |
| ↑W vs Uncategorised (W) | 111.79 | 1 | ***3.97E-26*** |
| ↓W vs Uncategorised (W) | 29.05 | 1 | ***7.07E-08*** |
| ***Feeling down, depressed or hopeless*** | | | |
| ↑W vs ↓W | 225.05 | 1 | ***7.16E-51*** |
| ↑W vs Uncategorised (W) | 105.62 | 1 | ***8.94E-25*** |
| ↓W vs Uncategorised (W) | 28.84 | 1 | ***7.85E-08*** |
| ***Trouble falling or staying asleep or sleeping too much*** | | | |
| ↑W vs ↓W | 201.14 | 1 | ***1.18E-45*** |
| ↑W vs Uncategorised (W) | 172.79 | 1 | ***1.82E-39*** |
| ↓W vs Uncategorised (W) | 1.38 | 1 | 0.24 |
| ***Feeling tired or having little energy*** | | | |
| ↑W vs ↓W | 435.35 | 1 | ***1.11E-96*** |
| ↑W vs Uncategorised (W) | 268.62 | 1 | ***2.26E-60*** |
| ↓W vs Uncategorised (W) | 27.01 | 1 | ***2.03E-07*** |
| ***Poor appetite or overeating*** | | | |
| ↑W vs ↓W | 779.85 | 1 | ***1.29E-171*** |
| ↑W vs Uncategorised (W) | 759.82 | 1 | ***2.93E-167*** |
| ↓W vs Uncategorised (W) | 0.05 | 1 | 0.818 |
| ***Feeling bad about yourself, that you are a failure, that you have let yourself or family down*** | | | |
| ↑W vs ↓W | 275.33 | 1 | ***7.83E-62*** |
| ↑W vs Uncategorised (W) | 177.94 | 1 | ***1.37E-40*** |
| ↓W vs Uncategorised (W) | 13.46 | 1 | ***2.44E-04*** |
| ***Trouble concentrating on things*** | | | |
| ↑W vs ↓W | 214.49 | 1 | ***1.44E-48*** |
| ↑W vs Uncategorised (W) | 148.17 | 1 | ***4.36E-34*** |
| ↓W vs Uncategorised (W) | 7.64 | 1 | *5.70E-03* |
| ***Noticeably moving or speaking slowly, or more being fidgety or restless than usual*** | | | |
| ↑W vs ↓W | 105.59 | 1 | ***9.05E-25*** |
| ↑W vs Uncategorised (W) | 84.78 | 1 | ***3.33E-20*** |
| ↓W vs Uncategorised (W) | 1.25 | 1 | 2.63E-01 |
| ***Thoughts you would be better off dead or hurting yourself*** | | | |
| ↑W vs ↓W | 46.46 | 1 | ***9.37E-12*** |
| ↑W vs Uncategorised (W) | 21.37 | 1 | ***3.80E-06*** |
| ↓W vs Uncategorised (W) | 5.99 | 1 | *1.44E-02* |

Significant comparisons after correction for multiple testing are in **bold** (*p* < 1.72 x 10^-3^, 0.05/24), nominally significant comparisons (*p* < 0.05) are *italicised*.

**Supplementary Table 14**. Results from comparisons of prevalence of neurological disorders between weight-sleep subgroups

|  | **Chi-square Value** | **Degrees of Freedom** | ***P*-value** |
| --- | --- | --- | --- |
| **Epilepsy** | | | |
| ↑WS vs ↓WS | 14.42 | 1 | ***1.47E-04*** |
| ↑WS vs Uncategorised (WS) | 15.02 | 1 | ***1.06E-04*** |
| ↓WS vs Uncategorised (WS) | 0.04 | 1 | 8.49E-01 |
| ↑WS vs Controls | 30.09 | 1 | ***4.12E-08*** |
| ↓WS vs Controls | 2.46 | 1 | 1.17E-01 |
| Uncategorised (WS) vs Controls | 5.70 | 1 | *1.70E-02* |
| Lifetime MDD vs Controls | 12.61 | 1 | ***3.85E-04*** |
| **Migraine** | | | |
| ↑WS vs ↓WS | 10.75 | 1 | ***1.04E-03*** |
| ↑WS vs Uncategorised (WS) | 8.47 | 1 | ***3.60E-03*** |
| ↓WS vs Uncategorised (WS) | 0.81 | 1 | 3.68E-01 |
| ↑WS vs Controls | 133.95 | 1 | ***5.61E-31*** |
| ↓WS vs Controls | 209.81 | 1 | ***1.51E-47*** |
| Uncategorised (WS) vs Controls | 365.88 | 1 | ***1.48E-81*** |
| Lifetime MDD vs Controls | 484.16 | 1 | ***2.65E-107*** |
| **Multiple Sclerosis** | | | |
| ↑WS vs ↓WS | 0.62 | 1 | 4.29E-01 |
| ↑WS vs Uncategorised (WS) | 0.75 | 1 | 3.88E-01 |
| ↓WS vs Uncategorised (WS) | 0.00 | 1 | 1.00E+00 |
| ↑WS vs Controls | - | - | 1.03E-01 |
| ↓WS vs Controls | 1.58 | 1 | 2.09E-01 |
| Uncategorised (WS) vs Controls | 2.38 | 1 | 1.23E-01 |
| Lifetime MDD vs Controls | 4.41 | 1 | *3.57E-02* |
| **Parental Dementia** | | | |
| ↑WS vs ↓WS | 1.47 | 1 | 2.26E-01 |
| ↑WS vs Uncategorised (WS) | 5.39 | 1 | *2.03E-02* |
| ↓WS vs Uncategorised (WS) | 5.06 | 1 | *2.45E-02* |
| ↑WS vs Controls | 2.48 | 1 | 1.15E-01 |
| ↓WS vs Controls | 0.44 | 1 | 5.08E-01 |
| Uncategorised (WS) vs Controls | 5.98 | 1 | *1.44E-02* |
| Lifetime MDD vs Controls | 1.22 | 1 | 2.69E-01 |

Significant comparisons after correction for multiple testing are shown in **bold** (*p* < 8.33 x 10^-3^), nominally significant comparisons (*p* < 0.05) are *italicised*.

**Supplementary Table 15**. Results from comparisons of prevalence of inflammatory disorders between weight-sleep subgroups

|  | **Test Statistic** | **Degrees of Freedom** | ***P*-value** |
| --- | --- | --- | --- |
| ***Crohn’s Disease*** | | | |
| ↑WS vs ↓WS | 0.00 | 1 | 1.00E+00 |
| ↑WS vs Uncategorised (WS) | 0.00 | 1 | 1.00E+00 |
| ↓WS vs Uncategorised (WS) | 0.27 | 1 | 6.06E-01 |
| ↑WS vs Controls | 0.04 | 1 | 8.33E-01 |
| ↓WS vs Controls | 1.85 | 1 | 1.73E-01 |
| Uncategorised (WS) vs Controls | 0.67 | 1 | 4.15E-01 |
| Lifetime MDD vs Controls | 1.92 | 1 | 1.65E-01 |
| ***Ulcerative Colitis*** | | | |
| ↑WS vs ↓WS | 0.03 | 1 | 8.60E-01 |
| ↑WS vs Uncategorised (WS) | 0.00 | 1 | 1.00E+00 |
| ↓WS vs Uncategorised (WS) | 0.13 | 1 | 7.16E-01 |
| ↑WS vs Controls | 0.09 | 1 | 7.59E-01 |
| ↓WS vs Controls | 3.26 | 1 | 7.09E-02 |
| Uncategorised (WS) vs Controls | 2.64 | 1 | 1.04E-01 |
| Lifetime MDD vs Controls | 1.92 | 1 | 1.65E-01 |
| ***Rheumatoid Arthritis*** | | | |
| ↑WS vs ↓WS | 0.34 | 1 | 5.58E-01 |
| ↑WS vs Uncategorised (WS) | 0.22 | 1 | 6.37E-01 |
| ↓WS vs Uncategorised (WS) | 0.06 | 1 | 8.09E-01 |
| ↑WS vs Controls | 0.79 | 1 | 3.74E-01 |
| ↓WS vs Controls | 17.47 | 1 | ***2.91E-05*** |
| Uncategorised (WS) vs Controls | 22.22 | 1 | ***2.43E-06*** |
| Lifetime MDD vs Controls | 31.34 | 1 | ***2.17E-08*** |
| ***Asthma*** | | | |
| ↑WS vs ↓WS | 19.44 | 1 | ***1.04E-05*** |
| ↑WS vs Uncategorised (WS) | 13.71 | 1 | ***2.13E-04*** |
| ↓WS vs Uncategorised (WS) | 2.69 | 1 | 1.01E-01 |
| ↑WS vs Controls | 124.86 | 1 | ***5.47E-29*** |
| ↓WS vs Controls | 160.14 | 1 | ***1.06E-36*** |
| Uncategorised (WS) vs Controls | 334.24 | 1 | ***1.15E-74*** |
| Lifetime MDD vs Controls | 450.43 | 1 | ***5.82E-100*** |
| ***Eczema*** | | | |
| ↑WS vs ↓WS | 2.27 | 1 | 1.32E-01 |
| ↑WS vs Uncategorised (WS) | 0.64 | 1 | 4.22E-01 |
| ↓WS vs Uncategorised (WS) | 2.06 | 1 | 1.51E-01 |
| ↑WS vs Controls | 9.41 | 1 | ***2.16E-03*** |
| ↓WS vs Controls | 8.49 | 1 | ***3.56E-03*** |
| Uncategorised (WS) vs Controls | 34.90 | 1 | ***3.47E-09*** |
| Lifetime MDD vs Controls | 39.38 | 1 | ***3.49E-10*** |

Significant comparisons after correction for multiple testing are shown in **bold** (*p* < 8.33 x 10^-3^), nominally significant comparisons (*p* < 0.05) are *italicised*.

**Supplementary Table 16**. Results from comparisons of prevalence of neurological disorders between weight-only subgroups

|  | **Chi-square Value** | **Degrees of Freedom** | ***P*-value** |
| --- | --- | --- | --- |
| ***Epilepsy*** | | | |
| ↑W vs ↓W | 10.74 | 1 | ***1.05E-03*** |
| ↑W vs Uncategorised (W) | 8.04 | 1 | ***4.58E-03*** |
| ↓W vs Uncategorised (W) | 0.18 | 1 | 6.69E-01 |
| ↑W vs Controls | 26.74 | 1 | ***2.33E-07*** |
| ↓W vs Controls | 1.32 | 1 | 2.51E-01 |
| Uncategorised (W) vs Controls | 3.31 | 1 | 6.87E-02 |
| Lifetime MDD vs Controls | 12.61 | 1 | ***3.85E-04*** |
| ***Migraine*** | | | |
| ↑W vs ↓W | 11.00 | 1 | ***9.13E-04*** |
| ↑W vs Uncategorised (W) | 10.93 | 1 | ***9.46E-04*** |
| ↓W vs Uncategorised (W) | 0.00 | 1 | 1.00E+00 |
| ↑W vs Controls | 277.43 | 1 | ***2.73E-62*** |
| ↓W vs Controls | 246.03 | 1 | ***1.91E-55*** |
| Uncategorised (W) vs Controls | 242.76 | 1 | ***9.83E-55*** |
| Lifetime MDD vs Controls | 484.16 | 1 | ***2.65E-107*** |
| ***Multiple Sclerosis*** | | | |
| ↑W vs ↓W | 1.12 | 1 | 2.90E-01 |
| ↑W vs Uncategorised (W) | 3.28 | 1 | 7.02E-02 |
| ↓W vs Uncategorised (W) | 0.63 | 1 | 4.27E-01 |
| ↑W vs Controls | 7.00 | 1 | ***8.17E-03*** |
| ↓W vs Controls | 2.48 | 1 | 1.15E-01 |
| Uncategorised (W) vs Controls | 0.11 | 1 | 7.37E-01 |
| Lifetime MDD vs Controls | 4.41 | 1 | *3.57E-02* |
| ***Parental Dementia*** | | | |
| ↑W vs ↓W | 0.19 | 1 | 6.65E-01 |
| ↑W vs Uncategorised (W) | 5.25 | 1 | *2.20E-02* |
| ↓W vs Uncategorised (W) | 5.48 | 1 | *1.92E-02* |
| ↑W vs Controls | 0.51 | 1 | 4.75E-01 |
| ↓W vs Controls | 0.08 | 1 | 7.79E-01 |
| Uncategorised (W) vs Controls | 7.46 | 1 | ***6.30E-03*** |
| Lifetime MDD vs Controls | 1.22 | 1 | 2.69E-01 |

Significant comparisons after correction for multiple testing are shown in **bold** (*p* < 8.33 x 10^-3^), nominally significant comparisons (*p* < 0.05) are *italicised*.

**Supplementary Table 17**. Results from comparisons of prevalence of inflammatory disorders between weight-only subgroups

|  | **Test Statistic** | **Degrees of Freedom** | ***P*-value** |
| --- | --- | --- | --- |
| ***Crohn’s Disease*** | | | |
| ↑W vs ↓W | 0.25 | 1 | 6.20E-01 |
| ↑W vs Uncategorised (W) | 1.04 | 1 | 3.08E-01 |
| ↓W vs Uncategorised (W) | 0.34 | 1 | 5.58E-01 |
| ↑W vs Controls | 0.00 | 1 | 9.91E-01 |
| ↓W vs Controls | 0.61 | 1 | 4.35E-01 |
| Uncategorised (W) vs Controls | 2.91 | 1 | 8.78E-02 |
| Lifetime MDD vs Controls | 1.92 | 1 | 1.65E-01 |
| ***Ulcerative Colitis*** | | | |
| ↑W vs ↓W | 0.47 | 1 | 4.94E-01 |
| ↑W vs Uncategorised (W) | 1.76 | 1 | 1.84E-01 |
| ↓W vs Uncategorised (W) | 0.59 | 1 | 4.42E-01 |
| ↑W vs Controls | 0.00 | 1 | 1.00E+00 |
| ↓W vs Controls | 1.78 | 1 | 1.82E-01 |
| Uncategorised (W) vs Controls | 6.11 | 1 | *1.35E-02* |
| Lifetime MDD vs Controls | 4.76 | 1 | *2.91E-02* |
| ***Rheumatoid Arthritis*** | | | |
| ↑W vs ↓W | 0.00 | 1 | 9.44E-01 |
| ↑W vs Uncategorised (W) | 0.00 | 1 | 9.49E-01 |
| ↓W vs Uncategorised (W) | 0.09 | 1 | 7.70E-01 |
| ↑W vs Controls | 8.77 | 1 | ***3.06E-03*** |
| ↓W vs Controls | 19.21 | 1 | ***1.17E-05*** |
| Uncategorised (W) vs Controls | 14.85 | 1 | ***1.17E-04*** |
| Lifetime MDD vs Controls | 31.34 | 1 | ***2.17E-08*** |
| ***Asthma*** | | | |
| ↑W vs ↓W | 35.18 | 1 | ***3.01E-09*** |
| ↑W vs Uncategorised (W) | 26.47 | 1 | ***2.68E-07*** |
| ↓W vs Uncategorised (W) | 0.84 | 1 | 3.58E-01 |
| ↑W vs Controls | 301.71 | 1 | ***1.40E-67*** |
| ↓W vs Controls | 173.54 | 1 | ***1.25E-39*** |
| Uncategorised (W) vs Controls | 207.12 | 1 | ***5.83E-47*** |
| Lifetime MDD vs Controls | 450.43 | 1 | ***5.82E-100*** |
| ***Eczema*** | | | |
| ↑W vs ↓W | 7.53 | 1 | ***6.07E-03*** |
| ↑W vs Uncategorised (W) | 7.15 | 1 | ***7.48E-03*** |
| ↓W vs Uncategorised (W) | 0.00 | 1 | 9.70E-01 |
| ↑W vs Controls | 37.43 | 1 | ***9.48E-10*** |
| ↓W vs Controls | 13.11 | 1 | ***2.94E-04*** |
| Uncategorised (W) vs Controls | 13.63 | 1 | ***2.23E-04*** |
| Lifetime MDD vs Controls | 39.38 | 1 | ***3.49E-10*** |

Significant comparisons after correction for multiple testing are shown in **bold** (*p* < 8.33 x 10^-3^), nominally significant comparisons (*p* < 0.05) are *italicised*.

**Supplementary Table 18**. Lead SNPs of suggestive significance (1.00 x 10^-5^) from subgroup case-control GWAS (adjusted for BMI)

|  | **rsID** | **Chr** | **BP** | **EA** | **EAF** | **Beta** | **SE** | **P** | **Nearest Gene** |
| --- | --- | --- | --- | --- | --- | --- | --- | --- | --- |
| ↑ Weight & Sleep | rs2254418 | 1 | 95427877 | A | 0.04 | 0.84 | 0.17 | 5.02E-07 | CNN3 |
|  | rs17431227 | 2 | 14901673 | A | 0.07 | 0.59 | 0.13 | 7.66E-06 | DDX1 |
|  | rs3731997 | 2 | 15496708 | T | 0.05 | 0.76 | 0.16 | 9.16E-07 | **NBAS** |
|  | **rs7608034** | **2** | **29868259** | **G** | **0.03** | **1.14** | **0.21** | **3.09E-08** | **ALK** |
|  | rs3811549 | 2 | 105153544 | T | 0.02 | 1.18 | 0.22 | 1.26E-07 | FHL2 |
|  | rs16842710 | 2 | 159037050 | G | 0.06 | 0.64 | 0.14 | 4.88E-06 | **CCDC148** |
|  | rs6809108 | 3 | 70044334 | T | 0.08 | 0.62 | 0.13 | 1.29E-06 | MITF |
|  | **rs1554675** | **3** | **134826917** | **C** | **0.06** | **0.92** | **0.15** | **8.29E-10** | **EPHB1** |
|  | **rs1426041** | **3** | **139697073** | **A** | **0.13** | **0.68** | **0.10** | **5.12E-11** | **CLSTN2** |
|  | rs12631069 | 3 | 191715361 | T | 0.02 | 1.06 | 0.23 | 2.89E-06 | FGF12 |
|  | rs28409430 | 4 | 93710269 | A | 0.16 | 0.45 | 0.09 | 1.27E-06 | **GRID2** |
|  | rs3797225 | 5 | 53592972 | T | 0.03 | 0.99 | 0.22 | 5.46E-06 | **ARL15** |
|  | rs12654812 | 5 | 176794191 | A | 0.34 | 0.36 | 0.07 | 7.88E-07 | **RGS14** |
|  | rs11154812 | 6 | 99836954 | A | 0.12 | 0.46 | 0.10 | 8.20E-06 | **COQ3** |
|  | rs17339094 | 7 | 72766960 | T | 0.03 | 0.98 | 0.20 | 8.54E-07 | **FKBP6** |
|  | rs573980 | 8 | 20836701 | A | 0.21 | 0.39 | 0.09 | 8.22E-06 |  |
|  | rs6983862 | 8 | 131550333 | T | 0.19 | 0.39 | 0.09 | 9.03E-06 | ASAP1 |
|  | rs348445 | 9 | 75568803 | A | 0.03 | 0.93 | 0.20 | 3.34E-06 | **ALDH1A1** |
|  | rs1999462 | 9 | 77813584 | T | 0.02 | 1.24 | 0.24 | 4.61E-07 |  |
|  | rs7048770 | 9 | 109538962 | G | 0.13 | 0.46 | 0.10 | 9.35E-06 |  |
|  | rs4838320 | 9 | 128831097 | T | 0.08 | 0.60 | 0.12 | 1.62E-06 |  |
|  | rs914976 | 9 | 133229311 | T | 0.08 | 0.58 | 0.13 | 7.58E-06 | HMCN2 |
|  | rs17200443 | 10 | 19765768 | A | 0.06 | 0.68 | 0.15 | 3.77E-06 | MALRD1 |
|  | rs12573809 | 10 | 85464025 | T | 0.05 | 0.73 | 0.16 | 4.86E-06 |  |
|  | rs728748 | 11 | 35894669 | C | 0.01 | 1.73 | 0.34 | 3.86E-07 |  |
|  | rs17096555 | 14 | 30816797 | C | 0.02 | 1.29 | 0.26 | 6.27E-07 |  |
|  | rs11637553 | 15 | 61512506 | C | 0.04 | 0.82 | 0.17 | 1.83E-06 | **RORA** |
|  | rs721186 | 19 | 10265312 | T | 0.01 | 1.45 | 0.29 | 6.46E-07 | **DNMT1** |
|  | rs856323 | 20 | 51325717 | A | 0.02 | 1.17 | 0.26 | 7.20E-06 |  |
|  | rs6121791 | 20 | 60309969 | A | 0.18 | 0.39 | 0.09 | 8.49E-06 | **CDH4** |
|  | rs2824977 | 21 | 20028118 | A | 0.01 | 1.65 | 0.34 | 1.01E-06 |  |
|  | rs12627037 | 21 | 21738573 | A | 0.18 | 0.43 | 0.09 | 1.17E-06 |  |
| ↓ Weight & Sleep | rs7519409 | 1 | 109399660 | G | 0.28 | 0.10 | 0.02 | 8.74E-07 | **AKNAD1** |
|  | rs7567744 | 2 | 55696124 | A | 0.17 | 0.12 | 0.03 | 1.06E-06 | CCDC88A |
|  | rs6786909 | 3 | 850922 | C | 0.34 | -0.10 | 0.02 | 1.80E-07 |  |
|  | rs4473877 | 6 | 144780581 | C | 0.02 | 0.31 | 0.07 | 2.86E-06 | **UTRN** |
|  | rs10271820 | 7 | 118777401 | A | 0.50 | 0.09 | 0.02 | 3.61E-06 |  |
| Uncategorised (Weight-Sleep) | rs12747042 | 1 | 245109861 | C | 0.13 | -0.09 | 0.02 | 7.32E-06 | EFCAB2 |
|  | rs2568871 | 3 | 196692717 | T | 0.39 | -0.06 | 0.01 | 9.86E-06 | **PIGZ** |
|  | rs8180202 | 4 | 93198845 | G | 0.08 | 0.11 | 0.02 | 4.69E-06 |  |
|  | rs7771598 | 6 | 27389258 | A | 0.38 | -0.06 | 0.01 | 9.72E-06 | ZNF391 |
|  | rs10995147 | 10 | 64165285 | A | 0.35 | 0.07 | 0.01 | 3.82E-06 | **ZNF365** |
|  | rs10761940 | 10 | 67060807 | A | 0.37 | -0.08 | 0.01 | 1.02E-07 |  |
|  | rs8022657 | 14 | 67390499 | T | 0.11 | -0.10 | 0.02 | 6.34E-06 | **GPHN** |
|  | rs11084813 | 19 | 35923858 | C | 0.26 | 0.07 | 0.02 | 6.29E-06 | FFAR2 |
| ↑ Weight | rs9792944 | 1 | 32419044 | T | 0.10 | 0.23 | 0.05 | 3.21E-06 | PTP4A2 |
|  | rs4700135 | 5 | 65891203 | T | 0.46 | 0.14 | 0.03 | 1.55E-06 | MAST4 |
|  | rs12517068 | 5 | 142096793 | G | 0.02 | 0.48 | 0.10 | 4.58E-06 | FGF1 |
|  | rs10216044 | 7 | 41812483 | T | 0.40 | 0.14 | 0.03 | 6.98E-06 |  |
|  | rs2225965 | 9 | 23074432 | G | 0.37 | 0.14 | 0.03 | 8.24E-06 |  |
|  | rs871449 | 9 | 78687998 | G | 0.18 | 0.17 | 0.04 | 9.95E-06 | **PCSK5** |
|  | rs12442719 | 15 | 25966038 | C | 0.02 | 0.60 | 0.12 | 5.35E-07 | **ATP10A** |
|  | rs6132939 | 20 | 26239964 | A | 0.34 | -0.15 | 0.03 | 4.07E-06 | FAM182B |
| ↓ Weight | rs839859 | 1 | 109470185 | C | 0.28 | 0.09 | 0.02 | 1.14E-06 | **GPSM2, AKNAD1** |
|  | rs4233750 | 2 | 30102650 | T | 0.21 | 0.10 | 0.02 | 5.22E-06 | **ALK** |
|  | rs7567744 | 2 | 55696124 | A | 0.17 | 0.11 | 0.02 | 4.60E-06 | CCDC88A |
|  | rs6786909 | 3 | 850922 | C | 0.34 | -0.09 | 0.02 | 5.45E-07 |  |
|  | rs576744 | 7 | 78479740 | T | 0.30 | 0.08 | 0.02 | 7.35E-06 | **MAGI2** |
|  | rs2022929 | 8 | 110630465 | G | 0.08 | 0.14 | 0.03 | 5.83E-06 | **SYBU** |
|  | rs2219684 | 14 | 47136591 | A | 0.44 | -0.08 | 0.02 | 9.22E-06 | RPL10L |
|  | rs590076 | 18 | 53260732 | A | 0.34 | 0.08 | 0.02 | 8.51E-06 | **TCF4** |
|  | rs3744927 | 18 | 60245247 | A | 0.23 | -0.10 | 0.02 | 2.77E-06 | **ZCCHC2** |
| Uncategorised (Weight-only) | rs17019186 | 4 | 93113274 | G | 0.07 | 0.14 | 0.03 | 9.44E-06 |  |
|  | rs4735916 | 8 | 677305 | A | 0.09 | -0.13 | 0.03 | 7.02E-06 | **ERICH1** |
|  | rs10995147 | 10 | 64165285 | A | 0.35 | 0.08 | 0.02 | 2.23E-06 | **ZNF365** |
|  | rs8022657 | 14 | 67390499 | T | 0.11 | -0.11 | 0.03 | 9.41E-06 | **GPHN** |
|  | rs3852787 | 16 | 72225733 | G | 0.50 | 0.08 | 0.02 | 1.22E-06 | PMFBP1 |
|  | rs11084813 | 19 | 35923858 | C | 0.26 | 0.09 | 0.02 | 9.44E-07 | FFAR2 |

*Chr = chromosome, BP = base pair, EA = effect allele, EAF = effect allele frequency, Beta = estimated effect, SE = estimated effect standard error, P = p-value. Genes encompassing loci are highlighted in bold. Genome-wide significant SNPs (5.00 x 10^-8^) are highlighted in orange.*

**Supplementary Table 19**. Lead SNPs of suggestive significance (1.00 x 10^-5^) from subgroup case-only GWAS (adjusted for BMI)

|  | **rsID** | **Chr** | **BP** | **EA** | **EAF** | **Beta** | **SE** | **P** | **Nearest Gene** |
| --- | --- | --- | --- | --- | --- | --- | --- | --- | --- |
| ↑ Weight & Sleep  vs  ↓ Weight & Sleep | rs2010753 | 2 | 14891882 | G | 0.4351 | -0.2407 | 0.05364 | 7.28E-06 | DDX1 |
|  | rs2204128 | 3 | 61688122 | G | 0.05874 | 0.4943 | 0.1112 | 8.81E-06 | **PTPRG** |
|  | rs1426041 | 3 | 139697073 | A | 0.1345 | 0.4154 | 0.07745 | 8.33E-08 | **CLSTN2** |
|  | rs10937320 | 3 | 187468973 | G | 0.1088 | 0.3801 | 0.08379 | 5.79E-06 | BCL6 |
|  | rs16992175 | 4 | 36366711 | C | 0.01102 | 1.123 | 0.2512 | 7.97E-06 | DTHD1 |
|  | rs6912049 | 6 | 70380290 | T | 0.01036 | 1.196 | 0.2565 | 3.19E-06 | LMBRD1 |
|  | rs7177677 | 15 | 23065667 | A | 0.01551 | 1.023 | 0.2121 | 1.43E-06 | **NIPA1** |
|  | rs11643923 | 16 | 80264072 | G | 0.04436 | 0.5743 | 0.1275 | 6.70E-06 |  |
|  | rs2595582 | 20 | 3630107 | A | 0.4001 | -0.2482 | 0.05379 | 3.97E-06 | ATRN |
| ↑ Weight & Sleep  vs  Uncategorised (Weight-Sleep) | rs28903985 | 2 | 234959167 | A | 0.01741 | 0.7687 | 0.1691 | 5.54E-06 | SPP2 |
|  | rs3762793 | 3 | 108481730 | C | 0.3902 | -0.2059 | 0.04551 | 6.10E-06 | RETNLB |
|  | rs1426041 | 3 | 139697073 | A | 0.1341 | 0.3363 | 0.06586 | 3.31E-07 | **CLSTN2** |
|  | rs2410633 | 8 | 20060856 | T | 0.2719 | -0.2501 | 0.05006 | 5.93E-07 | **ATP6V1B2** |
|  | rs3809251 | 12 | 3948718 | G | 0.04456 | 0.5007 | 0.1076 | 3.27E-06 | **PARP11** |
|  | rs9319941 | 18 | 56774313 | C | 0.1945 | 0.2602 | 0.05502 | 2.27E-06 | SEC11C |
| ↓ Weight & Sleep  vs  Uncategorised (Weight-Sleep) | rs4647992 | 4 | 103455347 | T | 0.04393 | -0.2285 | 0.04745 | 1.49E-06 | **NFKB1** |
|  | rs10498821 | 6 | 63327975 | A | 0.07637 | -0.1679 | 0.0365 | 4.27E-06 | KHDRBS2 |
|  | rs12006019 | 9 | 4215542 | T | 0.1169 | -0.1365 | 0.03031 | 6.72E-06 | **GLIS3** |
|  | rs911901 | 20 | 55952187 | A | 0.4028 | 0.08932 | 0.01985 | 6.84E-06 | **RAE1** |
| ↑ Weight  vs  ↓ Weight | rs4688064 | 3 | 117551526 | T | 0.2753 | 0.1628 | 0.03353 | 1.22E-06 | **LSAMP** |
|  | rs1012173 | 5 | 149713921 | C | 0.2629 | -0.161 | 0.03431 | 2.71E-06 | **ARSI** |
|  | rs17109539 | 10 | 100131458 | A | 0.03092 | -0.3917 | 0.08722 | 7.15E-06 | PYROXD2 |
|  | rs1834175 | 10 | 109677767 | C | 0.2075 | 0.1684 | 0.03699 | 5.33E-06 |  |
|  | rs17260832 | 18 | 68473541 | G | 0.178 | -0.1749 | 0.03921 | 8.24E-06 | GTSCR1 |
| ↑ Weight  vs  Uncategorised (Weight-only) | rs7564361 | 2 | 207859422 | C | 0.1944 | -0.1509 | 0.03369 | 7.60E-06 | KLF7 |
|  | rs160600 | 6 | 105491263 | A | 0.3772 | 0.1299 | 0.02766 | 2.66E-06 | **LIN28B** |
|  | rs17772576 | 9 | 24821079 | C | 0.2423 | -0.1481 | 0.03102 | 1.82E-06 |  |
|  | rs11027160 | 11 | 23363924 | T | 0.2996 | -0.1354 | 0.02924 | 3.67E-06 | SLC17A6 |
|  | rs17043415 | 12 | 77743656 | A | 0.1783 | -0.1618 | 0.03514 | 4.17E-06 |  |
|  | rs8013446 | 14 | 74359494 | T | 0.1207 | 0.1919 | 0.04104 | 2.96E-06 | **RP5-1021I20.4,  ZNF410** |
|  | rs1595296 | 16 | 59022389 | A | 0.4882 | -0.1276 | 0.02684 | 2.01E-06 |  |
| ↓ Weight  vs  Uncategorised (Weight-only) | rs1559497 | 2 | 31118058 | G | 0.4736 | -0.0885 | 0.02002 | 9.85E-06 | CAPN13 |
|  | rs1512522 | 3 | 110349192 | A | 0.0784 | 0.1643 | 0.03676 | 7.87E-06 |  |
|  | rs768029 | 4 | 73140229 | C | 0.01836 | 0.3273 | 0.07354 | 8.59E-06 | ADAMTS3 |
|  | rs10008978 | 4 | 157765129 | T | 0.07003 | 0.1797 | 0.03879 | 3.63E-06 | **PDGFC** |

*Chr = chromosome, BP = base pair, EA = effect allele, EAF = effect allele frequency, Beta = estimated effect, SE = estimated effect standard error, P = p-value. Genes encompassing loci are highlighted in bold.*

**Supplementary Table 20**. Lead SNPs of suggestive significance (1.00 x 10^-5^) from subgroup case-control GWAS (unadjusted for BMI)

|  | **rsID** | **Chr** | **BP** | **EA** | **EAF** | **Beta** | **SE** | **P** | **Nearest Gene** |
| --- | --- | --- | --- | --- | --- | --- | --- | --- | --- |
| ↑ Weight & Sleep | rs2164524 | 1 | 199281617 | T | 0.05 | 0.58 | 0.12 | 7.86E-07 |  |
|  | rs1426041 | 3 | 139697073 | A | 0.13 | 0.41 | 0.08 | 3.07E-07 | **CLSTN2** |
|  | rs9396811 | 6 | 17832478 | A | 0.04 | 0.63 | 0.13 | 2.47E-06 | **KIF13A** |
|  | rs3798756 | 6 | 152529260 | A | 0.13 | 0.36 | 0.08 | 3.62E-06 | SYNE1 |
|  | rs2157818 | 7 | 93528148 | T | 0.07 | 0.47 | 0.10 | 7.08E-06 | **GNGT1** |
|  | rs7099267 | 10 | 131212448 | G | 0.12 | 0.37 | 0.08 | 6.20E-06 | MGMT |
|  | rs12281926 | 11 | 29115687 | T | 0.02 | 0.93 | 0.19 | 1.26E-06 |  |
|  | rs17502383 | 13 | 30306277 | G | 0.17 | 0.31 | 0.07 | 8.93E-06 | UBL3 |
|  | rs16945081 | 17 | 59342375 | C | 0.14 | -0.35 | 0.08 | 8.48E-06 | **BCAS3** |
|  | rs2824977 | 21 | 20028118 | A | 0.01 | 1.27 | 0.26 | 1.40E-06 |  |
| ↓ Weight & Sleep | rs7519409 | 1 | 109399660 | G | 0.28 | 0.10 | 0.02 | 1.02E-06 | **AKNAD1** |
|  | rs7567744 | 2 | 55696124 | A | 0.17 | 0.12 | 0.03 | 1.09E-06 | CCDC88A |
|  | rs6786909 | 3 | 850922 | C | 0.34 | -0.10 | 0.02 | 2.22E-07 |  |
|  | rs4473877 | 6 | 144780581 | C | 0.02 | 0.30 | 0.07 | 3.10E-06 | **UTRN** |
|  | rs10271820 | 7 | 118777401 | A | 0.50 | 0.09 | 0.02 | 2.85E-06 |  |
| Uncategorised (Weight-Sleep) | rs12747042 | 1 | 245109861 | C | 0.13 | -0.09 | 0.02 | 3.17E-06 | EFCAB2 |
|  | rs2568871 | 3 | 196692717 | T | 0.39 | -0.07 | 0.01 | 2.75E-06 | **PIGZ** |
|  | rs8180202 | 4 | 93198845 | G | 0.09 | 0.11 | 0.02 | 5.80E-06 |  |
|  | rs10740076 | 10 | 64156110 | C | 0.32 | 0.07 | 0.01 | 7.25E-06 | **ZNF365** |
|  | rs10761940 | 10 | 67060807 | A | 0.37 | -0.07 | 0.01 | 3.41E-07 |  |
|  | rs2297620 | 14 | 66964538 | C | 0.13 | -0.10 | 0.02 | 2.86E-06 | GPHN |
|  | rs8008104 | 14 | 94186372 | G | 0.09 | 0.11 | 0.02 | 6.60E-06 | **PRIMA1** |
|  | rs11084813 | 19 | 35923858 | C | 0.26 | 0.07 | 0.02 | 5.36E-06 | FFAR2 |
| ↑ Weight | rs9792944 | 1 | 32419044 | T | 0.10 | 0.22 | 0.04 | 2.95E-07 | PTP4A2 |
|  | rs1867012 | 3 | 12188243 | C | 0.06 | 0.24 | 0.05 | 8.09E-06 | **SNY2** |
|  | rs1989916 | 5 | 136893719 | T | 0.37 | -0.12 | 0.03 | 8.06E-06 | **SPOCK1** |
|  | rs10216044 | 7 | 41812483 | T | 0.40 | 0.12 | 0.03 | 8.53E-06 |  |
|  | rs12442719 | 15 | 25966038 | C | 0.02 | 0.48 | 0.11 | 4.90E-06 | **ATP10A** |
|  | rs6132939 | 20 | 26239964 | A | 0.34 | -0.13 | 0.03 | 1.84E-06 | FAM182B |
| ↓ Weight | rs839859 | 1 | 109470185 | C | 0.28 | 0.09 | 0.02 | 9.99E-07 | **AKNAD1**, **GPSM2** |
|  | rs4233750 | 2 | 30102650 | T | 0.21 | 0.10 | 0.02 | 4.72E-06 | ALK |
|  | rs7567744 | 2 | 55696124 | A | 0.17 | 0.11 | 0.02 | 5.33E-06 |  |
|  | rs6786909 | 3 | 850922 | C | 0.34 | -0.09 | 0.02 | 6.62E-07 |  |
|  | rs2022929 | 8 | 110630465 | G | 0.09 | 0.14 | 0.03 | 5.29E-06 | SYBU |
|  | rs2219684 | 14 | 47136591 | A | 0.44 | -0.08 | 0.02 | 4.75E-06 | RPL10L |
|  | rs590076 | 18 | 53260732 | A | 0.34 | 0.08 | 0.02 | 7.70E-06 | **TCF4** |
|  | rs3744927 | 18 | 60245247 | A | 0.23 | -0.10 | 0.02 | 2.80E-06 | **ZCCHC2** |
| Uncategorised (Weight-only) | rs2079147 | 7 | 92332375 | A | 0.46 | -0.07 | 0.02 | 6.56E-06 | **CDK6** |
|  | rs4735916 | 8 | 677305 | A | 0.09 | -0.13 | 0.03 | 8.17E-06 | **ERICH1** |
|  | rs10995147 | 10 | 64165285 | A | 0.35 | 0.08 | 0.02 | 2.34E-06 | **ZNF365** |
|  | rs8022657 | 14 | 67390499 | T | 0.11 | -0.11 | 0.03 | 8.14E-06 | **GPHN** |
|  | rs3852787 | 16 | 72225733 | G | 0.50 | 0.08 | 0.02 | 7.08E-07 | PMFBP1 |
|  | rs11084813 | 19 | 35923858 | C | 0.26 | 0.09 | 0.02 | 8.15E-07 | FFAR2 |

*Chr = chromosome, BP = base pair, EA = effect allele, EAF = effect allele frequency, Beta = estimated effect, SE = estimated effect standard error, P = p-value. Genes encompassing loci are highlighted in bold.*

**Supplementary Table 21**. Lead SNPs of suggestive significance (1.00 x 10^-5^) from subgroup case-only GWAS (unadjusted for BMI)

|  | **rsID** | **Chr** | **BP** | **EA** | **EAF** | **Beta** | **SE** | **P** | **Nearest Gene** |
| --- | --- | --- | --- | --- | --- | --- | --- | --- | --- |
| ↑ Weight & Sleep  vs  ↓ Weight & Sleep | rs11691675 | 2 | 117257001 | G | 0.11 | 0.34 | 0.07 | 1.08E-06 |  |
|  | rs17062413 | 3 | 60152583 | A | 0.05 | 0.48 | 0.10 | 1.80E-06 |  |
|  | rs1426041 | 3 | 139697073 | A | 0.13 | 0.29 | 0.06 | 8.04E-06 | **CLSTN2** |
|  | rs7679231 | 4 | 136882276 | T | 0.11 | 0.36 | 0.07 | 1.98E-07 | **ANO1** |
|  | rs2509160 | 11 | 69970412 | G | 0.29 | 0.21 | 0.05 | 7.42E-06 | **LRRK2** |
|  | rs1491938 | 12 | 40645630 | G | 0.41 | -0.20 | 0.04 | 5.00E-06 | **PITPNM3** |
|  | rs897951 | 15 | 95567671 | A | 0.38 | 0.21 | 0.04 | 3.79E-06 |  |
|  | rs11657858 | 17 | 6366889 | T | 0.45 | -0.20 | 0.04 | 8.63E-06 |  |
|  | rs139553 | 22 | 42187199 | T | 0.21 | 0.24 | 0.05 | 6.76E-06 | **MEI1** |
| ↑ Weight & Sleep  vs  Uncategorised (Weight-Sleep) | rs11691675 | 2 | 117257001 | G | 0.11 | 0.32 | 0.07 | 4.61E-06 | DPP10 |
|  | rs28903985 | 2 | 234959167 | A | 0.02 | 0.78 | 0.17 | 2.83E-06 | SPP2 |
|  | rs1426041 | 3 | 139697073 | A | 0.13 | 0.29 | 0.06 | 7.19E-06 | **CLSTN2** |
|  | rs7782383 | 7 | 8737318 | G | 0.03 | 0.55 | 0.12 | 6.81E-06 | **NXPH1** |
|  | rs2410633 | 8 | 20060856 | T | 0.27 | -0.24 | 0.05 | 1.34E-06 | **ATP6V1B2** |
|  | rs12422304 | 12 | 93747731 | A | 0.43 | 0.20 | 0.04 | 9.55E-06 | NUDT4 |
|  | rs16945081 | 17 | 59342375 | C | 0.13 | -0.29 | 0.06 | 6.61E-06 | **BCAS3** |
| ↓ Weight & Sleep  vs  Uncategorised (Weight-Sleep) | rs884460 | 2 | 140926018 | C | 0.50 | 0.08 | 0.02 | 8.53E-06 |  |
|  | rs4647992 | 4 | 103455347 | T | 0.04 | -0.23 | 0.05 | 1.00E-06 | **NFKB1** |
|  | rs12006019 | 9 | 4215542 | T | 0.12 | -0.13 | 0.03 | 6.22E-06 | **GLIS3** |
|  | rs9901367 | 17 | 8982593 | T | 0.49 | 0.08 | 0.02 | 9.97E-06 | **NTN1** |
|  | rs911901 | 20 | 55952187 | A | 0.40 | 0.09 | 0.02 | 7.07E-06 | **RAE1** |
| ↑ Weight  vs  ↓ Weight | rs4688064 | 3 | 117551526 | T | 0.28 | 0.15 | 0.03 | 5.36E-08 | **LSAMP** |
|  | rs4130023 | 6 | 41934514 | T | 0.22 | 0.13 | 0.03 | 9.09E-06 | **CCND3** |
|  | rs12273363 | 11 | 27744859 | C | 0.21 | 0.14 | 0.03 | 5.74E-06 | BDNF |
|  | rs12149832 | 16 | 53842908 | A | 0.41 | 0.12 | 0.02 | 3.16E-06 | **FTO** |
|  | rs10775329 | 16 | 54751218 | G | 0.07 | 0.23 | 0.05 | 3.02E-06 |  |
| ↑ Weight  vs  Uncategorised (Weight-only) | rs10177805 | 2 | 235425433 | T | 0.07 | 0.22 | 0.05 | 5.77E-06 | ARL4C |
|  | rs7845912 | 8 | 72757932 | G | 0.13 | -0.17 | 0.04 | 2.29E-06 | MSC |
|  | rs1679029 | 9 | 22195321 | A | 0.39 | -0.12 | 0.03 | 4.56E-06 | CDKN2B |
|  | rs11027160 | 11 | 23363924 | T | 0.30 | -0.12 | 0.03 | 7.69E-06 | SLC17A6 |
|  | rs1595296 | 16 | 59022389 | A | 0.49 | -0.12 | 0.03 | 5.56E-06 |  |
|  | rs3852789 | 16 | 72228506 | C | 0.20 | 0.14 | 0.03 | 6.96E-06 | PMFBP1 |
| ↓ Weight  vs  Uncategorised (Weight-only) | rs1559497 | 2 | 31118058 | G | 0.47 | -0.09 | 0.02 | 7.92E-06 | CAPN13 |
|  | rs10008978 | 4 | 157765129 | T | 0.07 | 0.17 | 0.04 | 9.87E-06 | **PDGFC** |
|  | rs17071034 | 18 | 61048621 | T | 0.07 | 0.18 | 0.04 | 9.95E-06 | VPS4B |

*Chr = chromosome, BP = base pair, EA = effect allele, EAF = effect allele frequency, Beta = estimated effect, SE = estimated effect standard error, P = p-value. Genes encompassing loci are highlighted in bold.*

**Supplementary Table 22.** Results from comparisons of the estimated effect of rs4688064 in case-control and case-only GWAS

|  |  | GWAS 1 | Beta (SE) 1 | GWAS 2 | Beta (SE) 2 | P-value |
| --- | --- | --- | --- | --- | --- | --- |
| Adjusted for BMI | Case-control Analyses | ↑WS vs Controls | 0.09 (0.08) | ↓WS vs Controls | -0.06 (0.02) | 0.0642 |
|  |  | ↑WS vs Controls | 0.09 (0.08) | Uncategorised (WS) vs Controls | 0.03 (0.03) | 0.461 |
|  |  | ↓WS vs Controls | -0.06 (0.02) | Uncategorised (WS) vs Controls | 0.03 (0.03) | **6.61E-04** |
|  |  | ↑W vs Controls | 0.12 (0.03) | ↓W vs Controls | -0.05 (0.02) | **5.49E-06** |
|  |  | ↑W vs Controls | 0.12 (0.03) | Uncategorised (W) vs Controls | 0.02 (0.02) | **7.84E-03** |
|  |  | ↓W vs Controls | -0.05 (0.02) | Uncategorised (W) vs Controls | 0.02 (0.02) | **4.59E-03** |
|  | Case-only Analyses | ↑WS vs ↓WS | 0.19 (0.06) | ↑WS vs Uncategorised (WS) | 0.09 (0.05) | 0.201 |
|  |  | ↑WS vs ↓WS | 0.19 (0.06) | ↓WS vs Uncategorised (WS) | -0.08 (0.02) | **1.14E-05** |
|  |  | ↑WS vs Uncategorised (WS) | 0.09 (0.05) | ↓WS vs Uncategorised (WS) | -0.08 (0.02) | **9.20E-04** |
|  |  | ↑W vs ↓W | 0.16 (0.03) | ↑W vs Uncategorised (W) | 0.08 (0.03) | 0.0595 |
|  |  | ↑W vs ↓W | 0.16 (0.03) | ↓W vs Uncategorised (W) | -0.08 (0.02) | **2.96E-09** |
|  |  | ↑W vs Uncategorised (W) | 0.08 (0.03) | ↓W vs Uncategorised (W) | -0.08 (0.02) | **2.92E-05** |
| Unadjusted for BMI | Case-control Analyses | ↑WS vs Controls | 0.11 (0.06) | ↓WS vs Controls | -0.06 (0.02) | **8.20e-03** |
|  |  | ↑WS vs Controls | 0.11 (0.06) | Uncategorised (WS) vs Controls | 0.03 (0.02) | 0.226 |
|  |  | ↓WS vs Controls | -0.06 (0.02) | Uncategorised (WS) vs Controls | 0.03 (0.02) | **3.68e-04** |
|  |  | ↑W vs Controls | 0.12 (0.03) | ↓W vs Controls | -0.05 (0.03) | **5.05e-07** |
|  |  | ↑W vs Controls | 0.12 (0.03) | Uncategorised (W) vs Controls | 0.02 (0.02) | **3.73e-03** |
|  |  | ↓W vs Controls | -0.05 (0.03) | Uncategorised (W) vs Controls | 0.02 (0.02) | **3.28e-03** |
|  | Case-only Analyses | ↑WS vs ↓WS | 0.18 (0.05) | ↑WS vs Uncategorised (WS) | 0.10 (0.05) | 0.233 |
|  |  | ↑WS vs ↓WS | 0.18 (0.05) | ↓WS vs Uncategorised (WS) | -0.08 (0.02) | **4.93e-07** |
|  |  | ↑WS vs Uncategorised (WS) | 0.10 (0.05) | ↓WS vs Uncategorised (WS) | -0.08 (0.02) | **4.39e-04** |
|  |  | ↑W vs ↓W | 0.15 (0.03) | ↑W vs Uncategorised (W) | 0.08 (0.03) | 0.101 |
|  |  | ↑W vs ↓W | 0.15 (0.03) | ↓W vs Uncategorised (W) | -0.08 (0.02) | **1.39e-10** |
|  |  | ↑W vs Uncategorised (W) | 0.08 (0.03) | ↓W vs Uncategorised (W) | -0.08 (0.02) | **5.61e-06** |

*Beta = estimated effect, SE = estimated effect standard error*

**Supplementary Table 23.** Top 10 results of MAGMA gene-based tests of case-control GWAS (adjusted for BMI)

| **GWAS** | **Ensembl Gene ID** | **Chr** | **Start BP** | **Stop BP** | **Gene Symbol** | **Z-score** | **P-value** |
| --- | --- | --- | --- | --- | --- | --- | --- |
| ↑ Weight & Sleep | ENSG00000169220 | 5 | 176784838 | 176799602 | RGS14 | 4.95 | 3.76E-07 |
|  | ENSG00000127184 | 5 | 85913721 | 85916779 | COX7C | 4.26 | 1.05E-05 |
|  | ENSG00000196526 | 4 | 7760441 | 7941653 | AFAP1 | 3.96 | 3.68E-05 |
|  | ENSG00000169223 | 5 | 176758563 | 176778853 | LMAN2 | 3.75 | 8.82E-05 |
|  | ENSG00000139737 | 13 | 78272023 | 78338377 | SLAIN1 | 3.69 | 1.14E-04 |
|  | ENSG00000182944 | 22 | 29663998 | 29696515 | EWSR1 | 3.62 | 1.50E-04 |
|  | ENSG00000184302 | 14 | 60975669 | 60979568 | SIX6 | 3.58 | 1.75E-04 |
|  | ENSG00000123552 | 6 | 99880190 | 99969604 | USP45 | 3.57 | 1.78E-04 |
|  | ENSG00000179008 | 14 | 60863187 | 60982261 | C14orf39 | 3.54 | 2.00E-04 |
|  | ENSG00000100697 | 14 | 95552565 | 95624347 | DICER1 | 3.49 | 2.41E-04 |
| ↓ Weight & Sleep | ENSG00000162641 | 1 | 109358520 | 109506106 | AKNAD1 | 4.73 | 1.14E-06 |
|  | ENSG00000121957 | 1 | 109417972 | 109477167 | GPSM2 | 4.33 | 7.58E-06 |
|  | ENSG00000106460 | 7 | 12250867 | 12282993 | TMEM106B | 3.99 | 3.24E-05 |
|  | ENSG00000121940 | 1 | 109472130 | 109506111 | CLCC1 | 3.92 | 4.39E-05 |
|  | ENSG00000138594 | 15 | 52121825 | 52239492 | TMOD3 | 3.80 | 7.23E-05 |
|  | ENSG00000206536 | 3 | 98109510 | 98110475 | OR5K3 | 3.75 | 8.84E-05 |
|  | ENSG00000166477 | 15 | 52230222 | 52264003 | LEO1 | 3.74 | 9.37E-05 |
|  | ENSG00000112576 | 6 | 41902671 | 42018095 | CCND3 | 3.68 | 1.17E-04 |
|  | ENSG00000128872 | 15 | 52043758 | 52108565 | TMOD2 | 3.67 | 1.19E-04 |
|  | ENSG00000141664 | 18 | 60190240 | 60254942 | ZCCHC2 | 3.63 | 1.39E-04 |
| Uncategorised (Weight-Sleep) | ENSG00000138311 | 10 | 64133951 | 64431771 | ZNF365 | 4.73 | 1.15E-06 |
|  | ENSG00000171723 | 14 | 66974125 | 67648520 | GPHN | 4.02 | 2.86E-05 |
|  | ENSG00000102977 | 16 | 67691415 | 67694713 | ACD | 3.93 | 4.19E-05 |
|  | ENSG00000172717 | 14 | 67656110 | 67695267 | FAM71D | 3.86 | 5.58E-05 |
|  | ENSG00000119227 | 3 | 196673214 | 196695931 | PIGZ | 3.85 | 5.80E-05 |
|  | ENSG00000096654 | 6 | 27418522 | 27440897 | ZNF184 | 3.78 | 7.78E-05 |
|  | ENSG00000158457 | 7 | 128784712 | 128808671 | TSPAN33 | 3.67 | 1.19E-04 |
|  | ENSG00000100568 | 14 | 68113792 | 68141548 | VTI1B | 3.63 | 1.43E-04 |
|  | ENSG00000196628 | 18 | 52889562 | 53332018 | TCF4 | 3.62 | 1.45E-04 |
|  | ENSG00000197935 | 6 | 28962562 | 28973093 | ZNF311 | 3.62 | 1.46E-04 |
| ↑ Weight | ENSG00000078328 | 16 | 6069095 | 7763340 | RBFOX1 | 3.57 | 1.78E-04 |
|  | ENSG00000005007 | 19 | 18942747 | 18979045 | UPF1 | 3.55 | 1.91E-04 |
|  | ENSG00000112078 | 6 | 36410544 | 36458920 | KCTD20 | 3.53 | 2.05E-04 |
|  | ENSG00000075292 | 2 | 71503691 | 71662199 | ZNF638 | 3.45 | 2.82E-04 |
|  | ENSG00000129451 | 19 | 51515995 | 51523431 | KLK10 | 3.43 | 2.99E-04 |
|  | ENSG00000112079 | 6 | 36461669 | 36515247 | STK38 | 3.42 | 3.08E-04 |
|  | ENSG00000127928 | 7 | 93220885 | 93540577 | GNGT1 | 3.37 | 3.77E-04 |
|  | ENSG00000127184 | 5 | 85913721 | 85916779 | COX7C | 3.34 | 4.12E-04 |
|  | ENSG00000180245 | 4 | 110749150 | 110765760 | RRH | 3.34 | 4.22E-04 |
|  | ENSG00000174928 | 3 | 155480401 | 155524140 | C3orf33 | 3.30 | 4.79E-04 |
| ↓ Weight | ENSG00000162641 | 1 | 109358520 | 109506106 | AKNAD1 | 4.58 | 2.35E-06 |
|  | ENSG00000121957 | 1 | 109417972 | 109477167 | GPSM2 | 4.53 | 2.92E-06 |
|  | ENSG00000141664 | 18 | 60190240 | 60254942 | ZCCHC2 | 4.10 | 2.11E-05 |
|  | ENSG00000121940 | 1 | 109472130 | 109506111 | CLCC1 | 4.06 | 2.43E-05 |
|  | ENSG00000196628 | 18 | 52889562 | 53332018 | TCF4 | 4.03 | 2.83E-05 |
|  | ENSG00000248050 | 8 | 110656344 | 110660313 | RP11-422N16.3 | 4.01 | 3.06E-05 |
|  | ENSG00000267893 | 17 | 70036164 | 70036872 | AC007461.1 | 3.98 | 3.38E-05 |
|  | ENSG00000196277 | 3 | 6811688 | 7783215 | GRM7 | 3.92 | 4.50E-05 |
|  | ENSG00000187954 | 8 | 145674965 | 145691060 | CYHR1 | 3.80 | 7.35E-05 |
|  | ENSG00000106460 | 7 | 12250867 | 12282993 | TMEM106B | 3.59 | 1.62E-04 |
| Uncategorised (Weight-only) | ENSG00000138311 | 10 | 64133951 | 64431771 | ZNF365 | 4.71 | 1.22E-06 |
|  | ENSG00000145934 | 5 | 166711804 | 167691162 | TENM2 | 4.27 | 9.92E-06 |
|  | ENSG00000171723 | 14 | 66974125 | 67648520 | GPHN | 3.92 | 4.50E-05 |
|  | ENSG00000102977 | 16 | 67691415 | 67694713 | ACD | 3.84 | 6.25E-05 |
|  | ENSG00000172717 | 14 | 67656110 | 67695267 | FAM71D | 3.83 | 6.37E-05 |
|  | ENSG00000183653 | 21 | 30968360 | 31003067 | GRIK1-AS2 | 3.78 | 7.95E-05 |
|  | ENSG00000158457 | 7 | 128784712 | 128808671 | TSPAN33 | 3.56 | 1.87E-04 |
|  | ENSG00000159685 | 3 | 126423063 | 126679249 | CHCHD6 | 3.47 | 2.56E-04 |
|  | ENSG00000103187 | 16 | 84599200 | 84651683 | COTL1 | 3.47 | 2.63E-04 |
|  | ENSG00000118557 | 16 | 72146056 | 72210777 | PMFBP1 | 3.42 | 3.15E-04 |

*Chr = chromosome, BP = base pair*

**Supplementary Table 24.** Top 10 results of MAGMA gene-based tests of case-only GWAS (adjusted for BMI)

| **GWAS** | **Ensembl Gene ID** | **Chr** | **Start BP** | **Stop BP** | **Gene Symbol** | **Z-score** | **P-value** |
| --- | --- | --- | --- | --- | --- | --- | --- |
| ↑ Weight & Sleep  vs  ↓ Weight & Sleep | ENSG00000167074 | 22 | 41763337 | 41795330 | TEF | 3.92 | 4.45E-05 |
|  | ENSG00000172346 | 22 | 41956767 | 41973745 | CSDC2 | 3.72 | 1.02E-04 |
|  | ENSG00000183098 | 13 | 93879095 | 95059655 | GPC6 | 3.71 | 1.05E-04 |
|  | ENSG00000167077 | 22 | 42095503 | 42195460 | MEI1 | 3.66 | 1.25E-04 |
|  | ENSG00000100410 | 22 | 41855721 | 41864729 | PHF5A | 3.66 | 1.27E-04 |
|  | ENSG00000170113 | 15 | 23043277 | 23100005 | NIPA1 | 3.64 | 1.37E-04 |
|  | ENSG00000100418 | 22 | 41994032 | 42017100 | DESI1 | 3.61 | 1.52E-04 |
|  | ENSG00000100412 | 22 | 41865129 | 41924993 | ACO2 | 3.54 | 1.99E-04 |
|  | ENSG00000253379 | 8 | 72315675 | 72504260 | RP11-1102P16.1 | 3.53 | 2.06E-04 |
|  | ENSG00000100413 | 22 | 41921808 | 41940610 | POLR3H | 3.44 | 2.94E-04 |
| ↑ Weight & Sleep  vs  Uncategorised (Weight-Sleep) | ENSG00000183098 | 13 | 93879095 | 95059655 | GPC6 | 3.90 | 4.89E-05 |
|  | ENSG00000154832 | 18 | 47808713 | 47814674 | CXXC1 | 3.86 | 5.57E-05 |
|  | ENSG00000147416 | 8 | 20054878 | 20084330 | ATP6V1B2 | 3.77 | 8.11E-05 |
|  | ENSG00000130713 | 9 | 133569108 | 133580248 | EXOSC2 | 3.54 | 2.04E-04 |
|  | ENSG00000167077 | 22 | 42095503 | 42195460 | MEI1 | 3.53 | 2.11E-04 |
|  | ENSG00000167074 | 22 | 41763337 | 41795330 | TEF | 3.51 | 2.23E-04 |
|  | ENSG00000145020 | 3 | 49454211 | 49460186 | AMT | 3.48 | 2.53E-04 |
|  | ENSG00000173402 | 3 | 49506146 | 49573048 | DAG1 | 3.46 | 2.74E-04 |
|  | ENSG00000169220 | 5 | 176784838 | 176799602 | RGS14 | 3.41 | 3.22E-04 |
|  | ENSG00000141644 | 18 | 47793252 | 47808144 | MBD1 | 3.40 | 3.37E-04 |
| ↓ Weight & Sleep  vs  Uncategorised (Weight-Sleep) | ENSG00000101146 | 20 | 55926066 | 55954267 | RAE1 | 4.39 | 5.75E-06 |
|  | ENSG00000132376 | 17 | 1397865 | 1420182 | INPP5K | 3.79 | 7.50E-05 |
|  | ENSG00000197566 | 17 | 16524051 | 16557170 | ZNF624 | 3.59 | 1.67E-04 |
|  | ENSG00000132819 | 20 | 55966463 | 55984389 | RBM38 | 3.41 | 3.30E-04 |
|  | ENSG00000145354 | 4 | 103790135 | 103810399 | CISD2 | 3.37 | 3.70E-04 |
|  | ENSG00000184602 | 16 | 11762270 | 11773015 | SNN | 3.37 | 3.75E-04 |
|  | ENSG00000107249 | 9 | 3824127 | 4348392 | GLIS3 | 3.25 | 5.72E-04 |
|  | ENSG00000162641 | 1 | 109358520 | 109506106 | AKNAD1 | 3.25 | 5.86E-04 |
|  | ENSG00000182545 | 14 | 20973696 | 20979328 | RNASE10 | 3.24 | 5.98E-04 |
|  | ENSG00000109332 | 4 | 103715540 | 103790053 | UBE2D3 | 3.22 | 6.43E-04 |
| ↑ Weight  vs  ↓ Weight | ENSG00000183876 | 5 | 149675906 | 149718870 | ARSI | 4.35 | 6.84E-06 |
|  | ENSG00000105193 | 19 | 39923847 | 39926588 | RPS16 | 3.75 | 8.80E-05 |
|  | ENSG00000213922 | 19 | 39930212 | 39932082 | AC011500.1 | 3.70 | 1.09E-04 |
|  | ENSG00000198060 | 10 | 94050920 | 94113721 | MARCH5 | 3.53 | 2.05E-04 |
|  | ENSG00000187905 | 22 | 21400249 | 21418457 | AC002472.13 | 3.43 | 3.04E-04 |
|  | ENSG00000185565 | 3 | 115521235 | 117716095 | LSAMP | 3.40 | 3.43E-04 |
|  | ENSG00000186838 | 19 | 40005753 | 40011326 | SELV | 3.39 | 3.44E-04 |
|  | ENSG00000172331 | 7 | 134331560 | 134364565 | BPGM | 3.38 | 3.67E-04 |
|  | ENSG00000148288 | 9 | 136028340 | 136039332 | GBGT1 | 3.32 | 4.57E-04 |
|  | ENSG00000140987 | 16 | 3432085 | 3451065 | ZSCAN32 | 3.31 | 4.73E-04 |
| ↑ Weight  vs  Uncategorised (Weight-only) | ENSG00000187772 | 6 | 105404923 | 105531207 | LIN28B | 4.08 | 2.21E-05 |
|  | ENSG00000156050 | 14 | 74398204 | 74417117 | FAM161B | 3.95 | 3.94E-05 |
|  | ENSG00000118557 | 16 | 72146056 | 72210777 | PMFBP1 | 3.70 | 1.09E-04 |
|  | ENSG00000119725 | 14 | 74353320 | 74399214 | ZNF410 | 3.69 | 1.12E-04 |
|  | ENSG00000127184 | 5 | 85913721 | 85916779 | COX7C | 3.69 | 1.12E-04 |
|  | ENSG00000005007 | 19 | 18942747 | 18979045 | UPF1 | 3.48 | 2.50E-04 |
|  | ENSG00000127928 | 7 | 93220885 | 93540577 | GNGT1 | 3.45 | 2.85E-04 |
|  | ENSG00000112539 | 6 | 165693153 | 165723096 | C6orf118 | 3.42 | 3.12E-04 |
|  | ENSG00000079246 | 2 | 216972187 | 217071026 | XRCC5 | 3.38 | 3.67E-04 |
|  | ENSG00000140829 | 16 | 72127461 | 72146811 | DHX38 | 3.37 | 3.72E-04 |
| ↓ Weight  vs  Uncategorised (Weight-only) | ENSG00000139874 | 14 | 38677204 | 38682272 | SSTR1 | 3.75 | 8.77E-05 |
|  | ENSG00000172955 | 4 | 100123795 | 100140694 | ADH6 | 3.47 | 2.57E-04 |
|  | ENSG00000162641 | 1 | 109358520 | 109506106 | AKNAD1 | 3.43 | 3.05E-04 |
|  | ENSG00000057757 | 1 | 24104895 | 24114722 | PITHD1 | 3.40 | 3.32E-04 |
|  | ENSG00000184602 | 16 | 11762270 | 11773015 | SNN | 3.37 | 3.72E-04 |
|  | ENSG00000132376 | 17 | 1397865 | 1420182 | INPP5K | 3.35 | 4.02E-04 |
|  | ENSG00000198680 | 9 | 25676396 | 25678856 | TUSC1 | 3.33 | 4.37E-04 |
|  | ENSG00000182132 | 5 | 169780491 | 170163636 | KCNIP1 | 3.32 | 4.56E-04 |
|  | ENSG00000121957 | 1 | 109417972 | 109477167 | GPSM2 | 3.28 | 5.26E-04 |
|  | ENSG00000131791 | 1 | 146626685 | 146644129 | PRKAB2 | 3.24 | 5.91E-04 |

*Chr = chromosome, BP = base pair*

**Supplementary Table 25.** Top 10 results of MAGMA gene-based tests of case-control GWAS (unadjusted for BMI)

| **GWAS** | **Ensembl Gene ID** | **Chr** | **Start BP** | **Stop BP** | **Gene Symbol** | **Z-score** | **P-value** |
| --- | --- | --- | --- | --- | --- | --- | --- |
| ↑ Weight & Sleep | ENSG00000196526 | 4 | 7760441 | 7941653 | AFAP1 | 4.13 | 1.80E-05 |
|  | ENSG00000154832 | 18 | 47808713 | 47814674 | CXXC1 | 3.85 | 6.02E-05 |
|  | ENSG00000204681 | 6 | 29523406 | 29601753 | GABBR1 | 3.71 | 1.04E-04 |
|  | ENSG00000188906 | 12 | 40590546 | 40763087 | LRRK2 | 3.69 | 1.12E-04 |
|  | ENSG00000167074 | 22 | 41763337 | 41795330 | TEF | 3.59 | 1.64E-04 |
|  | ENSG00000137161 | 6 | 42896938 | 42907025 | CNPY3 | 3.56 | 1.88E-04 |
|  | ENSG00000100697 | 14 | 95552565 | 95624347 | DICER1 | 3.52 | 2.14E-04 |
|  | ENSG00000143149 | 1 | 165631453 | 165668100 | ALDH9A1 | 3.52 | 2.20E-04 |
|  | ENSG00000111911 | 6 | 126277927 | 126301387 | HINT3 | 3.42 | 3.08E-04 |
|  | ENSG00000183784 | 9 | 213108 | 215893 | C9orf66 | 3.42 | 3.17E-04 |
| ↓ Weight & Sleep | ENSG00000162641 | 1 | 109358520 | 109506106 | AKNAD1 | 4.72 | 1.16E-06 |
|  | ENSG00000121957 | 1 | 109417972 | 109477167 | GPSM2 | 4.31 | 8.24E-06 |
|  | ENSG00000106460 | 7 | 12250867 | 12282993 | TMEM106B | 4.06 | 2.45E-05 |
|  | ENSG00000121940 | 1 | 109472130 | 109506111 | CLCC1 | 3.94 | 4.12E-05 |
|  | ENSG00000138594 | 15 | 52121825 | 52239492 | TMOD3 | 3.84 | 6.28E-05 |
|  | ENSG00000166477 | 15 | 52230222 | 52264003 | LEO1 | 3.78 | 7.73E-05 |
|  | ENSG00000112576 | 6 | 41902671 | 42018095 | CCND3 | 3.72 | 1.01E-04 |
|  | ENSG00000128872 | 15 | 52043758 | 52108565 | TMOD2 | 3.70 | 1.07E-04 |
|  | ENSG00000206536 | 3 | 98109510 | 98110475 | OR5K3 | 3.66 | 1.28E-04 |
|  | ENSG00000141664 | 18 | 60190240 | 60254942 | ZCCHC2 | 3.61 | 1.52E-04 |
| Uncategorised (Weight-Sleep) | ENSG00000138311 | 10 | 64133951 | 64431771 | ZNF365 | 4.70 | 1.33E-06 |
|  | ENSG00000171723 | 14 | 66974125 | 67648520 | GPHN | 4.14 | 1.74E-05 |
|  | ENSG00000172717 | 14 | 67656110 | 67695267 | FAM71D | 4.00 | 3.23E-05 |
|  | ENSG00000119227 | 3 | 196673214 | 196695931 | PIGZ | 3.99 | 3.28E-05 |
|  | ENSG00000196628 | 18 | 52889562 | 53332018 | TCF4 | 3.91 | 4.62E-05 |
|  | ENSG00000102977 | 16 | 67691415 | 67694713 | ACD | 3.75 | 8.95E-05 |
|  | ENSG00000081181 | 14 | 68086515 | 68118437 | ARG2 | 3.68 | 1.19E-04 |
|  | ENSG00000100811 | 14 | 100704635 | 100749129 | YY1 | 3.66 | 1.25E-04 |
|  | ENSG00000100568 | 14 | 68113792 | 68141548 | VTI1B | 3.61 | 1.51E-04 |
|  | ENSG00000096654 | 6 | 27418522 | 27440897 | ZNF184 | 3.59 | 1.66E-04 |
| ↑ Weight | ENSG00000129744 | 11 | 3666358 | 3685646 | ART1 | 3.65 | 1.31E-04 |
|  | ENSG00000161973 | 17 | 8633252 | 8648537 | CCDC42 | 3.58 | 1.71E-04 |
|  | ENSG00000178096 | 1 | 149859440 | 149872351 | BOLA1 | 3.58 | 1.75E-04 |
|  | ENSG00000186130 | 9 | 125670335 | 125675609 | ZBTB6 | 3.52 | 2.13E-04 |
|  | ENSG00000188931 | 1 | 161334521 | 161337664 | C1orf192 | 3.47 | 2.65E-04 |
|  | ENSG00000142185 | 21 | 45770046 | 45862964 | TRPM2 | 3.46 | 2.73E-04 |
|  | ENSG00000225190 | 17 | 43513266 | 43568115 | PLEKHM1 | 3.44 | 2.88E-04 |
|  | ENSG00000128891 | 15 | 40820882 | 40857256 | C15orf57 | 3.38 | 3.60E-04 |
|  | ENSG00000005007 | 19 | 18942747 | 18979045 | UPF1 | 3.33 | 4.33E-04 |
|  | ENSG00000152520 | 13 | 28712643 | 28869475 | PAN3 | 3.31 | 4.66E-04 |
| ↓ Weight | ENSG00000162641 | 1 | 109358520 | 109506106 | AKNAD1 | 4.65 | 1.65E-06 |
|  | ENSG00000121957 | 1 | 109417972 | 109477167 | GPSM2 | 4.57 | 2.43E-06 |
|  | ENSG00000196628 | 18 | 52889562 | 53332018 | TCF4 | 4.09 | 2.15E-05 |
|  | ENSG00000121940 | 1 | 109472130 | 109506111 | CLCC1 | 4.08 | 2.22E-05 |
|  | ENSG00000141664 | 18 | 60190240 | 60254942 | ZCCHC2 | 4.07 | 2.39E-05 |
|  | ENSG00000248050 | 8 | 110656344 | 110660313 | RP11-422N16.3 | 4.03 | 2.75E-05 |
|  | ENSG00000267893 | 17 | 70036164 | 70036872 | AC007461.1 | 3.94 | 4.07E-05 |
|  | ENSG00000196277 | 3 | 6811688 | 7783215 | GRM7 | 3.85 | 5.98E-05 |
|  | ENSG00000187954 | 8 | 145674965 | 145691060 | CYHR1 | 3.77 | 8.17E-05 |
|  | ENSG00000106460 | 7 | 12250867 | 12282993 | TMEM106B | 3.66 | 1.24E-04 |
| Uncategorised (Weight-only) | ENSG00000138311 | 10 | 64133951 | 64431771 | ZNF365 | 4.72 | 1.18E-06 |
|  | ENSG00000145934 | 5 | 166711804 | 167691162 | TENM2 | 4.11 | 1.99E-05 |
|  | ENSG00000171723 | 14 | 66974125 | 67648520 | GPHN | 3.95 | 3.86E-05 |
|  | ENSG00000172717 | 14 | 67656110 | 67695267 | FAM71D | 3.84 | 6.20E-05 |
|  | ENSG00000102977 | 16 | 67691415 | 67694713 | ACD | 3.74 | 9.07E-05 |
|  | ENSG00000183653 | 21 | 30968360 | 31003067 | GRIK1-AS2 | 3.67 | 1.21E-04 |
|  | ENSG00000118557 | 16 | 72146056 | 72210777 | PMFBP1 | 3.66 | 1.26E-04 |
|  | ENSG00000158457 | 7 | 128784712 | 128808671 | TSPAN33 | 3.53 | 2.06E-04 |
|  | ENSG00000103187 | 16 | 84599200 | 84651683 | COTL1 | 3.45 | 2.79E-04 |
|  | ENSG00000159685 | 3 | 126423063 | 126679249 | CHCHD6 | 3.41 | 3.23E-04 |

*Chr = chromosome, BP = base pair*

**Supplementary Table 26.** Top 10 results of MAGMA gene-based tests of case-only GWAS (unadjusted for BMI)

| **GWAS** | **Ensembl Gene ID** | **Chr** | **Start BP** | **Stop BP** | **Gene Symbol** | **Z-score** | **P-value** |
| --- | --- | --- | --- | --- | --- | --- | --- |
| ↑ Weight & Sleep  vs  ↓ Weight & Sleep | ENSG00000167074 | 22 | 41763337 | 41795330 | TEF | 4.32 | 7.82E-06 |
|  | ENSG00000167077 | 22 | 42095503 | 42195460 | MEI1 | 4.28 | 9.40E-06 |
|  | ENSG00000188906 | 12 | 40590546 | 40763087 | LRRK2 | 4.21 | 1.25E-05 |
|  | ENSG00000100418 | 22 | 41994032 | 42017100 | DESI1 | 3.92 | 4.49E-05 |
|  | ENSG00000198060 | 10 | 94050920 | 94113721 | MARCH5 | 3.88 | 5.19E-05 |
|  | ENSG00000100138 | 22 | 42069934 | 42086508 | NHP2L1 | 3.80 | 7.16E-05 |
|  | ENSG00000172346 | 22 | 41956767 | 41973745 | CSDC2 | 3.79 | 7.58E-05 |
|  | ENSG00000100410 | 22 | 41855721 | 41864729 | PHF5A | 3.73 | 9.76E-05 |
|  | ENSG00000134504 | 18 | 24034874 | 24237365 | KCTD1 | 3.69 | 1.10E-04 |
|  | ENSG00000184860 | 16 | 82031221 | 82045093 | SDR42E1 | 3.69 | 1.13E-04 |
| ↑ Weight & Sleep  vs  Uncategorised (Weight-Sleep) | ENSG00000154832 | 18 | 47808713 | 47814674 | CXXC1 | 3.98 | 3.47E-05 |
|  | ENSG00000174989 | 12 | 117348761 | 117468953 | FBXW8 | 3.76 | 8.34E-05 |
|  | ENSG00000147416 | 8 | 20054878 | 20084330 | ATP6V1B2 | 3.65 | 1.30E-04 |
|  | ENSG00000188906 | 12 | 40590546 | 40763087 | LRRK2 | 3.65 | 1.33E-04 |
|  | ENSG00000111664 | 12 | 6949118 | 6956557 | GNB3 | 3.48 | 2.54E-04 |
|  | ENSG00000111665 | 12 | 6953957 | 6961230 | CDCA3 | 3.48 | 2.54E-04 |
|  | ENSG00000167074 | 22 | 41763337 | 41795330 | TEF | 3.46 | 2.70E-04 |
|  | ENSG00000167077 | 22 | 42095503 | 42195460 | MEI1 | 3.45 | 2.85E-04 |
|  | ENSG00000172361 | 18 | 47753563 | 47792892 | CCDC11 | 3.40 | 3.36E-04 |
|  | ENSG00000090061 | 14 | 99947506 | 100001381 | CCNK | 3.37 | 3.75E-04 |
| ↓ Weight & Sleep  vs  Uncategorised (Weight-Sleep) | ENSG00000101146 | 20 | 55926066 | 55954267 | RAE1 | 4.52 | 3.10E-06 |
|  | ENSG00000182545 | 14 | 20973696 | 20979328 | RNASE10 | 3.79 | 7.48E-05 |
|  | ENSG00000197566 | 17 | 16524051 | 16557170 | ZNF624 | 3.75 | 8.95E-05 |
|  | ENSG00000162641 | 1 | 109358520 | 109506106 | AKNAD1 | 3.60 | 1.60E-04 |
|  | ENSG00000065320 | 17 | 8924859 | 9147317 | NTN1 | 3.50 | 2.36E-04 |
|  | ENSG00000105662 | 19 | 18794487 | 18893004 | CRTC1 | 3.44 | 2.93E-04 |
|  | ENSG00000184602 | 16 | 11762270 | 11773015 | SNN | 3.35 | 4.03E-04 |
|  | ENSG00000172955 | 4 | 100123795 | 100140694 | ADH6 | 3.34 | 4.14E-04 |
|  | ENSG00000132376 | 17 | 1397865 | 1420182 | INPP5K | 3.32 | 4.50E-04 |
|  | ENSG00000166262 | 15 | 49619159 | 49913128 | FAM227B | 3.26 | 5.64E-04 |
| ↑ Weight  vs  ↓ Weight | ENSG00000176697 | 11 | 27676440 | 27743605 | BDNF | 4.40 | 5.41E-06 |
|  | ENSG00000198060 | 10 | 94050920 | 94113721 | MARCH5 | 4.23 | 1.15E-05 |
|  | ENSG00000140718 | 16 | 53737875 | 54155853 | FTO | 4.06 | 2.48E-05 |
|  | ENSG00000183876 | 5 | 149675906 | 149718870 | ARSI | 4.03 | 2.83E-05 |
|  | ENSG00000105193 | 19 | 39923847 | 39926588 | RPS16 | 4.01 | 3.00E-05 |
|  | ENSG00000213922 | 19 | 39930212 | 39932082 | AC011500.1 | 3.90 | 4.74E-05 |
|  | ENSG00000184602 | 16 | 11762270 | 11773015 | SNN | 3.90 | 4.76E-05 |
|  | ENSG00000204427 | 6 | 31654726 | 31671221 | ABHD16A | 3.82 | 6.79E-05 |
|  | ENSG00000198792 | 22 | 38615298 | 38669040 | TMEM184B | 3.80 | 7.30E-05 |
|  | ENSG00000196296 | 16 | 28889726 | 28915830 | ATP2A1 | 3.75 | 8.91E-05 |
| ↑ Weight  vs  Uncategorised (Weight-only) | ENSG00000118557 | 16 | 72146056 | 72210777 | PMFBP1 | 4.03 | 2.77E-05 |
|  | ENSG00000170647 | 11 | 100862811 | 100864663 | TMEM133 | 3.86 | 5.66E-05 |
|  | ENSG00000156050 | 14 | 74398204 | 74417117 | FAM161B | 3.60 | 1.59E-04 |
|  | ENSG00000187772 | 6 | 105404923 | 105531207 | LIN28B | 3.57 | 1.76E-04 |
|  | ENSG00000138767 | 4 | 78634541 | 78740769 | CNOT6L | 3.54 | 2.00E-04 |
|  | ENSG00000118785 | 4 | 88896819 | 88904562 | SPP1 | 3.53 | 2.08E-04 |
|  | ENSG00000140829 | 16 | 72127461 | 72146811 | DHX38 | 3.49 | 2.38E-04 |
|  | ENSG00000198060 | 10 | 94050920 | 94113721 | MARCH5 | 3.35 | 4.04E-04 |
|  | ENSG00000127184 | 5 | 85913721 | 85916779 | COX7C | 3.33 | 4.38E-04 |
|  | ENSG00000204519 | 19 | 58193337 | 58228669 | ZNF551 | 3.32 | 4.53E-04 |
| ↓ Weight  vs  Uncategorised (Weight-only) | ENSG00000139874 | 14 | 38677204 | 38682272 | SSTR1 | 3.71 | 1.05E-04 |
|  | ENSG00000172955 | 4 | 100123795 | 100140694 | ADH6 | 3.67 | 1.22E-04 |
|  | ENSG00000162641 | 1 | 109358520 | 109506106 | AKNAD1 | 3.55 | 1.92E-04 |
|  | ENSG00000131791 | 1 | 146626685 | 146644129 | PRKAB2 | 3.46 | 2.71E-04 |
|  | ENSG00000100711 | 14 | 104182067 | 104200005 | ZFYVE21 | 3.42 | 3.14E-04 |
|  | ENSG00000057757 | 1 | 24104895 | 24114722 | PITHD1 | 3.42 | 3.17E-04 |
|  | ENSG00000184602 | 16 | 11762270 | 11773015 | SNN | 3.39 | 3.48E-04 |
|  | ENSG00000065320 | 17 | 8924859 | 9147317 | NTN1 | 3.39 | 3.51E-04 |
|  | ENSG00000121957 | 1 | 109417972 | 109477167 | GPSM2 | 3.37 | 3.72E-04 |
|  | ENSG00000182132 | 5 | 169780491 | 170163636 | KCNIP1 | 3.29 | 5.00E-04 |

*Chr = chromosome, BP = base pair*

**Supplementary Table 27**. Polygenic risk score results of subgroup PRS from UK Biobank case-control GWAS (adjusted for BMI) tested for association in Generation Scotland samples

| UK Biobank PRS | | | Generation Scotland Sample | R-squared (Observed) | P-value | Coefficient | Standard Error |
| --- | --- | --- | --- | --- | --- | --- | --- |
| GWAS Summary Statistics | P-value Threshold | Number of SNPs |  |  |  |  |  |
| ↑WS & Controls | 5 x 10^-8^ | 3 | ↑WS & Controls | 4.80E-04 | 2.18E-01 | 0.156 | 0.126 |
|  |  |  | ↓WS & Controls | 2.94E-04 | 3.25E-01 | -0.058 | 0.059 |
|  |  |  | Uncategorised (WS) & Controls | 2.41E-06 | 9.29E-01 | -0.004 | 0.048 |
|  |  |  | ↑WS & ↓WS | 2.03E-03 | 3.46E-01 | 0.129 | 0.137 |
|  |  |  | ↑WS & Uncategorised (WS) | 1.70E-03 | 3.45E-01 | 0.129 | 0.137 |
|  |  |  | ↓WS & Uncategorised (WS) | 5.84E-05 | 8.29E-01 | -0.016 | 0.072 |
|  |  |  | ↑W & Controls | 2.26E-05 | 7.87E-01 | -0.019 | 0.069 |
|  |  |  | ↓W & Controls | 3.50E-04 | 2.81E-01 | -0.059 | 0.055 |
|  |  |  | Uncategorised (W) & Controls | 3.58E-04 | 2.81E-01 | 0.067 | 0.062 |
|  |  |  | ↑W & ↓W | 1.29E-05 | 9.28E-01 | 0.007 | 0.082 |
|  |  |  | ↑W & Uncategorised (W) | 2.94E-03 | 2.45E-01 | -0.107 | 0.092 |
|  |  |  | ↓W & Uncategorised (W) | 2.04E-03 | 2.46E-01 | -0.096 | 0.082 |
| ↓WS & Controls | 0.1 | 20294 | ↑WS & Controls | 1.89E-05 | 8.07E-01 | 0.031 | 0.128 |
|  |  |  | **↓WS & Controls** | **2.73E-03** | **2.66E-03** | **0.179** | **0.059** |
|  |  |  | Uncategorised (WS) & Controls | 5.30E-04 | 1.84E-01 | 0.065 | 0.049 |
|  |  |  | ↑WS & ↓WS | 1.10E-03 | 4.87E-01 | -0.097 | 0.140 |
|  |  |  | ↑WS & Uncategorised (WS) | 3.84E-05 | 8.87E-01 | -0.021 | 0.149 |
|  |  |  | ↓WS & Uncategorised (WS) | 3.59E-03 | 8.91E-02 | 0.125 | 0.073 |
|  |  |  | ↑W & Controls | 6.06E-04 | 1.62E-01 | 0.098 | 0.070 |
|  |  |  | **↓W & Controls** | **2.52E-03** | **3.83E-03** | **0.159** | **0.055** |
|  |  |  | Uncategorised (W) & Controls | 1.07E-06 | 9.53E-01 | 0.004 | 0.063 |
|  |  |  | ↑W & ↓W | 1.84E-03 | 2.79E-01 | -0.091 | 0.084 |
|  |  |  | ↑W & Uncategorised (W) | 2.33E-03 | 3.00E-01 | 0.105 | 0.101 |
|  |  |  | ↓W & Uncategorised (W) | 6.41E-03 | 3.93E-02 | 0.170 | 0.082 |
| Uncategorised (WS) & Controls | 0.1 | 20336 | ↑WS & Controls | 1.54E-06 | 9.44E-01 | -0.009 | 0.128 |
|  |  |  | ↓WS & Controls | 2.64E-04 | 3.50E-01 | 0.056 | 0.060 |
|  |  |  | **Uncategorised (WS) & Controls** | **2.82E-03** | **2.16E-03** | **0.149** | **0.048** |
|  |  |  | ↑WS & ↓WS | 3.35E-03 | 2.25E-01 | -0.168 | 0.138 |
|  |  |  | ↑WS & Uncategorised (WS) | 3.83E-03 | 1.56E-01 | -0.201 | 0.141 |
|  |  |  | ↓WS & Uncategorised (WS) | 3.88E-03 | 7.72E-02 | -0.128 | 0.072 |
|  |  |  | ↑W & Controls | 2.46E-04 | 3.72E-01 | 0.063 | 0.070 |
|  |  |  | ↓W & Controls | 5.35E-04 | 1.83E-01 | 0.074 | 0.055 |
|  |  |  | Uncategorised (W) & Controls | 1.96E-03 | 1.16E-02 | 0.158 | 0.062 |
|  |  |  | ↑W & ↓W | 6.56E-05 | 8.38E-01 | -0.017 | 0.083 |
|  |  |  | ↑W & Uncategorised (W) | 4.97E-03 | 1.30E-01 | -0.152 | 0.100 |
|  |  |  | ↓W & Uncategorised (W) | 1.42E-03 | 3.33E-01 | -0.077 | 0.080 |
| ↑W & Controls | 5 x 10^-4^ | 187 | ↑WS & Controls | 1.32E-03 | 4.09E-02 | -0.265 | 0.130 |
|  |  |  | ↓WS & Controls | 2.98E-04 | 3.21E-01 | -0.060 | 0.060 |
|  |  |  | Uncategorised (WS) & Controls | 2.65E-04 | 3.47E-01 | -0.046 | 0.049 |
|  |  |  | ↑WS & ↓WS | 6.26E-03 | 9.70E-02 | -0.226 | 0.136 |
|  |  |  | ↑WS & Uncategorised (WS) | 7.19E-03 | 5.20E-02 | -0.270 | 0.139 |
|  |  |  | ↓WS & Uncategorised (WS) | 3.70E-05 | 8.63E-01 | 0.012 | 0.072 |
|  |  |  | ↑W & Controls | 6.62E-04 | 1.44E-01 | -0.104 | 0.071 |
|  |  |  | ↓W & Controls | 1.91E-04 | 4.26E-01 | -0.044 | 0.056 |
|  |  |  | Uncategorised (W) & Controls | 1.90E-04 | 4.33E-01 | -0.050 | 0.063 |
|  |  |  | ↑W & ↓W | 1.95E-03 | 2.64E-01 | -0.093 | 0.084 |
|  |  |  | ↑W & Uncategorised (W) | 8.14E-04 | 5.40E-01 | -0.059 | 0.097 |
|  |  |  | ↓WS & Uncategorised (W) | 3.20E-04 | 6.46E-01 | 0.036 | 0.079 |
| ↓W & Controls | 0.1 | 20485 | ↑WS & Controls | 2.24E-06 | 9.33E-01 | 0.011 | 0.128 |
|  |  |  | ↓WS & Controls | 2.01E-03 | 1.00E-02 | 0.154 | 0.060 |
|  |  |  | Uncategorised (WS) & Controls | 2.96E-04 | 3.20E-01 | 0.049 | 0.049 |
|  |  |  | ↑WS & ↓WS | 4.14E-04 | 6.70E-01 | -0.060 | 0.140 |
|  |  |  | ↑WS & Uncategorised (WS) | 9.52E-07 | 9.82E-01 | -0.003 | 0.137 |
|  |  |  | ↓WS & Uncategorised (WS) | 2.30E-03 | 1.73E-01 | 0.097 | 0.071 |
|  |  |  | ↑W & Controls | 3.48E-04 | 2.89E-01 | 0.075 | 0.070 |
|  |  |  | ↓W & Controls | 1.74E-03 | 1.62E-02 | 0.133 | 0.055 |
|  |  |  | Uncategorised (W) & Controls | 1.42E-07 | 9.83E-01 | 0.001 | 0.063 |
|  |  |  | ↑W & ↓W | 7.12E-05 | 8.31E-01 | -0.018 | 0.082 |
|  |  |  | ↑W & Uncategorised (W) | 3.04E-03 | 2.36E-01 | 0.112 | 0.094 |
|  |  |  | ↓W & Uncategorised (W) | 4.11E-03 | 9.89E-02 | 0.133 | 0.081 |
| Uncategorised (W) & Controls | 0.5 | 64764 | ↑WS & Controls | 1.83E-04 | 4.47E-01 | 0.099 | 0.130 |
|  |  |  | ↓WS & Controls | 3.45E-04 | 2.86E-01 | 0.065 | 0.061 |
|  |  |  | **Uncategorised (WS) & Controls** | **3.54E-03** | **5.93E-04** | **0.169** | **0.049** |
|  |  |  | ↑WS & ↓WS | 5.06E-06 | 9.62E-01 | 0.007 | 0.146 |
|  |  |  | ↑WS & Uncategorised (WS) | 1.24E-03 | 4.20E-01 | -0.115 | 0.142 |
|  |  |  | ↓WS & Uncategorised (WS) | 6.40E-03 | 2.32E-02 | -0.166 | 0.073 |
|  |  |  | ↑W & Controls | 2.56E-04 | 3.63E-01 | 0.065 | 0.071 |
|  |  |  | ↓W & Controls | 6.64E-04 | 1.38E-01 | 0.084 | 0.056 |
|  |  |  | **Uncategorised (W) & Controls** | **3.09E-03** | **1.54E-03** | **0.201** | **0.063** |
|  |  |  | ↑W & ↓W | 3.00E-06 | 9.65E-01 | 0.004 | 0.085 |
|  |  |  | ↑W & Uncategorised (W) | 5.35E-03 | 1.16E-01 | -0.159 | 0.101 |
|  |  |  | ↓W & Uncategorised (W) | 1.97E-03 | 2.54E-01 | -0.094 | 0.083 |

**Supplementary Table 28**. Polygenic risk score results of subgroup PRS from UK Biobank case-control GWAS (unadjusted for BMI) tested for association in Generation Scotland samples

| UK Biobank PRS | | | Generation Scotland Sample | R-squared (Observed) | P-value | Coefficient | Standard Error |
| --- | --- | --- | --- | --- | --- | --- | --- |
| GWAS Summary Statistics | P-value Threshold | Number of SNPs |  |  |  |  |  |
| ↑WS & Controls | 0.001 | 378 | ↑WS & Controls | 1.49E-03 | 3.01E-02 | -0.280 | 0.129 |
|  |  |  | ↓WS & Controls | 3.47E-04 | 2.84E-01 | 0.065 | 0.060 |
|  |  |  | Uncategorised (WS) & Controls | 3.27E-04 | 2.97E-01 | 0.051 | 0.049 |
|  |  |  | ↑WS & ↓WS | 1.15E-02 | 2.44E-02 | -0.322 | 0.143 |
|  |  |  | ↑WS & Uncategorised (WS) | 9.73E-03 | 2.37E-02 | -0.335 | 0.148 |
|  |  |  | ↓WS & Uncategorised (WS) | 5.28E-05 | 8.37E-01 | 0.015 | 0.075 |
|  |  |  | ↑W & Controls | 1.81E-04 | 4.44E-01 | 0.054 | 0.071 |
|  |  |  | ↓W & Controls | 2.87E-04 | 3.29E-01 | 0.054 | 0.056 |
|  |  |  | Uncategorised (W) & Controls | 1.67E-04 | 4.62E-01 | 0.047 | 0.064 |
|  |  |  | ↑W & ↓W | 2.23E-04 | 7.06E-01 | -0.032 | 0.086 |
|  |  |  | ↑W & Uncategorised (W) | 2.40E-05 | 9.16E-01 | 0.011 | 0.102 |
|  |  |  | ↓W & Uncategorised (W) | 1.05E-05 | 9.34E-01 | -0.007 | 0.085 |
| ↓WS & Controls | 0.1 | 20241 | ↑WS & Controls | 5.74E-06 | 8.93E-01 | -0.017 | 0.128 |
|  |  |  | **↓WS & Controls** | **3.30E-03** | **9.64E-04** | **0.196** | **0.059** |
|  |  |  | Uncategorised (WS) & Controls | 4.02E-04 | 2.47E-01 | 0.057 | 0.049 |
|  |  |  | ↑WS & ↓WS | 1.95E-03 | 3.54E-01 | -0.129 | 0.139 |
|  |  |  | ↑WS & Uncategorised (WS) | 3.52E-04 | 6.68E-01 | -0.064 | 0.148 |
|  |  |  | ↓WS & Uncategorised (WS) | 4.94E-03 | 4.62E-02 | 0.145 | 0.073 |
|  |  |  | ↑W & Controls | 3.70E-04 | 2.74E-01 | 0.077 | 0.070 |
|  |  |  | **↓W & Controls** | **3.07E-03** | **1.40E-03** | **0.175** | **0.055** |
|  |  |  | Uncategorised (W) & Controls | 7.74E-06 | 8.74E-01 | -0.010 | 0.063 |
|  |  |  | ↑W & ↓W | 3.23E-03 | 1.51E-01 | -0.121 | 0.084 |
|  |  |  | ↑W & Uncategorised (W) | 1.71E-03 | 3.75E-01 | 0.089 | 0.101 |
|  |  |  | ↓W & Uncategorised (W) | 8.44E-03 | 1.80E-02 | 0.194 | 0.082 |
| Uncategorised (WS) & Controls | 0.1 | 20453 | ↑WS & Controls | 3.68E-05 | 7.33E-01 | -0.044 | 0.128 |
|  |  |  | ↓WS & Controls | 2.26E-04 | 3.87E-01 | 0.052 | 0.060 |
|  |  |  | **Uncategorised (WS) & Controls** | **3.19E-03** | **1.11E-03** | **0.159** | **0.049** |
|  |  |  | ↑WS & ↓WS | 3.28E-03 | 2.30E-01 | -0.165 | 0.137 |
|  |  |  | ↑WS & Uncategorised (WS) | 5.57E-03 | 8.73E-02 | -0.245 | 0.143 |
|  |  |  | ↓WS & Uncategorised (WS) | 4.70E-03 | 5.18E-02 | -0.142 | 0.073 |
|  |  |  | ↑W & Controls | 3.28E-04 | 3.03E-01 | 0.073 | 0.071 |
|  |  |  | ↓W & Controls | 5.16E-04 | 1.91E-01 | 0.073 | 0.055 |
|  |  |  | Uncategorised (W) & Controls | 1.70E-03 | 1.89E-02 | 0.147 | 0.063 |
|  |  |  | ↑W & ↓W | 4.36E-07 | 9.87E-01 | -0.001 | 0.084 |
|  |  |  | ↑W & Uncategorised (W) | 3.57E-03 | 1.99E-01 | -0.129 | 0.100 |
|  |  |  | ↓W & Uncategorised (W) | 7.76E-04 | 4.74E-01 | -0.057 | 0.080 |
| ↑W & Controls | 0.01 | 2940 | ↑WS & Controls | 1.22E-04 | 5.35E-01 | -0.081 | 0.130 |
|  |  |  | ↓WS & Controls | 1.12E-03 | 5.46E-02 | 0.117 | 0.061 |
|  |  |  | Uncategorised (WS) & Controls | 1.84E-03 | 1.33E-02 | 0.122 | 0.049 |
|  |  |  | ↑WS & ↓WS | 2.15E-03 | 3.31E-01 | -0.139 | 0.143 |
|  |  |  | ↑WS & Uncategorised (WS) | 3.18E-03 | 1.97E-01 | -0.181 | 0.140 |
|  |  |  | ↓WS & Uncategorised (WS) | 4.23E-05 | 8.54E-01 | 0.013 | 0.073 |
|  |  |  | ↑W & Controls | 1.46E-03 | 2.99E-02 | 0.155 | 0.072 |
|  |  |  | ↓W & Controls | 5.85E-04 | 1.64E-01 | 0.078 | 0.056 |
|  |  |  | Uncategorised (W) & Controls | 1.09E-03 | 6.01E-02 | 0.120 | 0.064 |
|  |  |  | ↑W & ↓W | 1.29E-03 | 3.65E-01 | 0.076 | 0.084 |
|  |  |  | ↑W & Uncategorised (W) | 7.24E-05 | 8.55E-01 | 0.018 | 0.099 |
|  |  |  | ↓WS & Uncategorised (W) | 1.07E-04 | 7.90E-01 | -0.022 | 0.081 |
| ↓W & Controls | 1 | 93093 | ↑WS & Controls | 1.81E-04 | 4.50E-01 | -0.097 | 0.128 |
|  |  |  | ↓WS & Controls | 1.94E-03 | 1.14E-02 | 0.151 | 0.060 |
|  |  |  | Uncategorised (WS) & Controls | 7.07E-04 | 1.25E-01 | 0.075 | 0.049 |
|  |  |  | ↑WS & ↓WS | 4.91E-03 | 1.42E-01 | -0.211 | 0.144 |
|  |  |  | ↑WS & Uncategorised (WS) | 3.37E-03 | 1.84E-01 | -0.184 | 0.138 |
|  |  |  | ↓WS & Uncategorised (WS) | 1.23E-03 | 3.20E-01 | 0.071 | 0.072 |
|  |  |  | ↑W & Controls | 1.13E-04 | 5.45E-01 | 0.042 | 0.070 |
|  |  |  | ↓W & Controls | 1.76E-03 | 1.56E-02 | 0.134 | 0.055 |
|  |  |  | Uncategorised (W) & Controls | 1.80E-04 | 4.45E-01 | 0.048 | 0.063 |
|  |  |  | ↑W & ↓W | 1.21E-03 | 3.80E-01 | -0.073 | 0.083 |
|  |  |  | ↑W & Uncategorised (W) | 3.76E-05 | 8.95E-01 | 0.013 | 0.095 |
|  |  |  | ↓W & Uncategorised (W) | 1.29E-03 | 3.56E-01 | 0.076 | 0.082 |
| Uncategorised (W) & Controls | 0.1 | 20267 | ↑WS & Controls | 2.95E-04 | 3.35E-01 | 0.124 | 0.129 |
|  |  |  | ↓WS & Controls | 9.53E-05 | 5.75E-01 | 0.034 | 0.060 |
|  |  |  | **Uncategorised (WS) & Controls** | **4.49E-03** | **1.09E-04** | **0.189** | **0.049** |
|  |  |  | ↑WS & ↓WS | 9.55E-04 | 5.18E-01 | 0.090 | 0.139 |
|  |  |  | ↑WS & Uncategorised (WS) | 5.79E-04 | 5.82E-01 | -0.080 | 0.145 |
|  |  |  | **↓WS & Uncategorised (WS)** | **1.09E-02** | **2.96E-03** | **-0.215** | **0.072** |
|  |  |  | ↑W & Controls | 1.12E-03 | 5.67E-02 | 0.135 | 0.071 |
|  |  |  | ↓W & Controls | 3.23E-04 | 3.01E-01 | 0.058 | 0.056 |
|  |  |  | **Uncategorised (W) & Controls** | **3.33E-03** | **9.98E-04** | **0.207** | **0.063** |
|  |  |  | ↑W & ↓W | 1.86E-03 | 2.76E-01 | 0.092 | 0.084 |
|  |  |  | ↑W & Uncategorised (W) | 2.25E-03 | 3.09E-01 | -0.104 | 0.102 |
|  |  |  | ↓W & Uncategorised (W) | 3.86E-03 | 1.10E-01 | -0.129 | 0.081 |

**Supplementary Table 29**. Polygenic risk score results of subgroup PRS from UK Biobank case-only GWAS (adjusted for BMI) tested for association in Generation Scotland samples

| UK Biobank PRS | | | Generation Scotland Sample | R-squared (Observed) | P-value | Coefficient | Standard Error |
| --- | --- | --- | --- | --- | --- | --- | --- |
| GWAS Summary Statistics | P-value Threshold | Number of SNPs |  |  |  |  |  |
| ↑WS & ↓WS | 0.001 | 395 | ↑WS & Controls | 2.30E-03 | 6.99E-03 | 0.347 | 0.129 |
|  |  |  | ↓WS & Controls | 1.66E-04 | 4.59E-01 | 0.045 | 0.060 |
|  |  |  | Uncategorised (WS) & Controls | 5.64E-07 | 9.65E-01 | 0.002 | 0.049 |
|  |  |  | ↑WS & ↓WS | 7.02E-03 | 7.89E-02 | 0.261 | 0.148 |
|  |  |  | ↑WS & Uncategorised (WS) | 9.89E-03 | 2.25E-02 | 0.316 | 0.138 |
|  |  |  | ↓WS & Uncategorised (WS) | 6.78E-04 | 4.60E-01 | 0.055 | 0.074 |
|  |  |  | ↑W & Controls | 8.44E-04 | 9.85E-02 | 0.116 | 0.070 |
|  |  |  | ↓W & Controls | 6.64E-05 | 6.39E-01 | 0.026 | 0.056 |
|  |  |  | Uncategorised (W) & Controls | 2.71E-06 | 9.25E-01 | -0.006 | 0.063 |
|  |  |  | ↑W & ↓W | 3.59E-03 | 1.30E-01 | 0.130 | 0.085 |
|  |  |  | ↑W & Uncategorised (W) | 1.06E-02 | 2.65E-02 | 0.215 | 0.097 |
|  |  |  | ↓W & Uncategorised (W) | 3.22E-04 | 6.45E-01 | 0.039 | 0.084 |
| ↑WS & Uncategorised (WS) | 5 x 10^-4^ | 179 | ↑WS & Controls | 1.91E-04 | 4.37E-01 | -0.099 | 0.128 |
|  |  |  | ↓WS & Controls | 3.63E-04 | 2.74E-01 | -0.065 | 0.059 |
|  |  |  | Uncategorised (WS) & Controls | 1.92E-04 | 4.24E-01 | 0.039 | 0.049 |
|  |  |  | ↑WS & ↓WS | 2.95E-04 | 7.19E-01 | -0.053 | 0.147 |
|  |  |  | ↑WS & Uncategorised (WS) | 2.63E-03 | 2.40E-01 | -0.172 | 0.146 |
|  |  |  | ↓WS & Uncategorised (WS) | 2.74E-03 | 1.37E-01 | -0.110 | 0.074 |
|  |  |  | ↑W & Controls | 7.45E-05 | 6.24E-01 | 0.034 | 0.070 |
|  |  |  | ↓W & Controls | 1.37E-04 | 5.01E-01 | -0.037 | 0.055 |
|  |  |  | Uncategorised (W) & Controls | 1.67E-04 | 4.61E-01 | 0.046 | 0.063 |
|  |  |  | ↑W & ↓W | 2.11E-03 | 2.45E-01 | 0.102 | 0.088 |
|  |  |  | ↑W & Uncategorised (W) | 1.06E-04 | 8.25E-01 | -0.023 | 0.103 |
|  |  |  | ↓W & Uncategorised (W) | 3.23E-03 | 1.44E-01 | -0.121 | 0.083 |
| ↓WS & Uncategorised (WS) | 0.1 | 19510 | ↑WS & Controls | 3.68E-04 | 2.81E-01 | -0.141 | 0.131 |
|  |  |  | ↓WS & Controls | 1.44E-03 | 2.90E-02 | 0.134 | 0.061 |
|  |  |  | Uncategorised (WS) & Controls | 6.11E-05 | 6.52E-01 | -0.022 | 0.050 |
|  |  |  | ↑WS & ↓WS | 4.03E-03 | 1.83E-01 | -0.182 | 0.137 |
|  |  |  | ↑WS & Uncategorised (WS) | 5.14E-04 | 6.04E-01 | -0.072 | 0.139 |
|  |  |  | ↓WS & Uncategorised (WS) | 7.59E-03 | 1.33E-02 | 0.181 | 0.073 |
|  |  |  | ↑W & Controls | 7.47E-06 | 8.77E-01 | -0.011 | 0.071 |
|  |  |  | ↓W & Controls | 1.03E-03 | 6.47E-02 | 0.105 | 0.057 |
|  |  |  | Uncategorised (W) & Controls | 2.58E-04 | 3.60E-01 | -0.059 | 0.064 |
|  |  |  | ↑W & ↓W | 2.31E-03 | 2.24E-01 | -0.100 | 0.082 |
|  |  |  | ↑W & Uncategorised (W) | 2.36E-03 | 2.97E-01 | 0.099 | 0.095 |
|  |  |  | ↓W & Uncategorised (W) | 6.09E-03 | 4.45E-02 | 0.166 | 0.083 |
| ↑W & ↓W | 0.1 | 19561 | ↑WS & Controls | 3.19E-04 | 3.16E-01 | 0.132 | 0.131 |
|  |  |  | ↓WS & Controls | 5.81E-04 | 1.66E-01 | -0.085 | 0.062 |
|  |  |  | Uncategorised (WS) & Controls | 3.35E-06 | 9.16E-01 | -0.005 | 0.050 |
|  |  |  | ↑WS & ↓WS | 3.71E-03 | 2.02E-01 | 0.186 | 0.146 |
|  |  |  | ↑WS & Uncategorised (WS) | 6.21E-04 | 5.68E-01 | 0.079 | 0.138 |
|  |  |  | ↓WS & Uncategorised (WS) | 7.41E-04 | 4.40E-01 | -0.056 | 0.072 |
|  |  |  | ↑W & Controls | 1.51E-05 | 8.25E-01 | 0.016 | 0.071 |
|  |  |  | ↓W & Controls | 6.46E-04 | 1.43E-01 | -0.084 | 0.057 |
|  |  |  | Uncategorised (W) & Controls | 6.62E-06 | 8.83E-01 | 0.009 | 0.065 |
|  |  |  | ↑W & ↓W | 2.03E-03 | 2.55E-01 | 0.096 | 0.084 |
|  |  |  | ↑W & Uncategorised (W) | 3.33E-05 | 9.01E-01 | 0.012 | 0.094 |
|  |  |  | ↓WS & Uncategorised (W) | 8.85E-04 | 4.45E-01 | -0.064 | 0.084 |
| ↑W & Uncategorised (W) | 1 | 93110 | ↑WS & Controls | 2.30E-04 | 3.94E-01 | 0.110 | 0.129 |
|  |  |  | ↓WS & Controls | 2.63E-04 | 3.52E-01 | -0.057 | 0.061 |
|  |  |  | Uncategorised (WS) & Controls | 6.72E-05 | 6.36E-01 | -0.023 | 0.049 |
|  |  |  | ↑WS & ↓WS | 2.24E-03 | 3.21E-01 | 0.145 | 0.146 |
|  |  |  | ↑WS & Uncategorised (WS) | 5.84E-04 | 5.80E-01 | 0.078 | 0.140 |
|  |  |  | ↓WS & Uncategorised (WS) | 4.01E-05 | 8.58E-01 | 0.013 | 0.073 |
|  |  |  | ↑W & Controls | 1.54E-04 | 4.80E-01 | -0.049 | 0.070 |
|  |  |  | ↓W & Controls | 3.68E-04 | 2.69E-01 | -0.063 | 0.057 |
|  |  |  | Uncategorised (W) & Controls | 7.04E-05 | 6.33E-01 | 0.030 | 0.064 |
|  |  |  | ↑W & ↓W | 4.28E-03 | 9.82E-02 | 0.140 | 0.085 |
|  |  |  | ↑W & Uncategorised (W) | 4.21E-03 | 1.63E-01 | 0.134 | 0.096 |
|  |  |  | ↓W & Uncategorised (W) | 8.76E-06 | 9.39E-01 | 0.006 | 0.084 |
| ↓W & Uncategorised (W) | 0.1 | 19535 | ↑WS & Controls | 8.15E-04 | 1.09E-01 | -0.207 | 0.129 |
|  |  |  | ↓WS & Controls | 5.91E-04 | 1.63E-01 | 0.084 | 0.060 |
|  |  |  | Uncategorised (WS) & Controls | 2.18E-05 | 7.87E-01 | -0.013 | 0.049 |
|  |  |  | ↑WS & ↓WS | 5.66E-03 | 1.15E-01 | -0.215 | 0.136 |
|  |  |  | ↑WS & Uncategorised (WS) | 2.40E-03 | 2.62E-01 | -0.158 | 0.141 |
|  |  |  | ↓WS & Uncategorised (WS) | 3.11E-03 | 1.14E-01 | 0.112 | 0.071 |
|  |  |  | ↑W & Controls | 2.28E-05 | 7.86E-01 | 0.019 | 0.070 |
|  |  |  | ↓W & Controls | 3.76E-04 | 2.64E-01 | 0.062 | 0.055 |
|  |  |  | Uncategorised (W) & Controls | 4.98E-04 | 2.04E-01 | -0.081 | 0.063 |
|  |  |  | ↑W & ↓W | 4.56E-05 | 8.65E-01 | -0.014 | 0.080 |
|  |  |  | ↑W & Uncategorised (W) | 3.14E-03 | 2.29E-01 | 0.118 | 0.098 |
|  |  |  | ↓W & Uncategorised (W) | 3.13E-03 | 1.50E-01 | 0.117 | 0.081 |

**Supplementary Table 30**. Polygenic risk score results of subgroup PRS from UK Biobank case-only GWAS (unadjusted for BMI) tested for association in Generation Scotland samples

| UK Biobank PRS | | | Generation Scotland Sample | R-squared (Observed) | P-value | Coefficient | Standard Error |
| --- | --- | --- | --- | --- | --- | --- | --- |
| GWAS Summary Statistics | P-value Threshold | Number of SNPs |  |  |  |  |  |
| ↑WS & ↓WS | 1 |  | ↑WS & Controls | 1.20E-03 | 5.15E-02 | 0.250 | 0.128 |
|  |  |  | ↓WS & Controls | 1.80E-05 | 8.08E-01 | -0.015 | 0.060 |
|  |  |  | Uncategorised (WS) & Controls | 5.14E-05 | 6.79E-01 | -0.020 | 0.049 |
|  |  |  | ↑WS & ↓WS | 4.82E-03 | 1.12E-01 | 0.221 | 0.139 |
|  |  |  | ↑WS & Uncategorised (WS) | 2.73E-04 | 6.40E-01 | -0.034 | 0.073 |
|  |  |  | ↓WS & Uncategorised (WS) | 7.67E-04 | 1.15E-01 | 0.110 | 0.070 |
|  |  |  | ↑W & Controls | 1.99E-04 | 4.17E-01 | -0.045 | 0.055 |
|  |  |  | ↓W & Controls | 6.12E-06 | 8.88E-01 | -0.009 | 0.063 |
|  |  |  | Uncategorised (W) & Controls | 7.12E-03 | 3.28E-02 | 0.173 | 0.081 |
|  |  |  | ↑W & ↓W | 7.41E-03 | 6.41E-02 | 0.177 | 0.095 |
|  |  |  | ↑W & Uncategorised (W) | 2.43E-04 | 6.89E-01 | -0.034 | 0.084 |
|  |  |  | ↓W & Uncategorised (W) | 5.92E-03 | 1.07E-01 | 0.229 | 0.142 |
| ↑WS & Uncategorised (WS) | 1 x 10^-5^ |  | ↑WS & Controls | 1.70E-03 | 2.05E-02 | 0.303 | 0.131 |
|  |  |  | ↓WS & Controls | 5.32E-04 | 1.85E-01 | 0.080 | 0.060 |
|  |  |  | Uncategorised (WS) & Controls | 2.14E-05 | 7.89E-01 | -0.013 | 0.050 |
|  |  |  | ↑WS & ↓WS | 1.31E-02 | 8.56E-03 | 0.360 | 0.136 |
|  |  |  | ↑WS & Uncategorised (WS) | 3.29E-03 | 1.04E-01 | 0.116 | 0.071 |
|  |  |  | ↓WS & Uncategorised (WS) | 1.52E-04 | 4.84E-01 | 0.050 | 0.072 |
|  |  |  | ↑W & Controls | 7.28E-04 | 1.20E-01 | 0.087 | 0.056 |
|  |  |  | ↓W & Controls | 9.03E-05 | 5.88E-01 | 0.035 | 0.064 |
|  |  |  | Uncategorised (W) & Controls | 4.78E-05 | 8.61E-01 | -0.014 | 0.081 |
|  |  |  | ↑W & ↓W | 1.61E-03 | 3.89E-01 | 0.082 | 0.095 |
|  |  |  | ↑W & Uncategorised (W) | 3.88E-04 | 6.13E-01 | 0.040 | 0.080 |
|  |  |  | ↓W & Uncategorised (W) | 8.10E-03 | 5.90E-02 | 0.256 | 0.135 |
| ↓WS & Uncategorised (WS) | 0.1 |  | ↑WS & Controls | 2.01E-04 | 4.25E-01 | -0.104 | 0.130 |
|  |  |  | ↓WS & Controls | 1.59E-03 | 2.21E-02 | 0.140 | 0.061 |
|  |  |  | Uncategorised (WS) & Controls | 3.08E-04 | 3.11E-01 | -0.050 | 0.050 |
|  |  |  | ↑WS & ↓WS | 1.42E-04 | 7.85E-01 | -0.038 | 0.138 |
|  |  |  | ↑WS & Uncategorised (WS) | 9.04E-03 | 6.90E-03 | 0.199 | 0.073 |
|  |  |  | ↓WS & Uncategorised (WS) | 1.15E-06 | 9.51E-01 | -0.004 | 0.071 |
|  |  |  | ↑W & Controls | 1.19E-03 | 4.68E-02 | 0.112 | 0.057 |
|  |  |  | ↓W & Controls | 1.04E-03 | 6.57E-02 | -0.118 | 0.064 |
|  |  |  | Uncategorised (W) & Controls | 2.86E-03 | 1.77E-01 | -0.113 | 0.084 |
|  |  |  | ↑W & ↓W | 5.49E-03 | 1.11E-01 | 0.151 | 0.095 |
|  |  |  | ↑W & Uncategorised (W) | 1.01E-02 | 9.63E-03 | 0.213 | 0.082 |
|  |  |  | ↓W & Uncategorised (W) | 3.73E-03 | 2.01E-01 | -0.180 | 0.140 |
| ↑W & ↓W | 0.5 |  | ↑WS & Controls | 2.14E-04 | 4.11E-01 | 0.108 | 0.131 |
|  |  |  | ↓WS & Controls | 5.46E-04 | 1.79E-01 | -0.083 | 0.061 |
|  |  |  | Uncategorised (WS) & Controls | 4.49E-05 | 6.99E-01 | 0.019 | 0.050 |
|  |  |  | ↑WS & ↓WS | 2.59E-05 | 9.07E-01 | 0.016 | 0.141 |
|  |  |  | ↑WS & Uncategorised (WS) | 2.45E-03 | 1.60E-01 | -0.105 | 0.075 |
|  |  |  | ↓WS & Uncategorised (WS) | 3.84E-04 | 2.66E-01 | 0.080 | 0.071 |
|  |  |  | ↑W & Controls | 1.02E-03 | 6.55E-02 | -0.105 | 0.057 |
|  |  |  | ↓W & Controls | 1.94E-04 | 4.28E-01 | 0.051 | 0.065 |
|  |  |  | Uncategorised (W) & Controls | 7.25E-03 | 3.13E-02 | 0.184 | 0.085 |
|  |  |  | ↑W & ↓W | 1.89E-04 | 7.68E-01 | 0.029 | 0.098 |
|  |  |  | ↑W & Uncategorised (W) | 2.96E-03 | 1.62E-01 | -0.120 | 0.086 |
|  |  |  | ↓WS & Uncategorised (W) | 2.53E-03 | 2.92E-01 | 0.159 | 0.150 |
| ↑W & Uncategorised (W) | 0.01 |  | ↑WS & Controls | 1.23E-03 | 4.87E-02 | -0.260 | 0.132 |
|  |  |  | ↓WS & Controls | 2.71E-04 | 3.44E-01 | -0.058 | 0.061 |
|  |  |  | Uncategorised (WS) & Controls | 2.37E-06 | 9.29E-01 | -0.004 | 0.050 |
|  |  |  | ↑WS & ↓WS | 8.11E-03 | 3.90E-02 | -0.307 | 0.148 |
|  |  |  | ↑WS & Uncategorised (WS) | 4.09E-04 | 5.67E-01 | -0.043 | 0.075 |
|  |  |  | ↓WS & Uncategorised (WS) | 8.94E-05 | 5.91E-01 | -0.039 | 0.072 |
|  |  |  | ↑W & Controls | 7.32E-04 | 1.19E-01 | -0.088 | 0.057 |
|  |  |  | ↓W & Controls | 2.53E-05 | 7.75E-01 | 0.019 | 0.065 |
|  |  |  | Uncategorised (W) & Controls | 3.48E-04 | 6.37E-01 | 0.040 | 0.085 |
|  |  |  | ↑W & ↓W | 3.51E-03 | 2.03E-01 | -0.134 | 0.105 |
|  |  |  | ↑W & Uncategorised (W) | 4.01E-03 | 1.03E-01 | -0.140 | 0.086 |
|  |  |  | ↓W & Uncategorised (W) | 3.12E-03 | 2.42E-01 | -0.169 | 0.144 |
| ↓W & Uncategorised (W) | 0.1 |  | ↑WS & Controls | 3.84E-04 | 2.71E-01 | -0.142 | 0.129 |
|  |  |  | ↓WS & Controls | 5.27E-04 | 1.87E-01 | 0.079 | 0.060 |
|  |  |  | Uncategorised (WS) & Controls | 3.14E-05 | 7.47E-01 | -0.016 | 0.049 |
|  |  |  | ↑WS & ↓WS | 1.10E-03 | 4.47E-01 | -0.108 | 0.142 |
|  |  |  | ↑WS & Uncategorised (WS) | 3.02E-03 | 1.19E-01 | 0.112 | 0.072 |
|  |  |  | ↓WS & Uncategorised (WS) | 1.08E-04 | 5.54E-01 | 0.042 | 0.071 |
|  |  |  | ↑W & Controls | 2.72E-04 | 3.42E-01 | 0.052 | 0.055 |
|  |  |  | ↓W & Controls | 6.50E-04 | 1.46E-01 | -0.092 | 0.063 |
|  |  |  | Uncategorised (W) & Controls | 1.82E-04 | 7.33E-01 | 0.028 | 0.082 |
|  |  |  | ↑W & ↓W | 5.99E-03 | 9.63E-02 | 0.164 | 0.099 |
|  |  |  | ↑W & Uncategorised (W) | 3.22E-03 | 1.44E-01 | 0.118 | 0.081 |
|  |  |  | ↓W & Uncategorised (W) | 3.13E-03 | 2.41E-01 | -0.163 | 0.139 |

**Supplementary Table 31**. Genetic correlations between depression subgroups and between subgroups and MDD (Wray *et al*., 2018)

|  | **Adjusted for BMI** | | | **Unadjusted for BMI** | | |
| --- | --- | --- | --- | --- | --- | --- |
|  | ***rg (SE)*** | ***Z-score*** | ***p*** | ***rg (SE)*** | ***Z-score*** | ***p*** |
| ***Between subgroups*** | | | | | | |
| ↑ Weight & Sleep +  ↓ Weight & Sleep | 0.5564  (0.2202) | 2.5265 | 0.0115 | 0.7198  (0.2537) | 2.8373 | 4.60E-03 |
| ↑ Weight & Sleep +  Uncategorised (WS) | 0.5949  (0.1906) | 3.1219 | 0.0018 | 0.8785  (0.2598) | 3.3815 | 7.00E-04 |
| ↓ Weight & Sleep +  Uncategorised (WS) | 0.9762  (0.1164) | 8.3894 | 4.89E-17 | 0.9456  (0.1149) | 8.2275 | 1.91E-16 |
| ↑ Weight +  ↓ Weight | 0.8686  (0.2423) | 3.5845 | 0.0003 | 0.8466  (0.1794) | 4.7187 | 2.37E-06 |
| ↑ Weight +  Uncategorised (W)eight | 0.904  (0.2243) | 4.0313 | 5.55E-05 | 0.8482  (0.1662) | 5.1043 | 3.32E-07 |
| ↓ Weight +  Uncategorised (W)eight | 0.8235  (0.1176) | 7.0002 | 2.56E-12 | 0.8122  (0.1176) | 6.9067 | 4.96E-12 |
| ***Between weight-sleep and weight-only subgroups*** | | | | | | |
| ↑ Weight & Sleep +  ↑ Weight | 0.8557  (0.1796) | 4.7649 | 1.89E-06 | 0.8667  (0.1779) | 4.8717 | 1.11E-06 |
| ↓ Weight & Sleep +  ↓ Weight | 0.9995  (0.0156) | 64.2464 | 0 | 1.0001  (0.0154) | 65.0777 | 0.00E+00 |
| Uncategorised (WS) +  Uncategorised (W)eight | 0.9792  (0.0161) | 60.7191 | 0 | 0.968  (0.0181) | 53.4496 | 0.00E+00 |
| ***Between subgroups and MDD (Wray et al., 2018)*** | | | | | | |
| ↑ Weight & Sleep +  MDD | 0.4046*  (0.153) | 2.6453 | 0.0082 | 0.5346  (0.1912) | 2.7969 | 5.20E-03 |
| ↓ Weight & Sleep + MDD | 0.7435  (0.1064) | 6.9899 | 2.75E-12 | 0.7325  (0.1051) | 6.9703 | 3.16E-12 |
| Uncategorised (WS) +  MDD | 0.9109*  (0.0943) | 9.6632 | 4.32E-22 | 0.93  (0.0925) | 10.0523 | 8.98E-24 |
| ↑ Weight +  MDD | 0.8659  (0.1791) | 4.8336 | 1.34E-06 | 0.8294  (0.1284) | 6.4572 | 1.07E-10 |
| ↓ Weight +  MDD | 0.7794  (0.0985) | 7.9153 | 2.47E-15 | 0.7674  (0.0979) | 7.8386 | 4.56E-15 |
| Uncategorised (W)eight +  MDD | 0.8051  (0.096) | 8.3876 | 4.96E-17 | 0.8184  (0.0949) | 8.6246 | 6.43E-18 |

**Supplementary Table 32.** The genetic correlation between summary statistics adjusted for BMI and summary statistics unadjusted for BMI for each subgroup

|  | **rg** | **se** | **Z score** | **P value** |
| --- | --- | --- | --- | --- |
| ↑ Weight & Sleep | 0.936 | 0.0874 | 10.70 | 1.01E-26 |
| ↓ Weight & Sleep | 0.999* | 0.0003 | 3967.69 | 0.00 |
| Uncategorised (WS) | 0.983* | 0.0031 | 317.81 | 0.00 |
| ↑ Weight | 0.970 | 0.0462 | 21.00 | 5.49E-98 |
| ↓ Weight | 0.999† | 0.0003 | 3161.35 | 0.00 |
| Uncategorised (W) | 0.995† | 0.0012 | 843.01 | 0.00 |

Significantly different comparison pairs are highlighted with matching symbols (*, †)

**Supplementary Table 33.** GWAS Results for loci in *ALK*, *EPHB1*, *CLSTN2* and *LSAMP* comparing methods of BMI adjustment

| **SNP**  **(Gene)** | **BMI Adjustment** | **↑WS vs Controls** | | **↑W vs Controls** | | **↑WS vs ↓WS** | | **↑W vs ↓W** | |
| --- | --- | --- | --- | --- | --- | --- | --- | --- | --- |
|  |  | ***Effect Size***  ***(95% CI)*** | ***P-value*** | ***Effect Size***  ***(95% CI)*** | ***P-value*** | ***Effect Size***  ***(95% CI)*** | ***P-value*** | ***Effect Size*** | ***P-value*** |
| rs7608034  (*ALK*) | Unadjusted | 0.606  (0.291-0.920) | 1.61E-04 | 0.160  (0.006-0.314) | 4.13E-02 | 0.238  (-0.008-0.484) | 5.81E-02 | 0.022  (-0.118-0.162) | 7.60E-01 |
|  | Predjusted | **1.140**  **(0.736-1.544)** | **3.09E-08** | 0.251  (0.076-0.426) | 5.03E-03 | 0.571  (0.275-0.868) | 1.61E-04 | 0.103  (-0.069-0.274) | 2.40E-01 |
|  | Covariate | 0.597  (0.284-0.910) | 1.84E-04 | 0.151  (0.000-0.302) | 5.01E-02 | 0.200  (-0.034-0.434) | 9.45E-02 | -0.009  (-0.139-0.121) | 8.93E-01 |
| rs1554675  (*EPHB1*) | Unadjusted | 0.334  (0.105-0.563) | 4.24E-03 | -0.001  (-0.114-0.111) | 9.81E-01 | 0.089  (-0.093-0.272) | 3.36E-01 | -0.055  (-0.159-0.050) | 3.05E-01 |
|  | Predjusted | **0.918**  **(0.625-1.211)** | **8.29E-10** | 0.074  (-0.054-0.202) | 2.56E-01 | 0.345  (0.125-0.564) | 2.08E-03 | -0.064  (-0.192-0.064) | 3.31E-01 |
|  | Covariate | 0.344  (0.116-0.572) | 3.04E-03 | 0.003  (-0.107-0.114) | 9.53E-01 | 0.134  (-0.039-0.308) | 1.29E-01 | -0.030  (-0.127-0.067) | 5.48E-01 |
| rs1426041  (*CLSTN2*) | Unadjusted | 0.413  (0.255-0.571) | 3.07E-07 | 0.114  (0.037-0.191) | 3.84E-03 | 0.286  (0.160-0.412) | 8.04E-06 | 0.066  (-0.006-0.137) | 7.16E-02 |
|  | Predjusted | **0.683**  **(0.479-0.887)** | **5.12E-11** | 0.161  (0.073-0.249) | 3.45E-04 | 0.415  (0.264-0.567) | 8.33E-08 | 0.096  (0.010-0.183) | 2.94E-02 |
|  | Covariate | 0.430  (0.273-0.587) | 8.19E-08 | 0.129  (0.053-0.205) | 8.94E-04 | 0.267  (0.147-0.386) | 1.25E-05 | 0.065  (-0.001-0.131) | 5.51E-02 |
| rs4688064  (*LSAMP*) | Unadjusted | 0.108  (-0.009-0.225) | 6.96E-02 | 0.122  (0.065-0.179) | 2.89E-05 | 0.182  (0.087-0.277) | 1.83E-04 | 0.149  (0.095-0.202) | 5.36E-08 |
|  | Predjusted | 0.088  (-0.062-0.238) | 2.52E-01 | 0.120  (0.055-0.186) | 3.03E-04 | 0.191  (0.076-0.306) | 1.17E-03 | 0.163  (0.097-0.229) | 1.22E-06 |
|  | Covariate | 0.094  (-0.022-0.210) | 1.13E-01 | 0.109  (0.053-0.166) | 1.38E-04 | 0.156  (0.066-0.247) | 7.22E-04 | 0.134  (0.084-0.183) | 1.37E-07 |

*Results of genome-wide significance (5e-08) are highlighted in* ***bold****. Results of suggestive significance are in underlined. 95% CI = 95% Confidence Interval.*

**Supplementary Table 34.** Association of loci in *ALK*, *EPHB1*, *CLSTN2* and *LSAMP* with loci previously associated with BMI

| **SNP**  **(Gene)** | **LD region** | **SNP** | **P-value** | **Study**  **(Phenotype)** | **d'** | **r^2^** | **P-value** |
| --- | --- | --- | --- | --- | --- | --- | --- |
| rs7608034  (*ALK*) | Chr2 29-31 Mb | rs7578465 | 8.00E-06 | Locke *et al*., 2015  (BMI) | 0.202 | 0.0005 | 0.7654 |
| rs1554675  (*EPHB1*) | Chr3 134.5-135.5 Mb | rs40157 | 3.00E-08 | Sakaue *et al*., 2021  (Weight) | 0.1911 | 0.0021 | 0.5386 |
| rs1426041  (*CLSTN2*) | Chr3 139.5-140 Mb | rs2554152 | 7.00E-06 | Fox *et al*., 2012  (Visceral adipose tissue adjusted for BMI) | 0.1862 | 0.0284 | 0.0231 |
| rs4688064  (*LSAMP*) | Chr3 117-118.5 Mb | **rs768917** | **1.00E-08** | **Zhu *et al.,* 2019**  **(BMI)** | **0.8761** | **0.586** | **<0.0001** |
|  |  |  | **3.00E-08** | **Kichaev *et al*., 2018**  **(BMI)** |  |  |  |
|  |  | rs779206 | 3.00E-10 | Pulit *et al*., 2018  (BMI) | 0.0836 | 0.0063 | 0.2859 |
|  |  | rs558882 | 1.00E-08 | Zhu et al., 2019  (BMI) | 0.037 | 0.0012 | 0.6428 |

*Loci in linkage disequilibrium are highlighted in* ***bold****.*
